# Viral Nuclease Inhibitors: Small molecule disruptors of the UL12 alkaline nuclease display broad anti-herpes virus activity

**DOI:** 10.64898/2026.06.03.729350

**Authors:** Nidhi Sharma, Xiaoyun Xie, Renata Szczepaniak, Chitra Rani, Sheida Khojasteh Khosro, Jolanta Krucinska, Xiaoling Chen, Dung Do, Lee Wright, Dennis Wright, Sandra Weller

## Abstract

Herpes simplex virus-1 (HSV-1) UL12 gene encodes a well-conserved 5’ → 3’ alkaline exonuclease. UL12 collaborates with the HSV single-strand DNA binding protein ICP8 to mediate recombination-dependent replication of viral DNA and is essential for the production of DNA that can be packaged into infectious virus. The UL12 gene has orthologs in the eight other human herpesviruses, including UL98 in HCMV and SOX in KSHV, which are also essential for virus production. We have developed viral nuclease inhibitors (VNIs) of HSV-1 UL12 that potently block its nuclease activity and display strong antiviral effects in cell culture. These inhibitors are also effective against alkaline nucleases from the β-HHV HCMV (UL98) and the γ-HHV KSHV (SOX), and we have demonstrated antiviral activity against HSV-1 and HCMV in cell culture. In this work, we describe the first crystal structure of an alphaherpesvirus alkaline nuclease (UL12.5), which was used to elucidate structure activity relationships and improve the selectivity of our inhibitors. These VNIs exhibit EC_50_ and IC_50_ values in the nanomolar to low micromolar range. Our findings highlight the potential of targeting HHV alkaline nucleases with novel small molecules, paving the way for the development of new therapies that can be broadly antiviral on their own or in combination with nucleoside analogs.

**Significance Statement:** Herpesviruses are widespread pathogens that establish lifelong infections and cause serious disease in immunocompromised individuals, neonates and older adults, yet treatment options remain limited. We report the first crystal structure of the HSV-1 alkaline nuclease UL12.5 and use it to design potent small-molecule inhibitors. These inhibitors exhibit antiviral activity against HSV-1 and HCMV, supporting alkaline nuclease as a conserved and druggable target across all herpesvirus subfamilies. The significance is threefold: it confirms alkaline nucleases as having an essential role in viral replication, provides a structural foundation for rational antiviral design and introduces a new class of inhibitors with potential as pan-herpesvirus therapeutics, alone or in combination, to overcome resistance and improve clinical outcomes.

## Introduction

Human herpesviruses (HHVs) are large DNA viruses with a profound global clinical impact. They establish lifelong latent infections characterized by episodes of reactivation. Clinical manifestations span a wide spectrum, from neonatal infections and cancer to encephalitis and severe opportunistic disease in immunocompromised individuals. The most prevalent HHVs are the α-herpesviruses (HSV-1 and HSV-2), which are responsible for orolabial and genital herpes. HSV-1 affects an estimated 3.8 billion people under the age of 50 (64% worldwide) (1), while HSV-2 infects about 520 million individuals aged 15-49 (13%) (2, 3).

Drug discovery for HHVs has centered largely on nucleoside analogs that inhibit the viral DNA polymerase. For HSV, the mainstay therapy is acyclovir (ACV) and its prodrug valaciclovir (VCV), which effectively reduces the severity and frequency of genital and orolabial lesions (4, 5). However, ACV, approved in 1982 (6) and VCV, approved in 1995 (7), do not prevent viral reactivation, viral shedding or partner transmission, nor are they effective against the β- (HCMV, HHV6A/B and HHV7) or γ- (EBV, KSHV) HHVs. Ganciclovir (licensed in 1989 (8)), and its oral prodrug valganciclovir (approved in 2001 (9)), are the first-line antivirals for HCMV infections, although their use is limited by off-target toxicities such as myelosuppression (10, 11). The only newly approved agent targeting HHVs since 2001 has been letermovir (12), which inhibits the terminase. However, letermovir has major limitations including narrow spectrum (HCMV only), lack of approval for treatment of active disease, rapid emergence of resistance and clinically significant drug-drug interactions (13–16). These limitations highlight the ongoing need to develop anti-HHV therapeutics that act on novel viral targets.

Since 1978 it has been recognized that HSV-1 encodes a 5’ → 3’ alkaline deoxyribonuclease (UL12) (17). All HHVs encode a well-conserved ortholog of UL12 (18–21), several of which (HCMV UL98, EBV BGLF5 and KSHV SOX) have been shown to be essential for viral growth (22–26). HSV-1 UL12 is the best-characterized alkaline nuclease (AN), and we have shown that it plays a critical role in the synthesis of HSV genomes that can be packaged into infectious virions (26). Interestingly, UL12 collaborates with the HSV-1 single-strand DNA binding protein/annealase, ICP8, to mediate recombination-dependent replication of viral DNA (27–29). UL12 has also been shown to promote the maintenance of replication fork progression (30). Interestingly, HSV-1 also expresses a truncated form of the AN known as UL12.5 that lacks the disordered tail of the full-length protein but retains the core catalytic domain and exonuclease activity (31). Unlike the full-length protein, UL12.5 localizes to the mitochondria during infection (32). From a drug discovery perspective, UL12 is an attractive target due to its essential role in viral DNA replication.

UL12 and its HHV AN orthologs are members of a large family of two metal ion-dependent (TMID) enzymes that share a dependency on magnesium ions for catalytic activity (33–35). The presence of a well-defined active site also underscores its potential for the development of novel antiviral agents. We previously identified small-molecule inhibitors of HSV-1 UL12 by exploiting its two-metal ion catalytic center (26, 33). We have now established a platform to target the highly conserved HHV ANs which involves: 1) screening a proprietary library of flexible, metal-directed inhibitors; 2) optimizing substitution patterns of initial hits; 3) enhancing molecular rigidity; and 4) conducting off-target profiling and synthetic refinement to mitigate liabilities. Drug discovery efforts targeting TMID enzymes in HIV and influenza have successfully yielded selective and safe inhibitors in the past (36–39), demonstrating that enzymes of this class can serve as viable drug targets.

## Results and Discussion

We previously reported inhibitors of HSV-1 UL12, including a series of natural and synthetic α-hydroxytropolones that demonstrated inhibition of *in vitro* nuclease activity (26). From an in-house compound library, we also identified a number of 8-hydroxyquinolines (8-HQ), exemplified by AK-157 and AK-166, that inhibited UL12 in the micromolar range and showed antiviral activity against both HSV-1 and HCMV (33); however, these compounds demonstrated some cytotoxicity, leading to reduced selectivity indexes (SI) (Fig. 1). Building on this work, we have identified a new scaffold and produced a novel series of amido substituted hydroxypyridinones (HPs) which exhibit excellent antiviral activity and desirable selectivity indexes.

**Figure 1:**
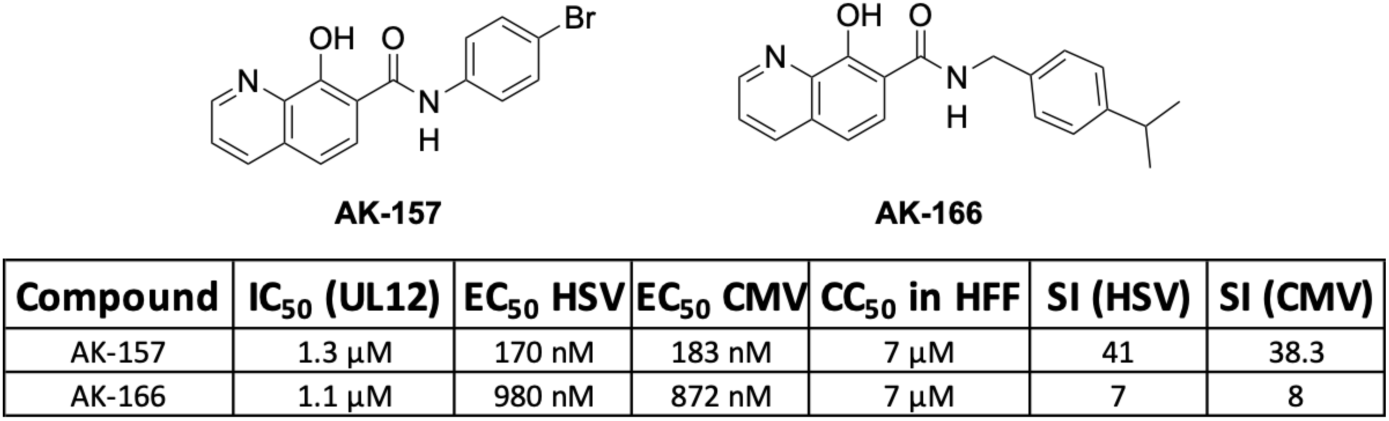
Structure and biological activity of representative hydroxyquinoline inhibitors.

### Determination of the crystal structure of UL12.5, the catalytic domain of HSV-1 UL12

Although crystal structures of γ-herpesvirus alkaline nucleases, such as BGLF5 in its apo form (40) and SOX in both apo and DNA/RNA-bound forms (41–43) have been previously solved, no α- or β-HHV AN structures have been reported. In this study, we present the first crystal structure of HSV-1 AN (UL12 residues 126-626) from an α-HHV. Efforts to crystallize full-length UL12 were hindered by the presence of a disordered N-terminal region comprising 126 amino acids. Therefore, the truncated protein (UL12 residues 126-626; also referred as UL12.5), was expressed in *E. coli* system with an N-terminal 12x His-tag and a TEV protease cleavage site to enable affinity purification. Despite its limited solubility, we successfully purified the protein using immobilized metal affinity and size exclusion chromatography, yielding approximately 1 mg of protein from a 1.5 L culture. This preparation of UL12.5 retained exonuclease activity at levels comparable to the full-length protein (data not shown). Subsequent crystallization trials yielded diffraction-quality crystals in the apo form, with the structure determined at 2.46 Å resolution via molecular replacement (Table S1).

Initial attempts to solve the UL12.5 structure by molecular replacement using available homologous structures, including SOX (41) and BGLF5 (40), were unsuccessful, likely due to limited sequence identity and structural divergence. Molecular replacement was ultimately successful using an AlphaFold2-predicted model of UL12.5 as the search template (44) which yielded a clear solution and well-defined electron density (Fig. 2A and 2B). The final model includes residues 1-73, 83-137, 144-166, 196-308, 317-418, 429-477 with the partial tag on the N-terminus (residues -8 to 0) and contains one molecule in the asymmetric unit. The overall architecture closely resembles that of the γ-herpesvirus nucleases SOX and BGLF5, adopting a canonical nuclease fold with distinct N- and C-terminal domains (Fig. 2A) (40, 41). The structural alignment of UL12.5 with SOX (PDB:3FHD) and BGLF5 (PDB:2W45) yielded RMSD values of 1.54 Å (205 atoms) and 1.63 Å (253 atoms), respectively (Fig. 2C). The catalytic triad, comprising Glu154, Asp 214 and Glu238 (residues Glu280, Asp340, and Glu364 respectively in full length-UL12) with clear electron density, is positioned within a crevice formed at the central core. However, no electron density corresponding to the bound Mg^2+^ ions was observed, despite their expected role in catalysis. Interestingly, while one Mg^2+^ is clearly resolved in the structure of SOX (3FHD), it is similarly absent in the apo structure of BGLF5 (2W45), indicating that metal ion binding may be substrate-dependent or influenced by subtle differences in structural dynamics among herpesvirus nucleases. Notably, the side chains of two residues of the catalytic triad, Glu154 and Asp214 in UL12.5 are oriented away from the metal-binding site in comparison to the SOX structure (Fig. 2D).

**Figure 2:**
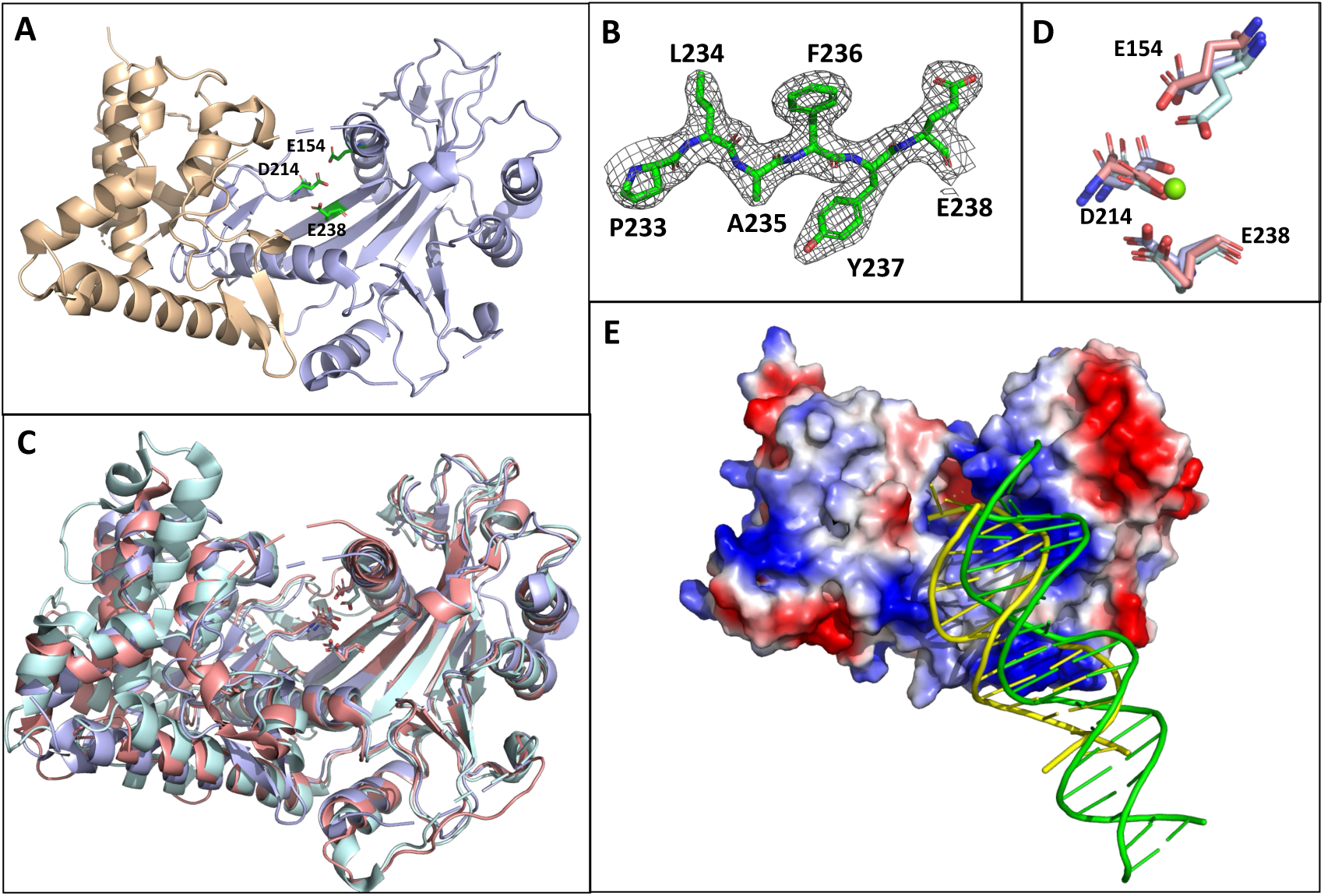
Structural analysis of HSV-1 alkaline nuclease UL12.5. (A) Overall structure of UL12.5, with the N-terminal domain shown in wheat and the core C-terminal domain in light blue. The conserved catalytic triad is depicted as green sticks. (B) Representative 2Fo-Fc electron density map contoured around residues 233-238. (C) Structural superposition of UL12.5 (light blue) with the Epstein-Barr virus alkaline nuclease BGLF5 (2W45: brick red) and the Kaposi’s sarcoma-associated herpesvirus alkaline nuclease SOX (3FHD: cyan). (D) The catalytic triads of UL12.5, BGLF5, and SOX are depicted as light blue, brick red, and cyan sticks, respectively, highlighting their relative positions and conformations. The Mg^2+^ ion coordinated in the SOX structure is represented as a green sphere. (E) Electrostatic surface representation of UL12.5, colored according to surface charge. Double-stranded DNA from structurally aligned DNA-bound complexes of KSHV SOX (green) and bacteriophage λ-exonuclease Redα (yellow) is shown to indicate the predicted DNA-binding path along the positively charged surface of UL12.5.

Comparative structural analysis shows that HSV-1 UL12 preserves the conserved 5′-phosphate-binding architecture characteristic of EBV BGLF5, KSHV SOX, and bacteriophage λ-exonuclease (Redα) (40, 45–47). In Redα, the DNA-bound structure reveals anchoring of the 5′-terminal phosphate within a deeply buried, positively charged pocket that is essential for substrate positioning and processive catalysis (48). An analogous anion-binding pocket is present in EBV BGLF5 and KSHV SOX, where sulfate, phosphate, or formate ions occupy the same site in apo and DNA-bound structures. Whereas SOX and BGLF5 contain a conserved Ser-Ser-Ser stretch within motif I, UL12 instead features a Thr-Ala-Ser sequence, a composition that better mirrors Redα and indicates functional rather than structural divergence (Fig. S1). Consistent with this conservation, UL12 displays clear electron density within the phosphate-binding pocket modeled as a formate ion, paralleling the anion-binding modes observed across the family. Together, these structures establish that UL12 retains the core framework for 5′-phosphate recognition shared with λ-exonuclease and other γ-herpesvirus nucleases, while subtle differences in residue substitutions may modulate substrate engagement and catalytic efficiency.

Our initial molecular docking efforts to design inhibitors relied on homology models of UL12 derived from the KSHV SOX structure. However, these models failed to accurately recapitulate key structure-activity relationships in the enzyme active site. The newly determined UL12.5 crystal structure from this work provided us a far more robust foundation for structure-based drug design.

### Molecular Modeling with the UL12.5 structure

In order to conduct docking studies with our newly solved UL12.5 structure, it was necessary to model in the active site divalent magnesium ions and bound waters. Structural modeling of UL12.5 was guided by the crystal structure of Redα bound to DNA (PDB: 3SM4) (48). Although a nucleic acid-bound structure is available for the orthologous KSHV SOX protein (42), we used λ Redα (48) due to its closer functional correspondence with UL12.5, as SOX also degrades mRNA – reflecting distinct active-site features. Proteins were aligned by superimposing conserved metal-coordinating residues (Fig. 3A). Following superposition, Redα protein and all non-bound waters were removed, leaving only active site waters and magnesium ions. Dihedral angles of the active site residues were adjusted as needed and the UL12.5 protein was prepared for docking using ProteinPrep in the Schrodinger Maestro suite (49–52).

**Figure 3.**
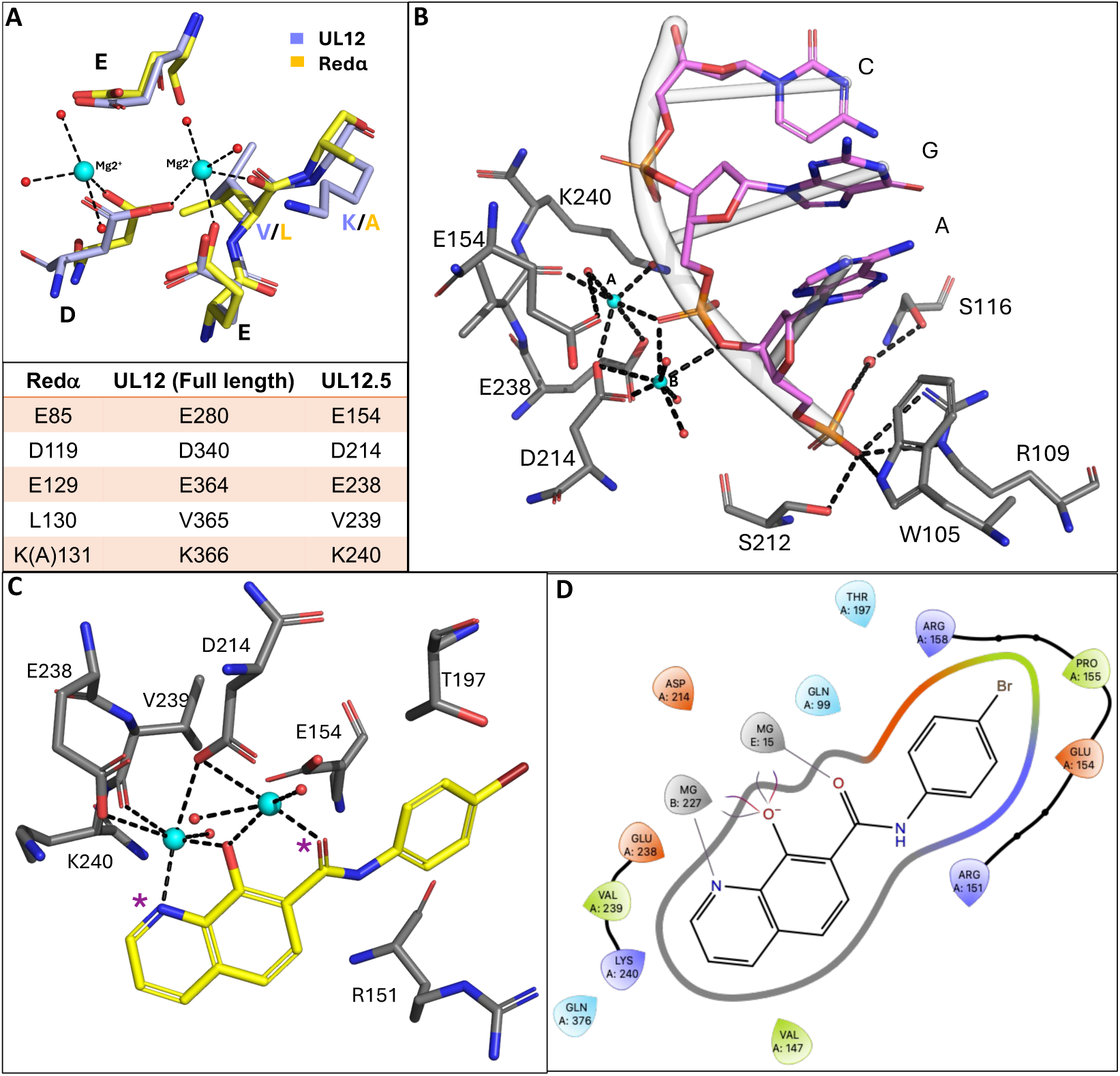
Structural comparison of the UL12.5 active site and docking of substrate and inhibitor. (**A**) Top panel: Structural superposition of conserved active-site residues from UL12.5 (blue) and λ Redα (yellow), shown as sticks. Residue 131 in λ Redα was mutated from Lys to Ala to prevent DNA cleavage; Bottom panel: Alignment of conserved active-site residues from λ Redα, full-length UL12, and UL12.5; residue numbering for UL12.5 is offset by 126 residues relative to full-length UL12. (**B**) Docking model of a trinucleotide containing a free 5′ phosphate bound to UL12.5, illustrating extensive interactions between the nucleotide bases and active-site residues, either directly or mediated by Mg^2+^ ions and water molecules. (**C**) Predicted binding orientation of the first-generation 8-hydroxyquinoline inhibitor AK-157 within the UL12.5 active site. Purple stars indicate water molecules displaced upon ligand binding. (**D**) Two-dimensional interaction diagram highlighting key contacts between UL12 and AK-157. *In panels **B-D**, UL12/UL12.5 active-site residues are shown as dark gray sticks, DNA bases as magenta sticks with the backbone in light gray, inhibitors as yellow sticks, Mg^2+^ ions and water molecules as cyan and red spheres respectively*.

To evaluate the model, a trinucleotide (AGC) bearing a free 5’ - phosphate was prepared with Schrodinger LigPrep (pH = 7+/-2) and docked to the protein using the Schrodinger Maestro Glide flexible docking (Fig. 3B) (52). Notable high affinity interactions include: (1) binding of the 5’-terminal phosphodiester bond to both magnesium ions, positioning it adjacent to the K240/magnesium **A**-bound nucleophilic water which performs the bond cleavage, (2) binding of the 5’-terminal phosphodiester oxygen to magnesium **B**, (3) a network of H-bonds between the 5’-free phosphate and residues W105, R109, S116 (via water), and S212 (53). Additionally, W105 is at an appropriate distance to engage the 5’ terminal nucleobase through hydrophobic π interactions.

### Docking of first generation hydroxyquinoline inhibitors

As previously reported, we identified a series of 8-hydroxyquinolines (HQs) that inhibited UL12 enzymatic function and demonstrated good antiviral activity against HSV-1 in cell culture (33). Extensive docking was performed with HQ inhibitors (i.e. AK-157, Fig. 1) to help elucidate the mode of binding and orientation of the molecule in the active site. Iterative modeling suggested that binding of the inhibitor (Fig. 3C) would promote the displacement of two bound water molecules from the active site. AK-157 interacts primarily through binding of the heteroatom triad with the magnesium ions, while the *p*-bromophenyl amide side chain sits in a hydrophobic region created by the hydrocarbon portion of the sidechains of residues R151, E154, and T197 (Fig. 3C, 3D and S2B).

### Implications of sequence homology for Pan-HHV activity

The HHV alkaline nucleases share significant homology (Fig. S2A), including identical active site catalytic arrays denoted by blue stars (UL12 PD-(D/E)XK motif: L339, D340, E364, V365, K366 with E280 (Fig. S2A, and 3B) as well as the 5’-phosphate binding pocket (WRSS denoted by green stars in Fig. S2A and S2B). We rationalized that potency of these compounds could be enhanced by engaging the unique 5’ terminal phosphate binding pocket (WRSS), which is not found in eukaryotic or prokaryotic exonucleases (48). Given the planar mode of binding of the HQs (Fig. S2B), we determined that a different scaffold would be required to access the terminal phosphate binding pocket. We selected a hydroxypyridinone (HP) chemotype, which would be more suitable for targeted design as the substitution on the nitrogen of the 4-pyridinone could be readily exchanged using well-established chemistry. Modeling indicated that aromatic groups in this region could potentially form contacts with the WRSS region and, thus, we prepared analogs bearing N-alkyl, aryl and benzyl substitutions on the pyridinone ring. Although this first iteration of HPs was inferior to the most potent HQs, the cellular CC_50_s were clearly improved (Fig. S2D). Molecular docking of representative compounds (i.e. DD-I-38) indicated that the pyridinone substitution was insufficiently flexible for accessing the WRSS pocket (Fig. S2C). Iterative molecular modeling determined that an azacycle spacer could effectively position the pendent aromatic groups proximal to the WRSS site. Subsequently, we prepared a series of HP analogs, including 4, 5, and 6-membered rings using the route shown in Scheme 1.

New compounds were screened against the UL12 enzyme, followed by antiviral evaluation (HSV-1) in cell culture. HPs that exhibited strong antiviral activity (EC_50_ < 500nM) were prioritized for additional evaluation (Fig. 4).

**Figure 4:**
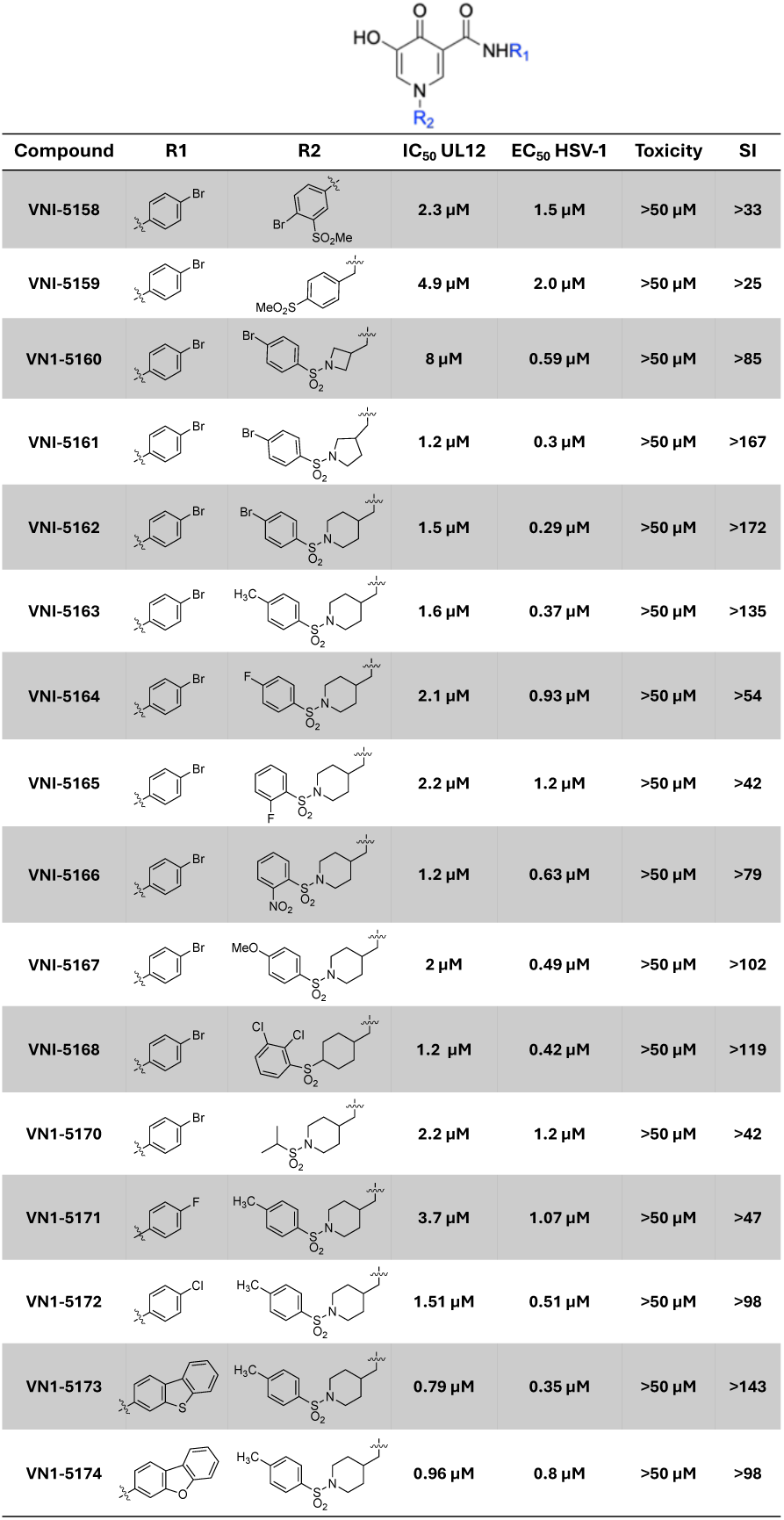
Chemical structure, UL12 inhibition values, antiviral activity against HSV-1 and corresponding cytotoxicity (at 24 hr) for representative HP inhibitors.

The sulfonamide design improved both target level activity and SI. Once baseline activities for the new series of compounds had been established, induced-fit molecular docking was performed to aid in developing a refined SAR model. Figure 5 shows representative sulfonamide class inhibitor VNI-5168 docked to the UL12.5 structure. The binding of the hydroxypyridinone core is dominated by interactions between the planar oxygen triad and the two divalent magnesium ions; however, the non-planar distortion of the aminomethyl piperidine group allows the aromatic sulfonamide to access the WRSS pocket, unlike the earlier HP inhibitors (Fig. S2C). Molecular docking predicted that the aromatic sulfonamides would engage this region through a combination of H-bonding to the sulfonamide oxygen atoms, π-interactions with the W105-R109 cation-π complex and H-X hydrogen bonding. The most active sulfonamide inhibitors (VNI-5162, 5163, 5167 and 5168) were selected for more detailed evaluation.

**Figure 5:**
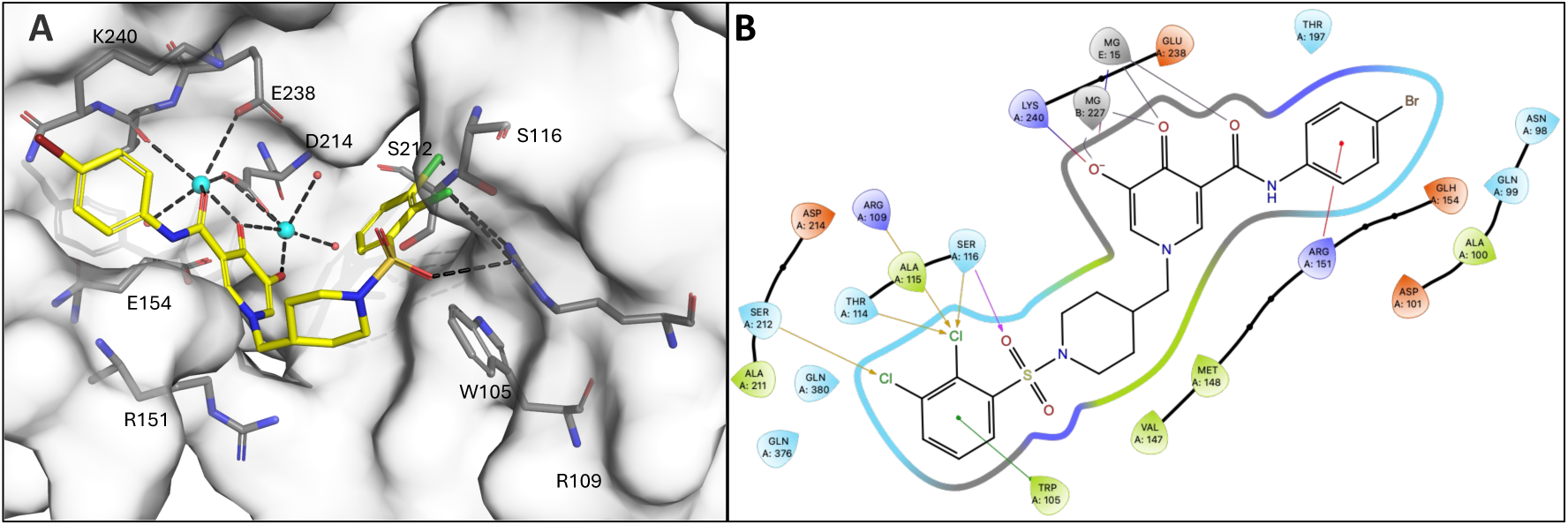
Docking of a representative sulfonamide inhibitor to UL12.5. (A) Docking model of the sulfonamide-class inhibitor VNI-5168 bound to the UL12.5 active site. The active-site pocket is shown in surface, with catalytic and ligand-interacting residues depicted as dark gray sticks. VNI-5168 is shown as yellow sticks, highlighting extensive interactions with UL12.5 residues and engagement of the WRSS pocket. Mg^2+^ ions and coordinating water molecules are depicted as cyan and red spheres respectively. (B) Two-dimensional ligand interaction diagram summarizing key contacts between UL12.5 and VNI-5168.

### Testing for breadth of coverage

As described above, there is a high level of active site homology across all seven HHV exonucleases. In principle, similar architectures could allow broad spectrum inhibition from a single compound. To better evaluate this potential, we compared the degree of inhibition of enzyme activity for the four select compounds between HSV-1 UL12 and two additional ANs, HCMV UL98, a β-herpesvirus and KSHV SOX, the γ-family homolog (Fig. 6A). Although all four analogs at a single fixed 5 µM concentration demonstrated complete inhibition of both UL12 and UL98, the activity against KSHV SOX was more variable. VNI-5162 and VNI-5168, which bear halogen substitutions on the pendent sulfonamide, exhibited strong inhibitory activity. In contrast, inhibition of SOX by the methyl (VNI-5163) and methoxy (VNI-5167) analogs was significantly lower. The observed disparity between the γ-(SOX) and α- (UL12)/ β- (UL98) ANs is potentially related to an additional RNA endonuclease activity of the γ orthologs.

**Figure 6:**
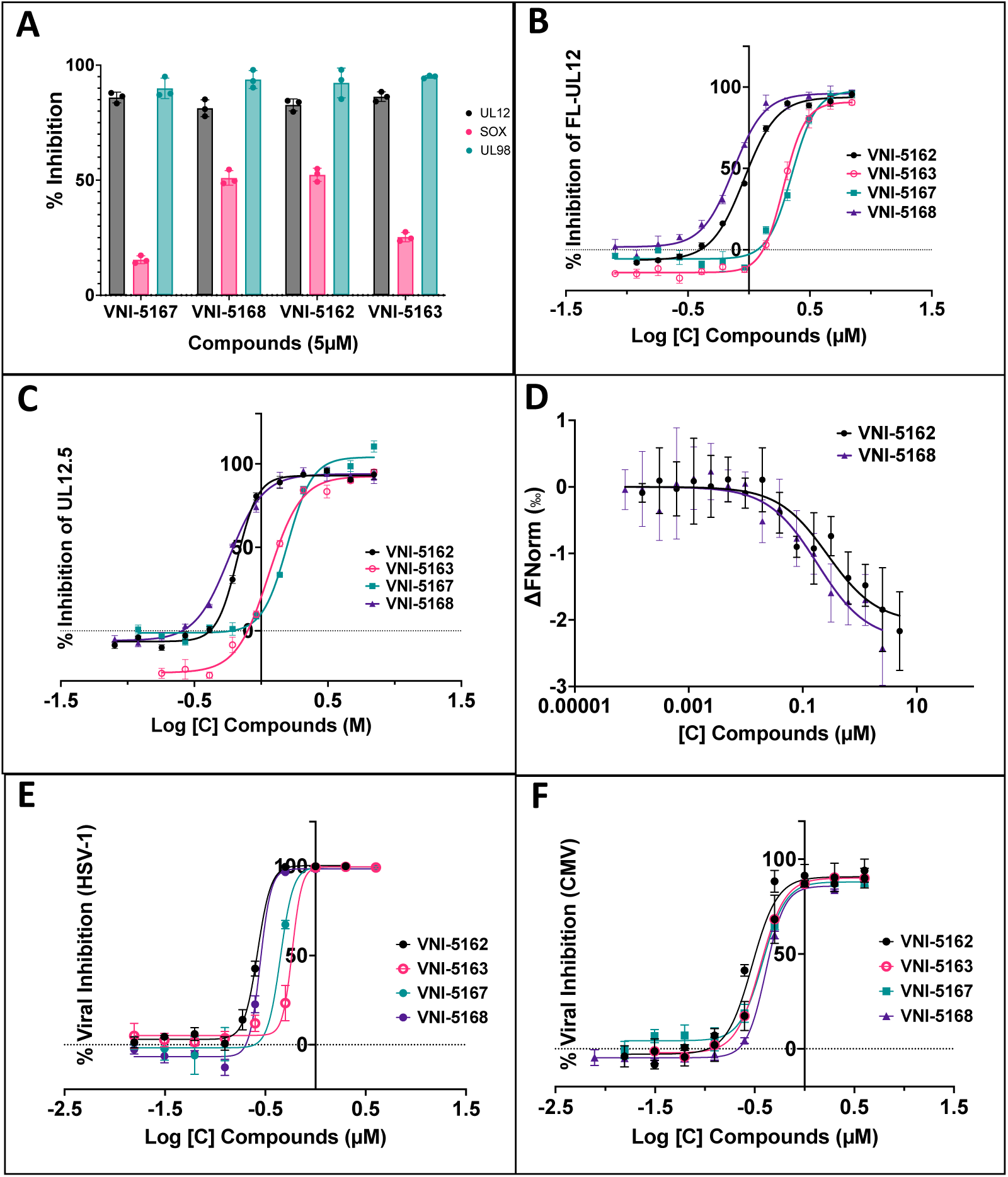
Biochemical and antiviral characterization of lead UL12 inhibitors. (A) Percentage inhibition of HSV-1 UL12, HCMV UL98, and KSHV SOX, representative alkaline nucleases from the alpha-, beta-, and gamma-herpesvirus subfamilies, respectively. Nuclease activity was measured using a PicoGreen-based assay in the presence of 5 µM inhibitor. Background fluorescence originating from the inhibitors was subtracted from the corresponding total relative fluorescence unit (RFU) values, and percent inhibition was calculated relative to the uninhibited DMSO control. Data represent the mean of three independent experiments, with each experiment consisting of 2–3 technical replicates. (B-F) Dose-response analyses for enzymatic inhibition, binding and antiviral activity. IC_50_ values for full-length (FL)-UL12 (B) and UL12.5 (C) were determined using the PicoGreen-based nuclease assay. (D) Binding affinities between UL12 and the indicated inhibitors were measured by microscale thermophoresis (MST), with normalized fluorescence plotted as a function of compound concentration. Data were fitted to a K_d_ binding model using MO.Affinity Analysis software (v3.0.5; NanoTemper Technologies) and plotted using GraphPad Prism. Data shown are from three independent experiments, each performed in technical duplicate. EC_50_ values for HSV-1 (E) and HCMV (F) were determined in HFF cells infected with fluorescently tagged viruses in the presence of increasing concentration of the compound and antiviral activity was evaluated using high-content image-based assay. *Percentage inhibition in panels B, C, E and F was calculated relative to the uninhibited DMSO control. IC_50_ and EC_50_ values were obtained by nonlinear regression analysis in GraphPad Prism using a four-parameter logistic model (log[inhibitor] versus response, variable slope). Data in panels B, C, E and F represent the mean of three independent experiments, with each experiment comprising 2-3 technical replicates*.

### Detailed characterization of lead sulfonamides

The four compounds were resynthesized and the inhibition of UL12 was re-confirmed, yielding IC_50_ values ranging from 0.59-2.1 µM (Table 1, column 1) with similar inhibitory profile observed against UL12.5 (Table 1, column 2 and Fig. 6B, 6C). Additionally, the equilibrium dissociation constants for the binding of the inhibitor to the enzyme (K_i_) values were extrapolated from the UL12 K_m_ (Fig. S3) and IC_50_s using the Cheng-Prusoff equation (54). The resulting sub-micromolar K_i_ values reflect robust inhibition of this nuclease (Table 1, column 3). However, it is important to note that UL12 is a processive enzyme, and interpretation of IC_50_/K_i_ values can be more complicated than those observed in more traditional distributive enzymes that follow classical Michaelis-Menten kinetics. In contrast, processive exonucleases employ a multi-turnover mechanism whereby multiple nucleotide excisions occur per each DNA-binding event. This often results in an under- or over-estimation of the true K_i_. To provide an independent assessment of target-engagement, dissociation constants (K_d_) for two lead inhibitors, VNI-5168 and VNI-5162, were determined using microscale thermophoresis (MST) (55, 56). As shown in Table 1 (column 4), direct binding was observed between UL12 and these two compounds. Titration of the labeled protein with increasing concentrations of the inhibitors yielded K_d_ values of 0.145 [0.069-0.304] µM for VNI-5168, and 0.249 [0.114-0.545] µM for VNI-5162 (Fig. 6D). These high binding affinity values align closely with our extrapolated K_i_ values (Table 1, column 3), further validating potent interactions between the sulfonamide inhibitors and UL12.

**Table 1:** Enzyme inhibition, antiviral efficacy, cytotoxicity, and selectivity indices of lead inhibitors targeting HSV-1 and HCMV.

|  | 1 | 2 | 3 | 4 | 5 | 6 | 7 | 8 | 9 | 10 |
| --- | --- | --- | --- | --- | --- | --- | --- | --- | --- | --- |
| VNI # | IC<sub>50</sub> [μM]<br>UL12 | IC<sub>50</sub> [μM]<br>UL12.5 | K<sub>i</sub> [μM]<br>UL12 | K<sub>d</sub> (μM)<sup>‡</sup><br>UL12 | EC<sub>50</sub> [μM]<br>HSV (fluor.) | EC<sub>50</sub> [μM]<br>CMV | EC<sub>50</sub> [μM]<br>HSV (yield) | CC<sub>50</sub><sup>*</sup><br>[μM] | SI<sup>**</sup><br>(HSV) | SI<br>(CMV) |
| 5162 | 0.97±0.09 | 0.68±0.02 | 0.28±0.03 | 0.249 | 0.30±0.03 | 0.23±0.03 | 0.11±0.01 | >100 | >330 (>900) | >430 |
| 5163 | 1.98±0.13 | 1.14±0.04 | 0.55±0.01 | ND | 0.51±0.09 | 0.33±0.04 | 0.23±0.05 | >10 | >19 (>43) | >30 |
| 5167 | 2.13±0.15 | 1.52±0.15 | 0.61±0.04 | ND | 0.42±0.09 | 0.42±0.04 | 0.14±0.01 | >100 | >230 (>710) | >230 |
| 5168 | 0.63±0.10 | 0.57±0.02 | 0.18±0.03 | 0.145 | 0.29±0.05 | 0.21±0.03 | 0.10±0.01 | >100 | >340 (>1000) | >470 |
| 5173 | 0.87±0.13 | ND | 0.25±0.04 | ND | 0.55±0.04 | 0.29±0.01 | 0.25±0.04 | >50 | >90 (>200) | >170 |
| 5174 | 1.14±0.16 | ND | 0.33±0.05 | ND | 0.60±0.05 | 0.33±0.06 | 0.27±0.01 | >50 | >80 (>180) | >150 |

<sup>‡</sup> MST derived K<sub>d</sub> values were obtained by global nonlinear regression with 68.3% confidence intervals [CI] of [0.114-0.545] for 5162 and [0.069-0.305] for 5168.
<sup>\*</sup>CC<sub>50</sub> values were reported at 65 hrs
<sup>\*\*</sup>Values in parentheses indicates SI calculated using EC<sub>50</sub> HSV-1 from yield assay IC<sub>50</sub> and EC<sub>50</sub> values are reported as mean ± SD from independent experiments.

### Antiviral activity of lead sulfonamides

The lead compounds described above were further evaluated against HSV-1 and HCMV using a fluorescence-based assay. HFF cells were infected with recombinant viruses expressing fluorescently tagged late gene products. Because late gene expression depends on ongoing viral DNA replication, this readout serves as a direct indicator of active replication. HFF cells were infected with an HSV-1 recombinant virus, eYFP-ICP0/gC-mCherry, generously provided by Dr. Colin Crump (Cambridge, UK) (57). A multiplicity of infection (MOI) of 1 PFU/cell was used to ensure that majority of the cells were infected, and expression of mCherry-tagged glycoprotein gC, encoded by the UL44 gene was measured using Cytation5 high-content microscopy. To assess inhibition of HCMV, HFF cells were infected with TB40/E-mCherry-UL99eGFP, generously provided by Eain A. Murphy (Upstate Medical University, Syracuse, NY) (58). Expression of GFP-tagged pp28 encoded by the late gene, UL99, was measured.

The lead compounds exhibited antiviral activity against HSV-1, with EC_50_ values comparable to the K_ᵢ_ values observed for UL12 inhibition (Table 1, column 5 and Fig. 6E). These compounds also showed significant inhibition of HCMV, consistent with their potency against the UL98 enzyme (Table 1, column 6 and Fig. 6F). We also tested antiviral activity against HSV-1 using a viral yield assay in which cells were infected at a low MOI (0.05 PFU/cell) for 65 h to test the ability of the virus to spread. In this assay, compounds produced a 2-3 fold stronger effect compared to the fluorescence assay (Table 1, column 7 Fig. S4A). This is likely due to the ability of the virus to undergo multiple rounds of infection at the low MOI used in this experiment. No significant cytotoxicity was observed at 24 hrs post-treatment, while some increase was evident at 65 hrs (Table 1, column 8, Figure S4B). However, for most compounds, even at the longer time-points, the CC_50_ for each compound was not reached at the highest compound concentration tested (100 µM). This resulted in selectivity indexes (SI) between 19-470 fold for fluorescence assays and even higher for yield assays (Table 1, column 9 and 10).

Exploration of SAR in the initial HQ series suggested a strong preference for halo aromatic units on the amido sidechain frequently seen in other nuclease inhibitors (59, 60). However, modeling suggested that alternative hydrophobic moieties could also be well tolerated in the pocket. To evaluate alternative substitutions, two additional analogs were prepared bearing a dibenzothiophene (VNI-5173) and dibenzofuran (VNI-5174) sidechain. These extended analogs exhibited strong inhibition of the recombinant UL12 enzyme and correspondingly strong antiviral activity with limited toxicity, suggesting additional freedom-to-operate in the region of the inhibitor structure (Table 1, Fig. S5).

## Conclusions

Extensive genetic and biochemical studies support a model in which HSV DNA synthesis proceeds through a recombination-dependent single-strand annealing pathway mediated by a two-component viral recombinase consisting of the alkaline nuclease UL12 and the SSAP ICP8 (28, 29, 61). The HSV-1 Exo/SSAP recombinase is reminiscent of related two-component recombination systems that are widely conserved among DNA viruses infecting bacteria, protozoa, plants, mammals and insects (62, 63). The preservation of viral Exo/SSAP mechanism suggests that they play a significant role in the evolutionary success of these viruses, supporting the choice of this complex as a novel target for drug discovery (26, 33).

Analysis of UL12 mutants lacking exonuclease activity, demonstrated a significant defect in the production of infectious viral progeny (26), suggesting that pharmacological inhibition of nuclease activity could be antiviral. To test this hypothesis, we leveraged our high-resolution structure of the catalytic domain of UL12 to design potent active site inhibitors. The resulting compounds show strong antiviral activity against HSV-1 with favorable selectivity (Table 1). Our structure provides the first view of an α-herpesvirus alkaline nuclease and establishes a robust platform for structure-based docking, SAR analysis and rational inhibitor design. Leveraging the high conservation across herpesvirus alkaline nucleases facilitated the design of broadly active antiviral inhibitors. Compounds that inhibited HSV-1 UL12 nuclease activity and viral replication also potently inhibited HCMV. Although inhibition of the KSHV SOX nuclease is more variable, several analogs effectively target all three enzymes, thus establishing the feasibility of developing pan-herpesvirus antivirals. While this study has been done against the human herpesviruses, it may also have broad veterinary applications including feline, bovine and equine herpesviruses. Ongoing efforts focus on probing drivers of selectivity between viral and key cellular nucleases as well as validating potent leads in animal models of infection.

## Materials and Methods

### Bacterial constructs

The pETM6T1-SOX vector was generously provided by Dr. Tracey E. Barrett. Codon-optimized UL12 and UL98 genes were synthesized by GenScript and subcloned into the pETM6T1-SOX backbone by replacing the SOX coding sequence with either UL12 or UL98. The NusA tag was subsequently removed and pETM6T2-UL12.5 construct was created by deleting 126 amino acids (378 bp) from the N-terminal region using NEB’s Q5® Site-Directed Mutagenesis Kit (NEB #E0554).

### Protein expression

All proteins bearing an N-terminal His tag were expressed in *Escherichia coli* BL21. For protein expression, a starter culture was prepared by inoculating cells into 100 mL of LB/2YT broth (Invitrogen) supplemented with kanamycin (25 µg/mL) and incubated overnight at 37°C with shaking. The following day, a 1:100 dilution of the overnight culture was transferred into 1.5-2L of fresh media and grown at 37°C until the optical density at 600 nm (A_600_) reached 0.6-0.8. Protein expression was induced by the addition of 1 mM isopropyl β-D-1-thiogalactopyranoside (IPTG), and cultures were incubated overnight at 18°C. Cells were harvested by centrifugation and stored at -80°C until further use.

### Protein purification

#### HSV-1 UL12.5

Cell pellets were resuspended in buffer A (25 mM Tris-HCl, pH 8.5; 300 mM NaCl; 5 mM β-mercaptoethanol) supplemented with 10 mM imidazole and 1 mM PMSF, followed by sonication on ice. Lysates were clarified by centrifugation at >26,000 × g for 1 hour, and the supernatant was filtered through a 0.45 µm syringe filter before incubation with of pre-equilibrated Ni-NTA resin (50% slurry) at 4°C for 1 hour with gentle shaking. Bound proteins were eluted using buffer A supplemented with 300 mM imidazole. Fractions containing the highest purity proteins were pooled, concentrated, and subjected to size exclusion chromatography using a Superdex 200 column pre-equilibrated with UL12 storage buffer (25 mM Tris-HCl, pH 8.5; 300 mM NaCl; 10% glycerol; 5 mM DTT). Fractions containing pure protein were verified by SDS-PAGE gel, pooled, concentrated, and stored at -70 °C until further use.

#### HCMV UL98

Cell pellets from 1.5 L of culture expressing UL98 were resuspended in buffer B (25 mM Tris-HCl, pH 8.5; 200 mM NaCl) supplemented with cOmplete protease inhibitor, DNase I, 1 mM PMSF, and 5 mM β-mercaptoethanol. Cells were lysed by sonication on ice, and the lysate was clarified by centrifugation at >26,000 × g for 1 h. The supernatant was filtered through a 0.45 µm syringe filter and incubated with pre-equilibrated Ni-NTA resin at 4 °C for 1 h with gentle agitation. Bound proteins were eluted with buffer B supplemented with 500 mM imidazole. The affinity tag was removed by addition of TEV protease during overnight dialysis against buffer B containing 2 mM DTT. The dialyzed sample was applied to a HiTrap Q FF column equilibrated in buffer B with 2 mM DTT and eluted using a linear NaCl gradient from 50 mM to 1 M. Fractions containing UL98 were pooled, concentrated, and further purified by size-exclusion chromatography on a Superdex 200 column equilibrated in storage buffer (25 mM Tris-HCl, pH 8.5; 300 mM NaCl; 10% glycerol; 5 mM DTT). Fractions containing pure protein were verified by SDS-PAGE gel, pooled, concentrated, and stored at −70 °C until further use.

#### KSHV SOX

KSHV SOX was purified as described in Bagneris et al (42).

### Protein crystallization

Initial crystallization trials with 192 conditions from NeXtal JCSG Core Suite (II and III) were conducted at a protein concentration of 5 mg/mL using the sitting drop vapor diffusion method at 4 °C. Crystallization screening was automated with a Gryphon crystallization robot (Art Robbins Instruments), where 200 nL of protein solution was mixed with 200 nL of crystallization solution and equilibrated against 80 µL of reservoir solution. Initial crystal hits were obtained from the Core Suite II, conditions G8 containing 0.16 M ammonium sulfate, 0.08 M sodium acetate (pH 4.6), 20% (w/v) PEG 4000, 20% (v/v) glycerol. To improve crystal quality, the above conditions were further optimized using the hanging drop vapor diffusion method with 2 µL each of protein and reservoir solutions. Diffraction-quality crystals grew within a few days from the well solution comprising 0.12 M ammonium sulfate, 0.08 M sodium acetate (pH 5.0), 15% (w/v) PEG 4000, 10% (v/v) glycerol, and 4% DMSO. Current efforts are focused on generating co-crystals of the UL12 with the most potent inhibitors.

### Data collection and structure solution

X-ray diffraction data were collected at the National Synchrotron Light Source II, using the 17-ID-1 (AMX) beamline at Brookhaven National Laboratory. Diffraction data were processed with XDS (64) for indexing and integration, followed by space-group determination with Pointless (65), which identified P 2₁ 2₁ 2₁ as the correct space group. Scaling and merging were performed in Aimless (66). The structure of UL12 was determined by molecular replacement using Phaser (67) as part of the CCP4i2 suite of programs (68) employing an AlphaFold-predicted model as the initial search template (69, 70). Iterative rounds of model building and refinement were conducted using Coot (71) and Refmac (72). Comprehensive data collection parameters, refinement statistics, and final structural metrics are summarized in Table S1.

### PicoGreen alkaline nuclease assay

Alkaline exonuclease (UL12) activity was assessed using the PicoGreen fluorescence assay, as previously described (33). In brief, 25-µL reaction mixtures containing 20 mM Tris-HCl (pH 8.2), 40 mM NaCl, 1 mM MgCl_2_, 1 mM dithiothreitol (DTT), 10 nM UL12, and either DMSO or varying concentrations of inhibitors (with final DMSO concentration of 1.6%) were preincubated at 25°C for 10 minutes. The nuclease reaction was initiated by the addition of 1 nM linearized pUC119 DNA, followed by a 10-minute incubation at 25°C. Reactions were quenched with 10 µL of 1.25 mM EDTA (pH 8.0). For fluorescence detection, 150 µL of PreciseGreen® dsDNA quantification reagent (Lumiprobe; diluted 1:500 in Tris-EDTA buffer, pH 7.5) was added to each quenched reaction. A total of 170 µL of the reaction mixture was transferred to a 96-well Flurotrac 200 black plate (Greiner Bio-One), and fluorescence was measured using a SpectraMax 3 plate reader with excitation and emission wavelengths of 480 nm and 520 nm, respectively.

The exonuclease activities of SOX and UL98 were assessed using similar assay conditions with a few variations. For SOX, reactions were carried out in a buffer containing 20 mM Tris-HCl (pH 8.8), 50 mM NaCl, 10 mM MgCl_2_, and 1 mM DTT.

For UL98, the reaction buffer consisted of 50 mM Tris-HCl (pH 8.8), 2 mM MgCl_2_, and 1 mM DTT. The final protein concentrations were 50 nM for SOX and 20 nM for UL98. All reactions were incubated at 37 °C for 10 minutes.

### Microscale Thermophoresis (MST)

Binding affinities of two most potent compounds, VNI-5168 and VNI-5162, with UL12 were determined using MST (55, 56). The protein was diluted to 200 nM in PBS-T buffer (137 mM NaCl, 2.5 mM KCl, 10 mM Na_2_HPO4, 2 mM KH_2_PO4, pH 7.4; 0.05 % Tween-20) supplemented with 2.5 mM MgCl_2_ and 2.5 mM DTT and was labeled with RED-tris-NTA 2^nd^ generation dye in the 1:2 ratio (100 nM dye, 200 nM protein) for 30 minutes at room temperature. Stock solutions of both compounds (20 mM in DMSO) were diluted to 5-10 μM in PBS-T buffer containing 3% DMSO, followed by 16 points, 1:1 serial dilution. Each ligand dilution was then mixed with an equal volume of labeled protein to yield final concentrations of 50 nM UL12 and 1.5% DMSO. After 20 minutes incubation time, all samples were centrifuged for 10 min at 12000 g in 4 °C and loaded into premium capillaries (Nanotemper Technologies). The measurements were performed on a NanoTemper MonolithNT.115 ^pico^ instrument. The initial fluorescence was uniform among the sixteen capillaries, and the thermographs showed no signs of aggregation or molecule adsorption to the capillary. Data was collected at 22°C, at 40% LED power and medium MST power. Three independent experiments were performed with technical duplicate. MO.Affinity Analysis software (v3.0.5, Nano Temper Technologies) was used to fit the data and to calculate the apparent K_d_ values with their respective 68.3% confidence intervals [CI] using a 1:1 binding model.

### Cells and viruses

African green monkey kidney epithelial cells (Vero) were obtained from ATCC and propagated in Dulbecco’s Modified Eagle Medium (DMEM) (Invitrogen, Carlsbad CA) supplemented with 5% fetal bovine serum (FBS, Gemini Bio-Products, Woodland CA). Human foreskin fibroblast (HFF) cells were purchased from ATCC and cultured in Eagle’s Minimal Essential Medium (MEM) (Invitrogen, Carlsbad, CA) supplemented with 10% FBS. Fluorescently labeled, recombinant eYFP-ICP0/gC-mCherry virus, a gift from Colin Crump (Cambridge, UK), expressing YFP tagged versions of ICP0 and mCherry tagged UL44 (gC) gene was used in HSV-1 inhibition assays (57). The TB40/E-*mCherry*-UL99*eGFP,* a recombinant HCMV TB40/E strain generously provided by Eain A. Murphy (Upstate Medical University, Syracuse, NY), expressing mCherry under SV40 promoter and GFP-tagged version of UL99 gene was used in HCMV inhibition assay (58).

### Antiviral experiments

HFF cells were seeded in 96-well clear plates in 5% FBS-MEM. At 24 h post seeding, cells were infected with eYFP-ICP0/gC-mCherry HSV-1 virus at a multiplicity of infection (MOI) of 1 PFU/cell for antiviral high-content image-based compound screen (fluorescent-based assay) or with MOI of 0.05 PFU/cell for yield-based assay and incubated for 20 h or 65 h, respectively, in the presence of various concentrations of the compounds or DMSO alone in 100 μL MEM at final concentrations of 2% FBS and 0.5% DMSO. For HCMV antiviral high-content image-based assays cells were infected at MOI of 2 PFU/cell with fluorescently labeled TB40/E-*mCherry*-UL99*eGFP* and incubated for 65 h under the same experimental setup as for HSV-1 virus.

For the HSV-1 inhibition assay, the entire plate was subjected to 2 freeze-thaw cycles, and viral yields were assessed by plaque assay on Vero cells as described previously (73). For HSV-1 and HCMV antiviral high-content image-based assays at 20 h or 65 h post infection, respectively, plates were fixed with 2% Paraformaldehyde for 15 min at room temperature (RT) in the dark. Plates were washed twice with phosphate buffered saline (PBS) and wells were overlayed with PBS. Images were acquired with Agilent BioTek Cytation5 high-content microscope using 4X air objective. For HSV-1 images were collected using Channel 1: YFP (excitation 500nm, emission 542nm), Channel 2: Texas Red (excitation 586nm, emission 647nm) and for HCMV Channel 1: GFP (excitation 469nm, emission 525nm), Channel 2: Texas Red (excitation 586nm, emission 647nm). Images from both channels were acquired, but only channel 1 images were analyzed for assessment of viral inhibition. For every image, mean intensity of the YFP (HSV-1) or GFP (HCMV) fluorescent representing true late gene expression in each virus was used for calculation corrected for the background from untreated cells. Signal obtained from DMSO treated infected cells was normalized to 100% infection (0% inhibition) and used for calculation of percent viral inhibition in infected cells treated with increasing concentrations of compound.

### Cytotoxicity assay

HFF cells were seeded in 96-well white plates in 5% FBS-MEM. At 24-h post seeding, cells were incubated in the presence of various concentrations of the compounds or DMSO alone in 100 μL of MEM at final concentrations of 2% FBS and 0.5% DMSO. At 24 h and 65 h post-treatment, cytotoxicity was assessed using CellTiter Glo (Promega, Madison WI) luciferase assay according to the manufacturer’s instructions.

### Software and programs used

Molecular modelling and docking were performed using ProteinPrep, LigPrep and Glide in Schrodinger Maestro suite (49–52). Multiple sequence alignment was performed using Clustal Omega (74, 75). Final structural figures were generated using PyMol Version 3.1.7.2 (Schrödinger, LLC). IC_50_ and EC_50_ values were obtained by nonlinear regression analysis using a four-parameter logistic model (log[inhibitor] versus response, variable slope) in GraphPad Prism version 10.6.1 for Mac OS X, GraphPad Software, Boston, Massachusetts USA, www.graphpad.com.

### General synthetic scheme

Compounds were synthesized as shown in Scheme 1. Commercially available pyrone 1 underwent aminolysis and ring transformation with the corresponding Boc-protected aminomethyl piperidine to afford protected hydroxypyridinone intermediate 2. Ester hydrolysis provided acid 3, which was coupled with the appropriate aniline to furnish amide intermediates 4. Subsequent Boc deprotection afforded free piperidine intermediates 5, which were functionalized with the corresponding sulfonyl chlorides to provide sulfonamide intermediates 6. Final debenzylation afforded the target hydroxypyridinone analogs.

**Scheme 1:**
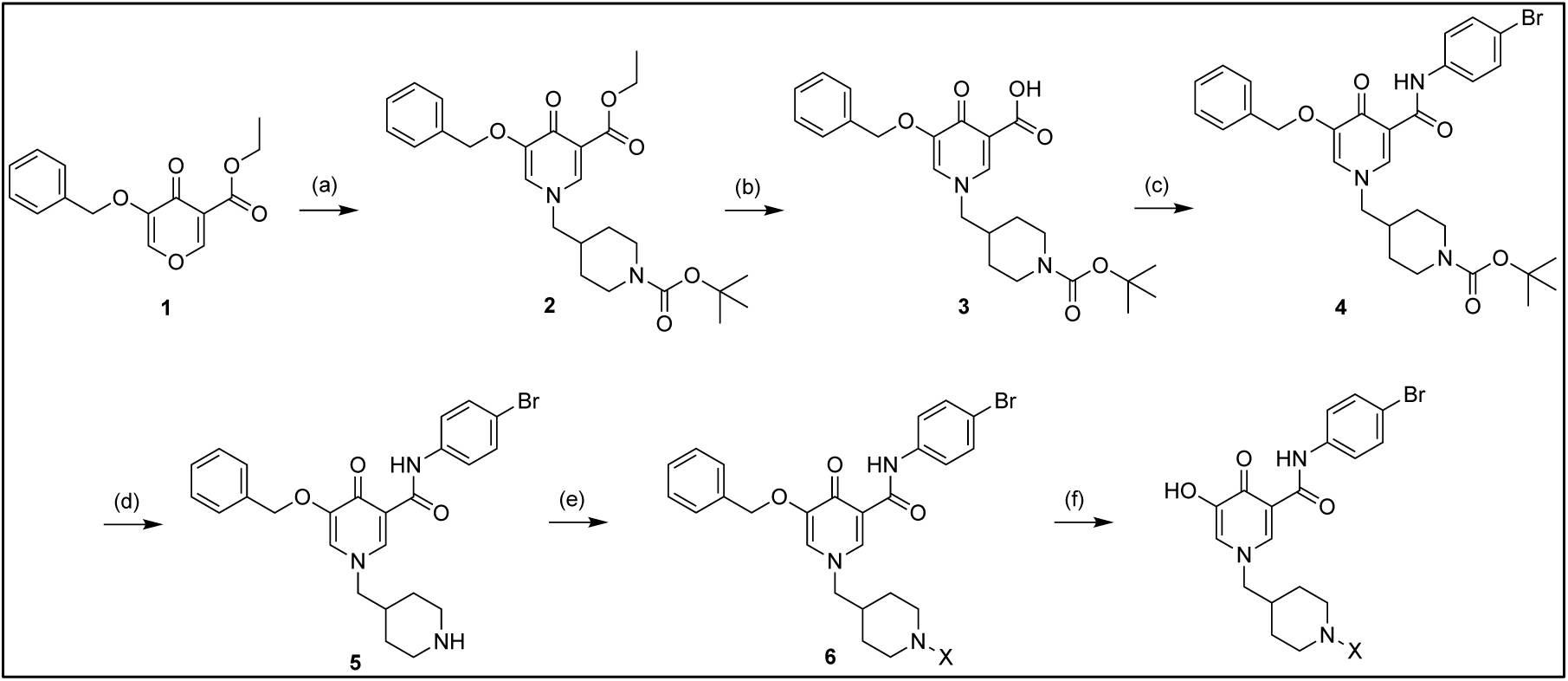
Reagents and conditions. (a) EtOH, HOAc, 2-amino-2-phenylethan-1-ol; (b) NaOH, EtOH, 80°C; (c) DIPEA, Pybop, DMSO, DMAP, 4-bromoaniline; (d) DCM, TFA; (e) DMF, DIPEA, X; (f) TFA 90°C.

## Acknowledgments

We thank the U.S. Department of Health and Human Services, and the NIH/NIAID provided funding (to S.K.W. and D.L.W.) under R56 AI173955. The funders had no role in study design, data collection and interpretation, or the decision to submit the work for publication. This research used resources at beamlines 17-ID-1 and 17-ID-2 of the National Synchrotron Light Source II, a U.S. Department of Energy (DOE) Office of Science User Facility operated for the DOE Office of Science by Brookhaven National Laboratory under Contract No. DE-SC0012704. The MST work was supported by grant S10OD028574 (to Dr. Heidi Erlandsen) for a Monolith NT.115pico system housed in the Center for Open Research Resources and Equipment at the University of Connecticut. We thank Dr. Chathura Abeywickrama for collecting HPLC data for the compounds. AI-assisted tools (ChatGPT and Microsoft Copilot) were used for language refinement of the manuscript.

## Author Contributions

N.S., X. X., R. S., C. R., S. K. K., J. K., X. C., D. D., L. W., D. W., and S. W. designed research; N.S., X. X., R. S., C. R., S. K.K., J. K., X. C., D. D., L. W. performed research; N.S., X. X., R. S., S. K.K., J. K., X. C., D. D., L. W. analyzed data; L. W., D. W., S. W. supervised research; and N. S., L. W., D. W., S. W. wrote the paper.

