## Supplemental data for "Viral Nuclease Inhibitors: Small molecule disruptors of the UL12 alkaline nuclease display broad anti-herpes virus activity"

**This PDF file includes:**

- Supporting Text
- Schemes S1 to S3
- Figures S1 to S5
- Tables S1
- Mass spectra for compounds
- HPLC data for compounds
- $^{1}$H NMR and $^{13}$C NMR spectra

#### SUPPORTING TEXT

##### Steady-state kinetics:

PicoGreen alkaline nuclease assay was used to obtain the Michaelis constant ($K_m$) for UL12 which then enabled the calculation of the equilibrium dissociation constant ($K_i$) of the UL12-inhibitor complex using the Cheng-Prusoff equation. The reactions were carried out in 96-well black, flat-bottom microplates at room temperature in a final volume of 200 µL. Linearized pUC119 served as the double-stranded DNA (dsDNA) substrate for the assays, with concentrations ranging from 0.25 nM to 1.875 nM, and UL12 was held at a fixed concentration of 5 nM. Prior to start of the reaction, the protein was preincubated in the dark with varying concentrations of pUC119 and PicoGreen in nuclease assay buffer containing 20 mM Tris-HCl (pH 8.2), 40 mM NaCl, 1 mM dithiothreitol (DTT). The final PicoGreen dilution was 1:115. The reaction was initiated by adding a final concentration of 1 mM $MgCl_2$. Fluorescence was recorded on a SpectraMax plate reader using Ex/Em = 480/520 nm, with measurements taken every 1 min 30 s for 60 min. A no-enzyme reaction mixture was included as a negative control and measured under identical conditions. $K_m$ values were obtained by plotting initial reaction velocities as a function of DNA substrate concentration and fitting the data to the Michaelis-Menten equation using GraphPad Prism. For inhibition analysis, $K_i$ values were estimated from experimentally determined $IC_{50}$ values using the Cheng-Prusoff relationship:

$$K_i = \frac{IC50}{1+\frac{[S]}{K_m}}$$

which corrects the $IC_{50}$ for the substrate concentration [S] used in the assay and the measured $K_m$ of UL12 for the DNA substrate.

**Synthesis:**

Chemicals and solvents were purchased from commercial suppliers (Sigma Aldrich, Acros Organics, TCI Chemicals, Fisher Scientific, and AAblocks Inc) and were used as received unless otherwise stated. Compound 1 was purchased from Sigma Aldrich. Air and moisture-sensitive reactions, when required, were carried out under an inert atmosphere of argon in oven-dried glassware that was allowed to cool under argon. TLC analyses were performed on Sorbent Technologies silica gel HL TLC plates with fluorescent indicators (UV254). Developed plates were air-dried, and compounds were visualized under ultraviolet light (254 or 365 nm). Flash column chromatography was carried out using silica gel as the stationary phase. The $^{1}H$ and $^{13}C$ NMR spectra were recorded on Bruker instruments at 500 MHz. Chemical shifts are reported in ppm and are referenced to residual solvent signals ($CDCl_3$: δ 7.24 and 77.23 ppm for $^{1}H$ and $^{13}C$; $CD_3OD$: δ 4.78, 3.31, and 49.15 ppm). Signal splitting patterns are described as singlet (s), doublet (d), triplet (t), quartet (q), multiplet (m), broad (b), or a combination thereof. Coupling constants (J) are quoted to the nearest 0.1 Hz. Liquid chromatography mass spectrometry (LC–MS) analyses were performed using a Waters ACQUITY UPLC system coupled to a Xevo G2-XS q-TOF mass spectrometer. Chromatographic separations were carried out on a Waters HSS T3 column (150 mm). LC-MS analyses were performed at a flow rate of 0.30 mL/min using a linear gradient of mobile phase B in A, where solvent A was water and solvent B was acetonitrile. The gradient was held at 5% B for 0.5 min, increased linearly from 5% to 95% B over 5.5 min, and held at 95% B for 0.5 min, followed by re-equilibration to 5% B until 8.0 min. Purity analyses were performed by reversed-phase high performance liquid chromatography (RP-HPLC) using a Shimadzu Prominence 20 instrument equipped with a Waters XSelect HSS T3 C18 column (5 µm, 4.6 mm × 250 mm) and monitored using a photodiode array (PDA) detector over a wavelength range of 190-800 nm. Mobile phase A consisted of water containing 0.05% trifluoroacetic acid (TFA), and mobile phase B consisted of acetonitrile containing 0.05% TFA. The gradient elution was performed at a flow rate of 2.50 mL/min as follows: 35% A/65% B from 0-2 min, 5% A/95% B from 2-14 min, held at 5% A/95% B until 17 min, returned to 35% A/65% B by 19 min, and equilibrated at 35% A/65% B until 21 min. The total run time was 22 min. Samples were prepared in HPLC-grade DMSO prior to analysis. The purity of all final compounds tested in biological assays was ≥95%.

**Procedure for Aminolysis and Ring Transformation to Dihydropyridines.** Piperidinylmethyl-substituted dihydropyridine ester (**2**). To a suspension of 2-amino-2-phenylethan-1-ol (5.15 g, 25.5 mmol) in EtOH (40 mL) and HOAc (20 mL) was added the corresponding pyran starting material (3.0 g, 10.9 mmol). The reaction mixture was stirred at room temperature overnight and subsequently heated at 90 °C for 4 h. Upon completion, as monitored by TLC, the solvents were removed under reduced pressure. The residue was purified by flash column chromatography on silica gel (1-3% MeOH in 1:1 DCM/EtOAc) to afford compound 2 as a light yellow solid (3.5 g, 68%). $^{1}$H NMR (500 MHz, $CDCl_3$) δ 8.55 (s, 1H), 8.28 (s, 1H), 7.49-7.24 (m, 5H), 4.98 (s, 2H), 4.26 (q, J = 7.1 Hz, 2H), 3.89-3.25 (m, 6H), 1.68-1.26 (m, 6H), 1.42 (s, 9H), 1.32 (t, J = 7.1 Hz, 3H). $^{13}$C NMR (126 MHz, $CDCl_3$) δ 181.5, 163.8, 158.6, 154.8, 146.0, 142.0, 135.1, 128.5, 128.2, 127.2, 125.1, 79.8, 71.2, 62.3, 60.8, 44.1, 35.0, 31.7, 28.4, 14.1. HRMS (ESI) m/z $[M + H]^+$ calcd for $C_{26}H_{35}N_2O_6$ 471.2490; found 471.2565.

**Procedure for Ester Hydrolysis to Dihydropyridine.** Piperidinylmethyl-substituted dihydropyridine carboxylic acid (**3**). To a solution of compound 2 (3.5 g, 7.43 mmol) in ethanol (20 mL) was added an aqueous solution of NaOH (18.59 mmol). The resulting mixture was heated at 78 °C and stirred overnight. Upon completion, as monitored by TLC, the reaction mixture was cooled to room temperature, and the ethanol was removed under reduced pressure. The residue was diluted with water (20 mL), stirred, and cooled in an ice bath. The solution was acidified to pH 6 by dropwise addition of 1 N HCl (approximately 19 mL), resulting in the precipitation of a white solid. The solid was collected by filtration, washed with cold water, and dried under vacuum to afford compound 3 as a white solid (2.8 g, 85%). $^{1}$H NMR (500 MHz, DMSO-$d_6$) δ 8.59 (s, 1H),

8.02 (s, 1H), 7.52-7.24 (m, 5H), 5.13 (s, 2H), 4.07-3.94 (m, 2H), 3.42-3.25 (m, 4H), 2.11 (m, 1H), 1.68-1.50 (m, 4H), 1.42 (s, 9H). $^{13}$C NMR (126 MHz, DMSO-d$_6$) δ 181.5, 165.9, 158.6, 154.8, 146.0, 142.0, 135.1, 128.5, 128.2, 127.2, 120.4, 79.8, 71.2, 62.3, 44.1, 35.0, 31.7, 28.4. HRMS (ESI) m/z [M + H]$^+$ calcd for $C_{24}H_{31}N_2O_6$, 443.2177; found 443.2196.

**Procedure for Amide Coupling of Dihydropyridine Carboxylic Acids.** Arylamide-substituted dihydropyridine (**4**). To a solution of compound 3 (2.45 g, 5.53 mmol) in anhydrous DMSO (12 mL) were added DIPEA (2.97 mL, 16.6 mmol) and PyBOP (3.46 g, 6.64 mmol). The mixture was stirred at room temperature for 15 min, after which 4-bromoaniline (1.14 g, 6.64 mmol) and DMAP (0.13 g, 1.1 mmol) were added. The reaction mixture was stirred at room temperature for 8 h. TLC analysis (DCM/EtOAc = 1:1) indicated that approximately 10% of the starting material remained; therefore, an additional portion of PyBOP (0.34 g, 0.65 mmol) was added, and the reaction was stirred overnight. The reaction was quenched with water and extracted with EtOAc. The combined organic layers were washed sequentially with saturated aqueous $NaHCO_3$ solution and brine, dried over anhydrous $Na_2SO_4$, filtered, and concentrated under reduced pressure. Purification by flash column chromatography on silica gel (15% EtOAc in DCM) afforded compound 6 as a light yellow solid (2.45 g, 74%). $^1$H NMR (500 MHz, DMSO-d$_6$) δ 12.94 (s, 1H), 8.55 (s, 1H), 8.52 (s, 1H), 8.28 (s, 1H), 7.84-7.72 (m, 4H), 7.45-7.24 (m, 5H), 4.98 (s, 2H), 3.89 (d, J = 3.5 Hz, 2H), 3.42-3.25 (m, 4H), 2.10 (m, 1H), 1.69-1.52 (m, 4H), 1.42 (s, 9H). $^{13}$C NMR (126 MHz, DMSO-d$_6$) δ 181.5, 163.8, 158.6, 154.8, 146.0, 142.0, 135.1, 131.9, 128.5, 128.2, 127.4, 127.2, 121.8, 120.4, 116.3, 79.8, 71.2, 62.3, 44.1, 35.0, 31.7, 28.4. HRMS (ESI) m/z [M + H]$^+$ calcd for $C_{30}H_{35}BrN_3O_5$, 596.1735; found 596.1743.

**Procedure for Boc Deprotection of Piperidine Derivatives.** Secondary piperidine dihydropyridine amide (**5**). To a solution of compound 4 (2.45 g, 4.1 mmol) in dichloromethane (6 mL) was added trifluoroacetic acid (3.14 mL, 41.0 mmol). The reaction mixture was stirred at room temperature for 2 h. Upon completion, the mixture was concentrated under reduced pressure. The residue was suspended in water (30 mL), and the pH was adjusted to >8 by the addition of 1

N aqueous NaOH, resulting in the formation of a white precipitate. The solid was collected by filtration, washed with water, and dried under vacuum to afford compound 5 as a white solid (2.00 g, 93%). $^1$H NMR (500 MHz, DMSO-$d_6$) δ 12.92 (s, 1H), 8.59 (s, 1H), 8.28 (s, 1H), 8.52 (s, 1H), 7.72 (m, 2H), 7.46-7.24 (m, 7H), 5.09 (s, 2H), 4.10-4.09 (m, 2H), 3.88 (d, J = 3.2 Hz, 2H), 2.94-2.64 (m, 4H), 1.99 (m, 1H), 1.84-1.67 (m, 4H). $^{13}$C NMR (126 MHz, DMSO-$d_6$) δ 181.5, 163.8, 158.6, 146.0, 142.0, 135.1, 131.9, 128.5, 128.2, 127.4, 127.2, 121.8, 120.4, 116.3, 71.2, 62.3, 46.6, 35.0, 33.4. HRMS (ESI) m/z: $[M + H]^+$ calcd for $C_{25}H_{27}BrN_3O_3$, 496.1236; found 496.1275.

**General Procedure for N-Functionalization of Piperidine Dihydropyridine Amides.** To a solution of the piperidine dihydropyridine amide (compound **5**, 1.0 mmol) in anhydrous DMF (5 mL) was added DIPEA (0.52 mL, 3.0 mmol), followed by portionwise addition of the corresponding sulfonyl chloride (1.1 mmol) at room temperature. The reaction mixture was stirred at room temperature overnight. Upon completion, the reaction mixture was poured into water (40 mL), and the resulting precipitate was collected by filtration. The crude solid was dissolved in EtOAc (50 mL) and washed sequentially with 5% aqueous $NaHCO_3$ and brine, dried over anhydrous $Na_2SO_4$, filtered, and concentrated under reduced pressure. The residue was further purified by trituration with methanol to remove a minor impurity observed by TLC. The purified solid was collected by filtration and dried under vacuum to afford the N-functionalized products (50-80% yield).

N-(4-bromophenyl)-5-(benzyloxy)-1-[4-bromo-3-(methylsulfonyl)phenyl]-3,6-dioxo-1,2,3,6-tetrahydropyridine-4-carboxamide (**6**). To a solution of compound **5** (0.50 g, 1.0 mmol) in anhydrous DMF (5 mL) was added DIPEA (0.52 mL, 3.0 mmol), followed by portionwise addition of methanesulfonyl chloride (0.085 mL, 1.1 mmol) at room temperature. The reaction mixture was stirred at room temperature overnight. Upon completion, the reaction mixture was poured into water (40 mL), and the resulting precipitate was collected by filtration. The crude solid was dissolved in EtOAc (50 mL) and washed sequentially with 5% aqueous $NaHCO_3$ and brine, dried over anhydrous $Na_2SO_4$, filtered, and concentrated under reduced pressure. The residue was further purified by trituration with methanol to remove a minor impurity observed by TLC. The purified solid was collected by filtration and dried under vacuum as an off-white solid (517 mg, 82% yield). $^{1}$H NMR (500 MHz, DMSO-$d_6$) δ 8.59 (s, 1H), 8.48 (s, 1H), 8.1-8.00 (m, 2H), 7.86-7.66 (m, 3H), 7.45-7.28 (m, 7H), 4.97 (s, 2H), 3.10 (s, 3H). $^{13}$C NMR (126 MHz, DMSO-$d_6$) δ 181.5, 163.8, 158.6, 144.3, 142.0, 136.2, 135.1, 133.4, 131.9, 128.5, 128.2, 127.4, 127.2, 121.8, 120.7, 120.4, 117.7, 116.3, 116.0, 71.2, 44.2. HRMS (ESI) m/z [M + H]$^+$ calcd for $C_{26}H_{21}Br_2N_2O_5S$, 630.9573; found 630.9565.

N-(4-bromophenyl)-1-[(4-(methylsulfonyl)benzyl)]-3-(benzyloxy)-4-oxo-1,4-dihydropyridine-2-carboxamide (**7**). Yellow solid (67% yield) $^{1}$H NMR (500 MHz, DMSO-d6) δ 3.08 (s, 3H), 4.92-5.03 (m, 4H), 7.24-7.50 (m, 9H), 7.59-7.78 (m, 4H), 8.30 (s, 1H), 8.50 (s, 1H). $^{13}$C NMR (126 MHz, DMSO-$d_6$) δ 181.5, 163.8, 158.6, 146.0, 142.0, 137.4, 136.5, 135.1, 131.9, 129.4, 128.5, 128.2, 127.8, 127.4, 127.2, 121.8, 120.4, 116.3, 71.2, 53.5, 44.5. HRMS (ESI) m/z [M + H]$^+$ calcd for $C_{27}H_{24}BrN_2O_5S$ 565.0570; found 565.0576.

N-(4-bromophenyl)-1-[(2-(4-bromophenylsulfonyl)azetidin-1-yl)methyl]-3-(benzyloxy)-4-oxo-1,4-dihydropyridine-2-carboxamide (**8**). Yellow solid; yield 70%. $^{1}$H NMR (500 MHz, DMSO-$d_6$) δ 2.12-2.47 (m, 5H), 3.40 (m, 1H), 3.90 (d, J = 3.2 Hz, 2H), 4.98 (s, 2H), 7.24-7.45 (m, 7H), 7.59 (m, 2H), 7.66-7.79 (m, 4H), 8.28 (s, 1H), 8.52 (s, 1H). $^{13}$C NMR (126 MHz, DMSO-$d_6$) δ 181.5, 163.8, 158.6, 146.0, 142.0, 135.6, 135.1, 133.1, 131.9, 129.5, 128.5, 128.2, 127.8, 127.4, 127.2, 121.8, 120.4, 116.3, 71.2, 48.5, 39.0, 31.8, 29.8. HRMS (ESI) m/z [M + H]$^+$ calcd for $C_{29}H_{26}Br_2N_3O_5S$ 686.40; found 686.46.

N-(4-bromophenyl)-1-[1-(4-bromophenylsulfonyl)pyrrolidin-2-yl]-3-(benzyloxy)-4-oxo-1,4-dihydropyridine-2-carboxamide (**9**). Yellow solid; yield 60%. $^{1}$H NMR (500 MHz, DMSO-$d_6$) δ 1.96-2.29 (m, 2H), 2.60 (m, 1H), 3.17-3.49 (m, 3H), 3.86-4.12 (m, 3H), 4.93-5.03 (m, 2H), 7.24-7.46 (m, 7H), 7.56 (m, 2H), 7.63-7.78 (m, 4H), 8.29 (s, 1H), 8.49 (s, 1H). $^{13}$C NMR (126 MHz, DMSO-$d_6$) δ 181.5, 163.8, 158.6, 146.0, 142.0, 135.1, 133.6, 133.1, 131.9, 129.5, 128.5, 128.2, 127.8,

127.4, 127.2, 121.8, 120.4, 116.3, 71.2, 57.6, 56.5, 47.3, 37.1, 28.9. HRMS (ESI) m/z $[M + H]^+$ calcd for $C_{30}H_{28}Br_2N_3O_5S$ 700.0111; found 700.0124.

N-(4-bromophenyl)-1-[1-(4-bromophenylsulfonyl)pyrrolidin-2-yl]-3-(benzyloxy)-4-oxo-1,4-dihydropyridine-2-carboxamide (**10**). Yellow solid; yield 64%. $^1$H NMR (500 MHz, DMSO-$d_6$) δ 1.61-1.79 (m, 4H), 2.10 (m, 1H), 3.22 (m, 2H), 3.59 (m, 2H), 3.89 (d, J = 3.2 Hz, 2H), 4.98 (s, 2H), 7.24-7.46 (m, 7H), 7.56 (m, 2H), 7.63-7.78 (m, 4H), 8.28 (s, 1H), 8.49 (s, 1H). $^{13}$C NMR (126 MHz, DMSO-$d_6$) δ 181.5, 163.8, 158.6, 146.0, 142.0, 135.1, 133.1, 132.1, 131.9, 129.5, 128.5, 128.2, 127.8, 127.4, 127.2, 121.8, 120.4, 116.3, 71.2, 62.3, 44.5, 35.0, 31.7. HRMS (ESI) m/z $[M + H]^+$ calcd for $C_{31}H_{30}Br_2N_3O_5S$ 714.0267; found 714.0271.

N-(4-bromophenyl)-1-[1-(4-methylphenylsulfonyl)pyrrolidin-2-yl]-3-(benzyloxy)-4-oxo-1,4-dihydropyridine-2-carboxamide (**11**). Yellow solid; yield 60%. $^1$H NMR (500 MHz, DMSO-$d_6$) δ 1.61-1.79 (m, 4H), 2.12 (m, 1H), 2.32 (s, 3H), 3.22 (m, 2H), 3.59 (m, 2H), 3.88 (d, J = 3.2 Hz, 2H), 7.25-7.43 (m, 4H), 7.63-7.84 (m, 5H), 8.51 (s, 1H). $^{13}$C NMR (126 MHz, DMSO-$d_6$) δ 184.1, 163.8, 147.4, 146.0, 144.9, 142.0, 132.1, 131.9, 129.4, 127.7, 127.4, 121.8, 120.4, 116.3, 62.3, 44.5, 35.0, 31.7, 21.4. HRMS (ESI) m/z $[M + H]^+$ calcd for $C_{32}H_{34}BrN_3O_5S$ 652.1455; found 652.1459.

N-(4-bromophenyl)-1-[1-(4-fluorophenyl)sulfonylpyrrolidin-2-yl]-3-(benzyloxy)-4-oxo-1,4-dihydropyridine-2-carboxamide (**12**). Yellow solid; yield 64%. $^1$H NMR (500 MHz, DMSO-$d_6$) δ 1.61-1.79 (m, 4H), 2.10 (m, 1H), 3.22 (m, 2H), 3.59 (m, 2H), 3.89 (d, J = 3.2 Hz, 2H), 4.98 (s, 2H), 7.24-7.46 (m, 7H), 7.56-7.78 (m, 6H), 8.28 (s, 1H), 8.49 (s, 1H). $^{13}$C NMR (126 MHz, DMSO-$d_6$) δ 181.5, 165.7, 163.8, 158.6, 146.0, 142.0, 135.1, 132.1, 131.9, 128.5, 128.2, 127.8, 127.4, 127.2, 121.8, 120.4, 116.7, 116.3, 71.2, 62.3, 44.5, 35.0, 31.7. HRMS (ESI) m/z $[M + H]^+$ calcd for $C_{31}H_{30}BrFN_3O_5S$ 654.1073; found 654.1064.

N-(4-bromophenyl)-1-[1-(2-fluorophenyl)sulfonylpyrrolidin-2-yl]-3-(benzyloxy)-4-oxo-1,4-dihydropyridine-2-carboxamide (**13**). Yellow solid; yield 61%. $^1$H NMR (500 MHz, DMSO-$d_6$) δ 1.61-1.79 (m, 4H), 2.10 (m, 1H), 3.22 (m, 2H), 3.59 (m, 2H), 3.89 (d, J = 3.2 Hz, 2H), 4.98 (s, 2H), 7.24-7.57 (m, 8H), 7.58-7.90 (m, 5H), 8.28 (s, 1H), 8.49 (s, 1H). $^{13}$C NMR (126 MHz, DMSO-$d_6$)

δ 181.5, 163.8, 159.2, 158.6, 146.0, 142.0, 136.2, 136.0, 135.1, 131.9, 129.5, 128.5, 128.2, 127.5, 127.4, 127.2, 121.8, 120.4, 117.3, 116.3, 71.2, 62.3, 44.5, 35.0, 31.7. HRMS (ESI) m/z $[M + H]^+$ calcd for $C_{31}H_{30}BrFN_3O_5S$ 654.1073; found 654.1075.

N-(4-bromophenyl)-1-[1-(2-nitrophenyl)sulfonylpyrrolidin-2-yl]-3-(benzyloxy)-4-oxo-1,4-dihydropyridine-2-carboxamide (**14**). Yellow solid; yield 55%. $^1H$ NMR (500 MHz, DMSO-$d_6$) δ 1.61-1.80 (m, 4H), 2.10 (m, 1H), 3.23 (m, 2H), 3.60 (m, 2H), 3.89 (d, J = 3.2 Hz, 2H), 4.98 (s, 2H), 7.24-7.46 (m, 7H), 7.66-7.92 (m, 4H), 7.96-8.15 (m, 2H), 8.28 (s, 1H), 8.49 (s, 1H). $^{13}C$ NMR (126 MHz, DMSO-$d_6$) δ 181.5, 163.8, 158.6, 146.0, 142.0, 137.8, 136.2, 135.1, 131.9, 129.5, 128.7-128.4 (m), 128.2, 127.5, 127.4, 127.2, 126.7, 121.8, 120.4, 116.3, 71.2, 62.3, 44.5, 35.0, 31.7. HRMS (ESI) m/z $[M + H]^+$ calcd for $C_{31}H_{30}BrN_4O_7S$ 682.0712; found 682.0720.

N-(4-bromophenyl)-1-[1-(4-methoxyphenyl)sulfonylpyrrolidin-2-yl]-3-(benzyloxy)-4-oxo-1,4-dihydropyridine-2-carboxamide (**15**). Yellow solid; yield 64%. $^1H$ NMR (500 MHz, DMSO-$d_6$) δ 1.61-1.79 (m, 4H), 2.10 (m, 1H), 3.22 (m, 2H), 3.59 (m, 2H), 3.89 (m, 2H), 3.84 (s, 3H), 4.98 (s, 2H), 6.97 (m, 2H), 7.24-7.46 (m, 7H), 7.59 (m, 2H), 7.72 (m, 2H), 8.28 (s, 1H), 8.49 (s, 1H). $^{13}C$ NMR (126 MHz, DMSO-$d_6$) δ 181.5, 163.9, 163.7, 158.6, 146.0, 142.0, 135.1, 132.1, 131.9, 128.5, 128.2, 127.8, 127.4, 127.2, 121.8, 120.4, 116.3, 114.5, 71.2, 62.3, 55.3, 44.5, 35.0, 31.7. HRMS (ESI) m/z $[M + H]^+$ calcd for $C_{32}H_{33}BrN_3O_6S$ 666.1273; found 666.1277.

N-(4-bromophenyl)-1-{[1-((2,3-dichlorophenyl)sulfonyl)piperidin-4-yl]methyl}-3-(benzyloxy)-4-oxo-1,4-dihydropyridine-2-carboxamide (**16**). Yellow solid (yield 60%). $^1H$ NMR (500 MHz, DMSO-$d_6$) δ 12.8-12.6 (br s, 1H), 8.45-8.20 (s, 1H), 7.90-7.20 (m, 12H, Ar-H), 5.00 (s, 2H, $OCH_2Ph$), 4.10-3.90 (m, 2H, $NCH_2$), 3.30-3.10 (m, 2H), 2.85-2.65 (m, 2H), 2.20-1.90 (m, 2H), 1.80-1.40 (m, 4H). $^{13}C$ NMR (126 MHz, DMSO-$d_6$) δ 181.5, 163.8, 158.6, 146.0, 142.0, 136.2, 135.1, 133.0, 131.9, 130.4, 129.6, 129.4, 128.9, 128.5, 128.2, 127.4, 127.2, 121.8, 120.4, 116.3, 71.2, 62.3, 44.5, 35.0, 31.7. HRMS (ESI) m/z: $[M + H]^+$ calcd for $C_{31}H_{29}BrCl_2N_3O_5S$, 704.0368; found 704.0360.

N-(4-bromophenyl)-1-[1-(isopropylsulfonyl)piperidin-4-yl]-3-(benzyloxy)-4-oxo-1,4-dihydropyridine-2-carboxamide (**17**). Yellow solid; yield 55%. $^{1}$H NMR (500 MHz, $CDCl_3$) δ 8.52 (s, 1H), 8.28 (s, 1H), 7.72 (d, J = 8.4 Hz, 2H), 7.46-7.24 (m, 7H), 4.98 (s, 2H), 3.89 (d, J = 3.2 Hz, 2H), 3.83-3.52 (m, 3H), 3.24 (ddd, J = 14.8, 6.8, 6.8 Hz, 2H), 2.10 (m, 1H), 1.79-1.60 (m, 4H), 1.35 (d, J = 7.0 Hz, 6H). $^{13}$C NMR ($CDCl_3$) δ 181.5, 163.8, 158.6, 146.0, 142.0, 135.1, 131.9, 128.5, 128.2, 127.4, 127.2, 121.8, 120.4, 116.3, 71.2, 62.3, 55.0, 44.5, 35.0, 31.7, 16.1. HRMS (ESI) m/z [M + H]$^+$ calcd for $C_{28}H_{33}BrN_3O_5S$ 602.1298; found 602.1306.

N-(4-fluorophenyl)-1-[1-(4-methylphenylsulfonyl)piperidin-4-yl]-3-(benzyloxy)-4-oxo-1,4-dihydropyridine-2-carboxamide (**18**). Yellow solid; yield 61%. $^{1}$H NMR (500 MHz, $CDCl_3$) δ 8.52 (s, 1H), 8.28 (s, 1H), 7.79-7.63 (m, 4H), 7.46-7.24 (m, 7H), 7.02 (d, J = 8.5 Hz, 2H), 4.98 (s, 2H), 3.89 (d, J = 3.2 Hz, 2H), 3.59 (ddd, J = 14.8, 2.8, 2.8 Hz, 2H), 3.22 (ddd, J = 14.8, 6.8, 6.8 Hz, 2H), 2.32 (s, 3H), 2.09 (m, 1H), 1.79-1.61 (m, 4H). $^{13}$C NMR ($CDCl_3$) δ 181.5, 163.8, 158.6, 157.8, 146.0, 144.9, 142.0, 135.1, 132.1, 129.4, 128.5, 128.2, 127.7, 127.4, 127.2, 120.6, 120.4, 115.1, 71.2, 62.3, 44.5, 35.0, 31.7, 21.4. HRMS (ESI) m/z [M + H]$^+$ calcd for $C_{32}H_{33}FN_3O_5S$ 590.2120; found 590.2128.

N-(4-chlorophenyl)-1-[1-(4-methylphenylsulfonyl)piperidin-4-yl]-3-(benzyloxy)-4-oxo-1,4-dihydropyridine-2-carboxamide (**19**). Yellow solid; yield 60%. $^{1}$H NMR (500 MHz, $CDCl_3$) δ 8.52 (s, 1H), 8.28 (s, 1H), 7.78-7.62 (m, 4H), 7.46-7.24 (m, 9H), 4.98 (s, 2H), 3.89 (d, J = 3.2 Hz, 2H), 3.59 (ddd, J = 14.8, 2.8, 2.8 Hz, 2H), 3.22 (ddd, J = 14.8, 6.8, 6.8 Hz, 2H), 2.32 (s, 3H), 2.09 (m, 1H), 1.79–1.61 (m, 4H). $^{13}$C NMR ($CDCl_3$) δ 181.5, 163.8, 158.6, 146.0, 144.9, 142.0, 135.1, 132.1, 129.4, 129.0, 128.5, 128.2, 127.7, 127.4, 127.2, 127.0, 121.6, 120.4, 71.2, 62.3, 44.5, 35.0, 31.7, 21.4. HRMS (ESI) m/z [M + H]$^+$ calcd for $C_{32}H_{33}ClN_3O_5S$ 606.1824; found 606.1830.

N-(benzothiophen-2-yl)-1-[1-(4-methylphenylsulfonyl)piperidin-4-yl]-3-(benzyloxy)-4-oxo-1,4-dihydropyridine-2-carboxamide (**20**). Yellow solid; yield 45%. ^1H NMR (500 MHz, $CDCl_3$) δ 8.50 (s, 1H), 8.29 (s, 1H), 8.05 (ddd, J = 8.1, 1.5, 0.4 Hz, 1H), 7.84-7.24 (m, 15H), 4.98 (s, 2H), 3.89 (d, J = 3.2 Hz, 2H), 3.59 (ddd, J = 14.8, 2.8, 2.8 Hz, 2H), 3.22 (ddd, J = 14.8, 6.8, 6.8 Hz,

2H), 2.32 (s, 3H), 2.09 (m, 1H), 1.79–1.61 (m, 4H). $^{13}$C NMR (126 MHz, $CDCl_3$) δ 181.5, 163.8, 158.6, 146.0, 144.9, 142.0, 139.6, 136.8, 135.9, 135.7, 135.1, 134.9, 132.1, 129.4, 128.5, 128.2, 127.7, 127.2, 127.0, 124.5, 122.9, 121.9, 121.2, 120.4, 118.1, 111.3, 71.2, 62.3, 44.5, 35.0, 31.7, 21.4. HRMS (ESI) m/z $[M + H]^+$ calcd for $C_{38}H_{36}N_3O_5S_2$ 678.2091; found 678.2083.

N-(dibenzofuran-2-yl)-1-[1-(4-methylphenylsulfonyl)piperidin-4-yl]-3-(benzyloxy)-4-oxo-1,4-dihydropyridine-2-carboxamide (**21**). Yellow solid; yield 46%. $^1$H NMR (500 MHz, $CDCl_3$) δ 8.50 (s, 1H), 8.29 (s, 1H), 7.96-7.61 (m, 7H), 7.55-7.16 (m, 9H), 4.98 (s, 2H), 3.89 (d, J = 3.2 Hz, 2H), 3.59 (ddd, J = 14.8, 2.8, 2.8 Hz, 2H), 3.22 (ddd, J = 14.8, 6.8, 6.8 Hz, 2H), 2.32 (s, 3H), 2.09 (m, 1H), 1.79–1.61 (m, 4H). $^{13}$C NMR (126 MHz, $CDCl_3$) δ 181.5, 163.8, 158.6, 156.1, 155.2, 146.0, 144.9, 142.0, 136.8, 135.1, 132.1, 129.4, 128.5, 128.2, 127.7, 127.5, 127.2, 124.3, 124.2, 122.6, 122.4, 121.2, 120.4, 118.1, 111.6, 110.3, 71.2, 62.3, 44.5, 35.0, 31.7, 21.4. HRMS (ESI) m/z $[M + H]^+$ calcd for $C_{38}H_{37}N_3O_6S$ 662.2319; found 662.2325.

**General Procedure for O-Deprotection of Benzyloxy Dihydropyridines.** Hydroxy-substituted dihydropyridine amides. To (e.g., compound **6**, 0.5 mmol scale) was added trifluoroacetic acid (2 mL), and the reaction mixture was heated at 90 °C for 2 h. Upon completion, as confirmed by TLC, the mixture was cooled to room temperature and concentrated under reduced pressure. The crude off-white solid was suspended in methanol (5 mL) and neutralized with 30% aqueous ammonia (0.1 mL). The resulting precipitate was collected by filtration. The material was further purified by suspension in dichloromethane (40 mL), stirring at room temperature for 30 min, and standing overnight. The purified solids were collected by filtration and dried under vacuum.

N-(4-bromophenyl)-5-hydroxy-1-[4-bromo-3-(methylsulfonyl)phenyl]-3,6-dioxo-1,2,3,6-tetrahydropyridine-4-carboxamide **VNI-5158**. To compound **6** (0.50 g, 0.83 mmol) was added trifluoroacetic acid (2 mL), and the reaction mixture was heated at 90 °C for 2 h. Upon completion, as confirmed by TLC, the mixture was cooled to room temperature and concentrated under reduced pressure. The crude off-white solid was suspended in methanol (5 mL) and neutralized with 30% aqueous ammonia (0.1 mL). The resulting precipitate was collected by filtration. The material was further purified by suspension in dichloromethane (40 mL), stirring at room temperature for 30 min, and standing overnight. The purified solid was collected by filtration and dried under vacuum to afford the final product as an off-white solid (250 mg, 62%). $^{1}$H NMR (500 MHz, DMSO-$d_6$) δ 12.61 (s, 1H), 8.58 (s, 1H), 8.13 (d, J = 8.2 Hz, 2H), 7.94 (s, 1H), 7.79 (d, J = 8.2 Hz, 2H), 7.72 (d, J = 8.3 Hz, 2H), 7.37 (d, J = 8.3 Hz, 2H), 3.10 (s, 3H). $^{13}$C NMR (126 MHz, DMSO-$d_6$) δ 171.01, 162.61, 149.77, 142.66, 141.16, 139.80, 138.28, 137.37, 132.40, 130.62, 126.73, 122.90, 122.04, 120.39, 116.25, 115.82, 42.37. HRMS (ESI) m/z $[M + H]^+$ calcd for $C_{19}H_{15}Br_2N_2O_5S$, 540.9068; found 540.9054.

N-(4-bromophenyl)-1-[(4-(methylsulfonyl)benzyl)]-3-hydroxy-4-oxo-1,4-dihydropyridine-2-carboxamide (**VNI-5159**). Yellow solid; yield 60%. $^{1}$H NMR (500 MHz, DMSO-$d_6$) δ 12.71 (s, 1H), 8.60 (s, 1H), 7.89 (d, J = 8.3 Hz, 2H), 7.68 (s, 1H), 7.56 (t, J = 7.5 Hz, 4H), 7.46 (d, J = 8.6 Hz, 2H), 5.39 (s, 2H), 3.13 (s, 3H). $^{13}$C NMR (126 MHz, DMSO-$d_6$) δ 184.1, 163.8, 147.4, 146.0, 142.0, 137.4, 136.5, 131.9, 129.4, 127.8, 127.4, 121.8, 120.4, 116.3, 53.5, 44.5. HRMS (ESI) m/z $[M + H]^+$ calcd for $C_{20}H_{18}BrN_2O_5S$ 477.0120; found 477.0117.

N-(4-bromophenyl)-1-[2-(4-bromophenylsulfonyl)azetidin-1-ylmethyl]-3-hydroxy-4-oxo-1,4-dihydropyridine-2-carboxamide (**VNI-5160**). Yellow solid; yield 66%. $^{1}$H NMR (500 MHz, DMSO-$d_6$) δ 12.77 (s, 2H), 8.37 (d, J = 2.3 Hz, 2H), 7.81 (d, J = 8.5 Hz, 3H), 7.69-7.59 (m, 7H), 7.52 (d, J = 8.9 Hz, 4H), 4.16 (d, J = 6.6 Hz, 4H), 3.74 (t, J = 8.4 Hz, 4H), 3.51 (t, J = 7.3 Hz, 3H), 2.97-2.88 (m, 1H). $^{13}$C NMR (126 MHz, DMSO-$d_6$) δ 169.51, 162.50, 149.20, 139.85, 138.02, 132.99, 132.56, 131.87, 129.99, 128.59, 127.73, 122.89, 121.50, 115.05, 114.78, 57.76, 52.99, 28.86. HRMS (ESI) m/z $[M + H]^+$ calcd for $C_{22}H_{20}Br_2N_3O_5S$ 597.9530; found 597.9464.

N-(4-bromophenyl)-1-[1-(4-bromophenylsulfonyl)pyrrolidin-2-yl]-3-hydroxy-4-oxo-1,4-dihydropyridine-2-carboxamide (**VNI-5161**). Yellow solid; yield 58%. $^{1}$H NMR (500 MHz, DMSO-$d_6$) δ 12.80 (s, 1H), 8.40 (d, J = 2.2 Hz, 1H), 7.81 (d, J = 8.6 Hz, 1H), 7.71 (d, J = 8.6 Hz, 2H), 7.65-7.60 (m, 2H), 7.51 (d, J = 8.9 Hz, 2H), 4.11-3.97 (m, 1H), 3.27-3.20 (m, 1H), 3.13-3.05 (m,

1H), 2.92-2.84 (m, 1H), 2.55 (p, J = 7.3 Hz, 1H), 2.48 (dt, J = 3.7, 1.9 Hz, 2H), 1.79-1.70 (m, 1H), 1.47 (dq, J = 12.3, 8.5 Hz, 1H). $^{13}$C NMR (126 MHz, DMSO-$d_6$) δ 169.55, 162.58, 149.76, 139.26, 138.78, 135.30, 133.57, 132.90, 130.30, 129.90, 129.30, 127.85, 122.35, 121.80, 114.80, 114.57, 58.26, 50.12, 45.15, 27.95. HRMS (ESI) m/z [M + H]$^+$ calcd for $C_{23}H_{22}Br_2N_3O_5S$ 609.9641; found 609.9672.

N-(4-bromophenyl)-1-[1-(4-bromophenylsulfonyl)pyrrolidin-2-yl]-3-hydroxy-4-oxo-1,4-dihydropyridine-2-carboxamide (**VNI-5162**). Off-white solid; yield 58%. $^1$H NMR (500 MHz, DMSO-$d_6$) δ 12.84 (s, 1H), 7.85 (d, J = 8.5 Hz, 2H), 7.64 (t, J = 8.0 Hz, 3H), 7.54 (d, J = 8.9 Hz, 1H), 3.99 (d, J = 7.5 Hz, 2H), 3.64 (d, J = 11.7 Hz, 3H), 2.22 (t, J = 10.9 Hz, 1H), 1.53 (d, J = 10.8 Hz, 2H), 1.32-1.21 (m, 1H). $^{13}$C NMR (126 MHz, DMSO-$d_6$) δ 169.94, 163.10, 149.64, 140.35, 138.48, 135.30, 132.93, 132.36, 129.84, 127.56, 123.58, 121.92, 115.53, 115.07, 61.71, 45.88, 40.51, 35.91, 28.25. HRMS (ESI) m/z [M + H]$^+$ calcd for $C_{24}H_{24}Br_2N_3O_5S$ 624.9835; found 624.9811.

N-(4-bromophenyl)-1-[1-(4-methylphenylsulfonyl)pyrrolidin-2-yl]-3-hydroxy-4-oxo-1,4-dihydropyridine-2-carboxamide (**VNI-5163**). Yellow solid; yield 66%. $^1$H NMR (500 MHz, DMSO-$d_6$) δ 12.77 (s, 1H), 8.31 (s, 1H), 7.57-7.35 (m, 8H), 3.91 (m, 1H), 3.57 (m, 1H), 2.33 (s, 3H), 2.11-2.03 (m, 1H), 1.72 (m, 1H), 1.45 (m, 1H), 1.23-1.15 (m, 4H). $^{13}$C NMR (126 MHz, DMSO-$d_6$) δ 169.93, 163.10, 149.63, 143.92, 140.35, 138.49, 133.03, 132.35, 130.24, 129.06, 127.90, 123.58, 121.91, 115.52, 115.06, 61.78, 55.38, 45.96, 36.03, 28.24, 21.46. HRMS (ESI) m/z [M + H]$^+$ calcd for $C_{25}H_{27}BrN_3O_5S$ 561.0928; found 561.0948.

N-(4-bromophenyl)-1-[1-(4-fluorophenyl)sulfonylpyrrolidin-2-yl]-3-hydroxy-4-oxo-1,4-dihydropyridine-2-carboxamide (**VNI-5164**). Yellow solid; yield 57%. $^1$H NMR (500 MHz, $CDCl_3$) δ 12.03 (s, 1H), 8.32 (d, J = 2.3 Hz, 1H), 7.79-7.73 (m, 2H), 7.61 (d, J = 8.5 Hz, 2H), 7.46 (d, J = 8.4 Hz, 2H), 7.22-7.19 (m, 2H), 3.85 (dd, J = 20.6, 9.6 Hz, 4H), 2.30-2.21 (m, 2H), 1.68 (d, J = 13.1 Hz, 2H), 1.45 (tt, J = 13.9, 7.0 Hz, 2H). $^{13}$C NMR (126 MHz, DMSO-$d_6$) δ 169.45, 162.61,

149.15, 139.86, 138.00, 131.86, 130.46, 130.38, 123.10, 121.43, 116.62, 116.44, 61.26, 45.43, 35.49, 27.73. HRMS (ESI) m/z $[M + H]^+$ calcd for $C_{24}H_{24}BrFN_3O_5S$ 564.0605; found 564.0642.

N-(4-bromophenyl)-1-[1-(2-fluorophenyl)sulfonylpyrrolidin-2-yl]-3-hydroxy-4-oxo-1,4-dihydropyridine-2-carboxamide (**VNI-5165**). Yellow solid; yield 58%. $^1$H NMR (500 MHz, $CDCl_3$) δ 12.03 (s, 1H), 8.34 (d, J = 2.2 Hz, 1H), 7.84 (t, J = 6.9 Hz, 1H), 7.60 (dd, J = 14.0, 7.5 Hz, 3H), 7.49-7.42 (m, 2H), 7.24-7.19 (m, 2H), 3.97 (d, J = 12.7 Hz, 2H), 3.84 (d, J = 7.4 Hz, 2H), 2.56 (t, J = 12.3 Hz, 2H), 1.69 (d, J = 13.2 Hz, 2H), 1.43 (tt, J = 14.2, 7.1 Hz, 2H). $^{13}$C NMR (126 MHz, DMSO-$d_6$) δ 169.46, 162.62, 149.14, 139.88, 138.00, 131.86, 130.73, 125.09, 123.14, 121.44, 117.62, 117.44, 115.04, 114.59, 61.24, 45.00, 35.55, 27.95. HRMS (ESI) m/z $[M + H]^+$ calcd for $C_{24}H_{24}BrFN_3O_5S$ 564.0605; found 564.0652.

N-(4-bromophenyl)-1-[1-(2-nitrophenyl)sulfonylpyrrolidin-2-yl]-3-hydroxy-4-oxo-1,4-dihydropyridine-2-carboxamide (**VNI-5166**). Yellow solid; yield 55%. $^1$H NMR (500 MHz, DMSO-$d_6$) δ 12.85 (s, 1H), 8.41 (d, J = 2.2 Hz, 1H), 7.97 (ddd, J = 7.2, 5.4, 1.4 Hz, 2H), 7.86 (dtd, J = 26.1, 7.6, 1.4 Hz, 2H), 7.67-.59 (m, 3H), 7.53 (d, J = 8.5 Hz, 2H), 4.02 (d, J = 7.3 Hz, 2H), 2.70 (td, J = 12.4, 2.5 Hz, 2H), 1.95 (m, 1H), 1.61-1.54 (m, 2H), 1.25 (qd, J = 12.4, 4.1 Hz, 2H). $^{13}$C NMR (126 MHz, DMSO-$d_6$) δ 169.47, 162.62, 149.16, 147.78, 139.89, 138.00, 134.63, 132.26, 131.85, 130.17, 129.80, 124.14, 123.14, 121.44, 115.04, 61.19, 45.15, 35.54, 28.03. HRMS (ESI) m/z $[M + H]^+$ calcd for $C_{24}H_{24}BrN_4O_7S$ 593.0477; found 593.0502.

N-(4-bromophenyl)-1-[1-(4-methoxyphenyl)sulfonylpyrrolidin-2-yl]-3-hydroxy-4-oxo-1,4-dihydropyridine-2-carboxamide (**VNI-5167**). Off-white solid; yield 62%. $^1$H NMR (500 MHz, DMSO-$d_6$) δ 12.82 (s, 1H), 8.36 (s, 1H), 7.61 (m, 4H), 7.52 (d, J = 8.4 Hz, 2H), 7.11 (d, J = 8.4 Hz, 2H), 3.96 (d, J = 7.2 Hz, 2H), 3.82 (s, 3H), 3.60 (d, J = 11.4 Hz, 2H), 2.11 (t, J = 11.7 Hz, 2H), 1.50 (d, J = 12.7 Hz, 2H), 1.24 (d, J = 13.2 Hz, 2H). $^{13}$C NMR (126 MHz, DMSO-$d_6$) δ 169.94, 163.12, 149.64, 140.35, 138.49, 132.35, 130.08, 127.42, 123.58, 121.91, 115.53, 115.07, 114.94, 61.80, 56.17, 45.98, 36.07, 28.24. HRMS (ESI) m/z $[M + H]^+$ calcd for $C_{25}H_{27}BrN_3O_6S$ 576.0798; found 576.0800.

N-(4-bromophenyl)-1-{[1-((2,3-dichlorophenyl)sulfonyl)piperidin-4-yl]methyl}-3-hydroxy-4-oxo-1,4-dihydropyridine-2-carboxamide (**VNI-5168**). Off-white solid; yield 64%. $^1$H NMR (500 MHz, DMSO-$d_6$) δ 12.83 (s, 1H), 8.41 (d, J = 2.2 Hz, 1H), 7.94 (dd, J = 8.1, 4.1 Hz, 2H), 7.67-7.50 (m, 7H), 4.01 (d, J = 7.3 Hz, 2H), 3.71 (d, J = 12.5 Hz, 2H), 2.78 (t, J = 12.2 Hz, 2H), 1.53 (d, J = 12.7 Hz, 2H), 1.27-1.15 (m, 2H). $^{13}$C NMR (126 MHz, DMSO-$d_6$) δ 169.98, 163.14, 149.68, 140.42, 138.99, 138.52, 135.25, 135.08, 132.38, 130.60, 129.62, 129.42, 129.08, 123.68, 121.95, 115.55, 115.13, 61.72, 45.41, 40.54, 36.28, 28.81. HRMS (ESI) m/z $[M + H]^+$ calcd for $C_{24}H_{23}BrCl_2N_3O_5S$ 613.9898; found 613.9943.

N-(4-bromophenyl)-1-[1-(isopropylsulfonyl)piperidin-4-yl]-3-hydroxy-4-oxo-1,4-dihydropyridine-2-carboxamide (**VNI-5170**). Yellow solid; yield 52%. $^1$H NMR (500 MHz, DMSO-$d_6$) δ 12.88 (s, 1H), 8.47 (d, J = 2.2 Hz, 1H), 7.72 (d, J = 2.1 Hz, 1H), 7.66 (d, J = 8.5 Hz, 2H), 7.56 (d, J = 8.6 Hz, 2H), 4.06 (d, J = 7.5 Hz, 1H), 3.67 (d, J = 12.6 Hz, 1H), 3.30 (q, J = 6.8 Hz, 1H), 2.89-2.81 (m, 2H), 2.00 (s, 1H), 1.54 (d, J = 13.0 Hz, 1H), 1.22 (d, J = 6.8 Hz, 6H). $^{13}$C NMR (126 MHz, DMSO-$d_6$) δ 169.96, 163.14, 149.67, 140.40, 138.51, 132.36, 123.69, 121.94, 115.54, 115.10, 61.90, 52.32, 45.59, 36.56, 29.27, 16.90. HRMS (ESI) m/z $[M + H]^+$ calcd for $C_{21}H_{27}BrN_3O_5S$ 512.0879; found 512.0854.

N-(4-fluorophenyl)-1-[1-(4-methylphenylsulfonyl)piperidin-4-yl]-3-hydroxy-4-oxo-1,4-dihydropyridine-2-carboxamide (**VNI-5171**). Yellow solid; yield 60%. $^1$H NMR (500 MHz, DMSO-$d_6$) δ 12.75 (s, 1H), 8.38 (d, J = 2.2 Hz, 1H), 7.67 (dd, J = 8.9, 5.1 Hz, 2H), 7.62-7.57 (m, 3H), 7.42 (d, J = 8.1 Hz, 2H), 7.24-7.15 (m, 2H), 3.98 (d, J = 7.4 Hz, 2H), 3.63 (d, J = 11.6 Hz, 2H), 2.30 (s, 3H), 2.13 (td, J = 12.1, 2.5 Hz, 2H), 1.55-1.45 (m, 2H), 1.25 (qd, J = 12.4, 4.3 Hz, 2H). $^{13}$C NMR (126 MHz, DMSO-$d_6$) δ 169.92, 162.89, 149.59, 143.92, 140.29, 135.58, 133.07, 130.25, 127.90, 123.52, 121.72, 116.22, 116.04, 61.77, 45.96, 36.04, 28.26, 21.46. HRMS (ESI) m/z $[M + H]^+$ calcd for $C_{25}H_{27}FN_3O_5S$ 500.1650; found 500.1689.

N-(4-chlorophenyl)-1-[1-(4-methylphenylsulfonyl)piperidin-4-yl]-3-hydroxy-4-oxo-1,4-dihydropyridine-2-carboxamide (**VNI-5172**). Yellow solid; yield 53%. $^1$H NMR (500 MHz, $CDCl_3$) δ

11.99 (s, 1H), 8.31 (d, J = 6.8 Hz, 1H), 7.68-7.60 (m, 2H), 7.32 (dd, J = 8.3, 5.6 Hz, 2H), 7.19 (d, J = 3.6 Hz, 1H), 3.86 (d, J = 12.3 Hz, 1H), 3.81 (d, J = 7.3 Hz, 1H), 2.43 (s, 3H), 2.27-2.19 (m, 1H), 1.76 (d, J = 3.8 Hz, 1H), 1.68 (s, 1H), 1.44 (qd, J = 12.4, 4.2 Hz, 1H). $^{13}$C NMR (126 MHz, DMSO-$d_6$) δ 169.61, 162.75, 149.29, 143.58, 140.01, 137.75, 132.72, 129.91, 129.11, 127.56, 127.19, 123.24, 121.20, 114.72, 61.45, 45.63, 35.69, 27.91, 21.13. HRMS (ESI) m/z [M + H]$^+$ calcd for $C_{25}H_{27}ClN_3O_5S$ 516.1354; found 516.1383.

N-(benzothiophen-2-yl)-1-[1-(4-methylphenylsulfonyl)piperidin-4-yl]-3-hydroxy-4-oxo-1,4-dihydropyridine-2-carboxamide (**VNI-5173**). Yellow solid; yield 46%. $^1$H NMR (500 MHz, $CDCl_3$) δ 12.7 (s, 1H), 8.52 (s, 1H), 8.05 (ddd, J = 8.1, 1.5, 0.4 Hz, 1H), 7.85-7.25 (m, 11H), 3.88 (d, J = 3.2 Hz, 2H), 3.59 (ddd, J = 14.8, 2.8, 2.8 Hz, 2H), 3.22 (ddd, J = 14.8, 6.8, 6.8 Hz, 2H), 2.32 (s, 3H), 2.12 (m, 1H), 1.79-1.61 (m, 4H). $^{13}$C NMR (126 MHz, $CDCl_3$) δ 184.1, 163.8, 147.4, 146.0, 144.9, 142.0, 139.6, 136.8, 135.9, 135.7, 134.9, 132.1, 129.4, 127.7, 127.0, 124.5, 122.9, 121.9, 121.2, 120.4, 118.1, 111.3, 62.3, 44.5, 35.0, 31.7, 21.4. HRMS (ESI) m/z [M + H]$^+$ calcd for $C_{31}H_{30}N_3O_5S_2$ 588.1508; found 588.1542.

N-(dibenzofuran-2-yl)-1-[1-(4-methylphenylsulfonyl)piperidin-4-yl]-3-hydroxy-4-oxo-1,4-dihydropyridine-2-carboxamide (**VNI-5174**). Yellow solid; yield 42%. $^1$H NMR (500 MHz, $CDCl_3$) δ 12.7 (s, 1H), 8.52 (s, 1H), 8.05 (ddd, J = 8.1, 1.5, 0.4 Hz, 1H), 7.95-7.15 (m, 11H), 3.89 (d, J = 3.2 Hz, 2H), 3.59 (ddd, J = 14.8, 2.8, 2.8 Hz, 2H), 3.22 (ddd, J = 14.8, 6.8, 6.8 Hz, 2H), 2.32 (s, 3H), 2.09 (m, 1H), 1.79-1.61 (m, 4H). $^{13}$C NMR (126 MHz, $CDCl_3$) δ 184.1, 163.8, 156.1, 155.2, 147.4, 146.0, 144.9, 142.0, 136.8, 132.1, 129.4, 127.7, 127.5, 124.3, 124.2, 122.6, 122.4, 121.2, 120.4, 118.1, 111.6, 110.3, 62.3, 44.5, 35.0, 31.7, 21.4. HRMS (ESI) m/z [M + H]$^+$ calcd for $C_{31}H_{30}N_3O_6S$ 572.1845; found 572.1848.

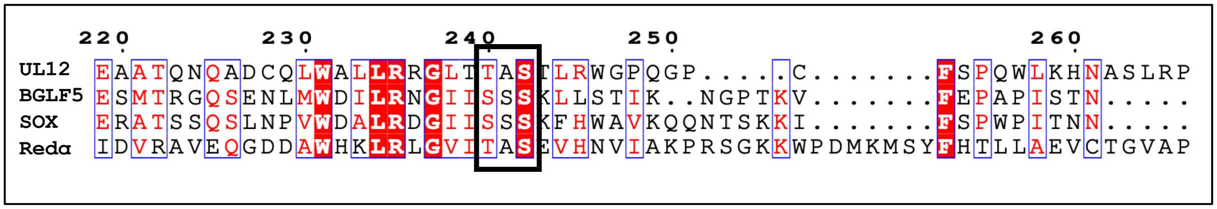


**Figure S1:** Sequence alignment of UL12, BGLF5, SOX, and Redα showing the conserved motif Thr-Ala-Ser in motif I of UL12 and λ exonuclease Redα and the Ser-Ser-Ser motif in the γ-herpesvirus nucleases BGLF5 and SOX (black box).

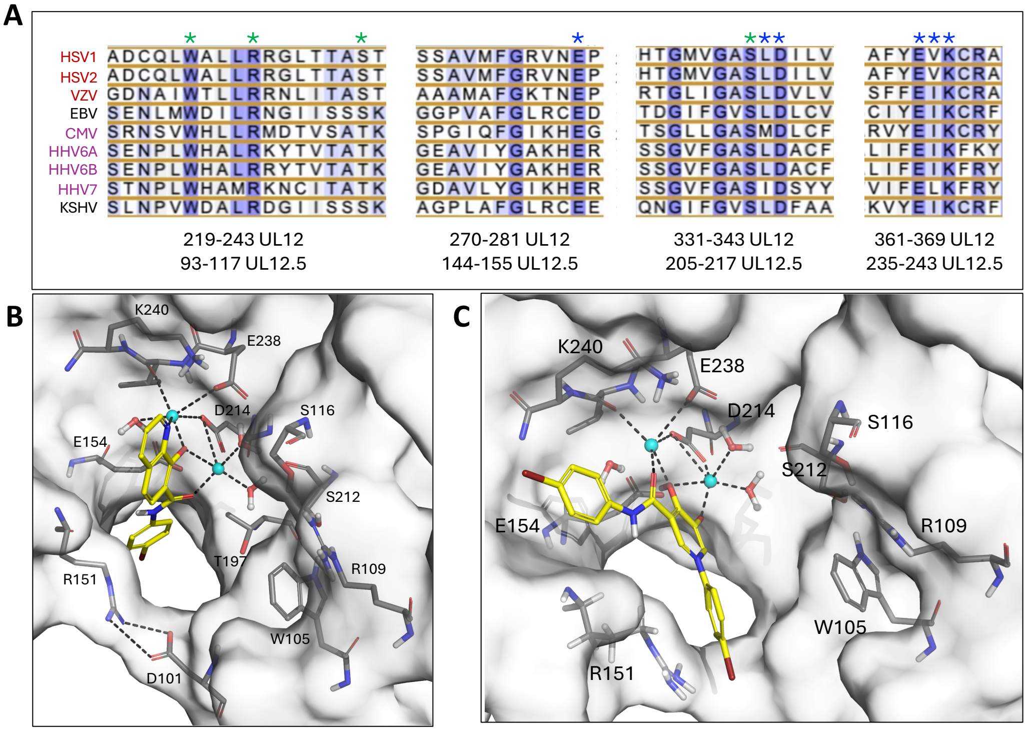


| Compound | $IC_{50}$ (UL12) | $EC_{50}$ HSV | $EC_{50}$ CMV | $CC_{50}$ in HFF | SI (HSV) | SI (CMV) |
|---|---|---|---|---|---|---|
| AK-157 | 1.3 μM | 0.170 μM | 0.183 μM | 7 μM | 41 | 38.3 |
| DD-I-38 | 3.1 μM | 1.94 μM | 1.81 μM | >50 | >25.8 | >27.6 |

**Figure S2: Conserved UL12 active-site features and predicted inhibitor binding modes.** (A) Multiple sequence alignment highlighting conserved, identical, and essential catalytic active-site residues (blue stars) and the 5′-phosphate binding pocket defined by the WRSS motif (green stars). (B) Predicted binding site of the HQ inhibitor AK-157 within the UL12 active site, showing its spatial separation from the WRSS pocket. (C) Predicted binding site of the HP inhibitor DD-I-38 within the UL12 active site, similarly positioned distal to the WRSS pocket. In (B) and (C), inhibitors are shown as yellow sticks, UL12 active-site residues as dark gray sticks, coordinating water molecules as red spheres, and $Mg^{2+}$ ions as cyan spheres. (D)

Summary of enzymatic inhibition ($IC_{50}$), cytotoxicity ($CC_{50}$) in human foreskin fibroblast (HFF) cells, antiviral activity ($EC_{50}$), and calculated selectivity indices (SI) for HSV and CMV.

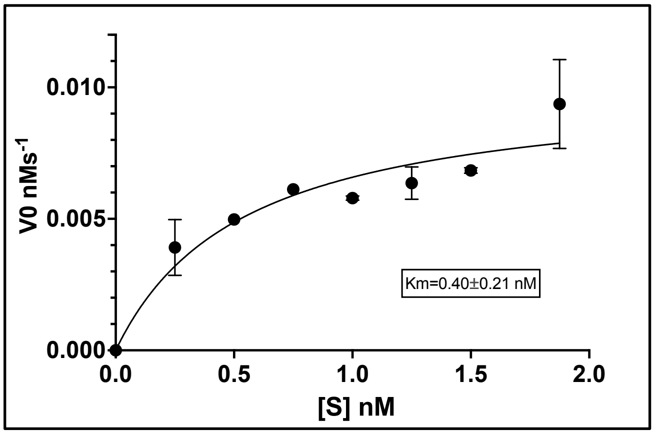


**Figure S3:** Michaelis–Menten plot showing initial reaction velocity ($v_0$) as a function of substrate concentration for UL12. Experiments were performed four times, and data points represent the mean ± SEM from two independent experiments. The solid line represents a nonlinear least-squares fit to the Michaelis–Menten equation using GraphPad Prism, from which the apparent $K_m$ value was determined. The $K_m$ value and corresponding $IC_{50}$ values were used to extrapolate the $K_i$ using the Cheng–Prusoff equation.

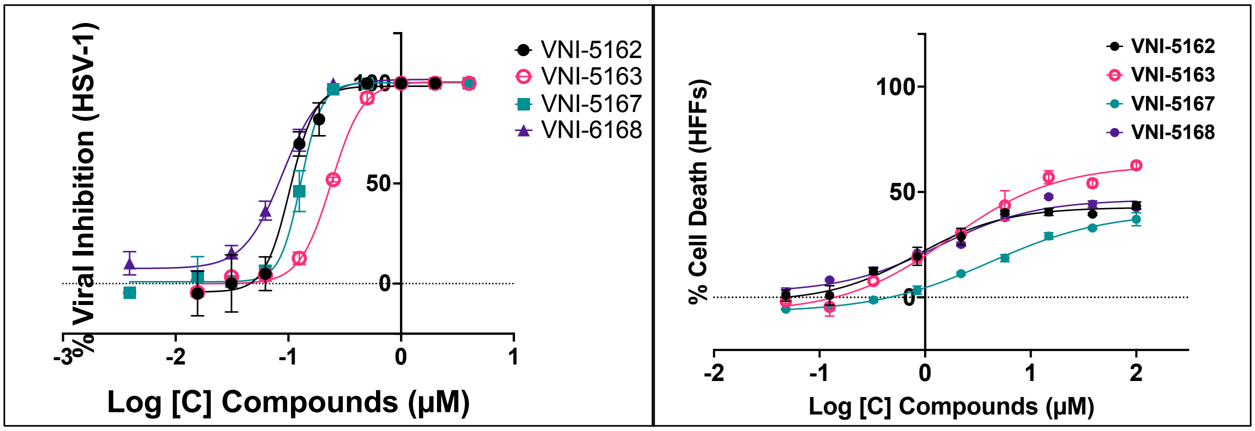


**Figure S4:** Dose response used to determine A) EC50 values for HSV-1 using yield assay in HFF cells infected with fluorescently tagged HSV-1 virus. B) cytotoxicity on HFF cells at 65 h post-treatment was measured using Promega Cell Titer-Glo.

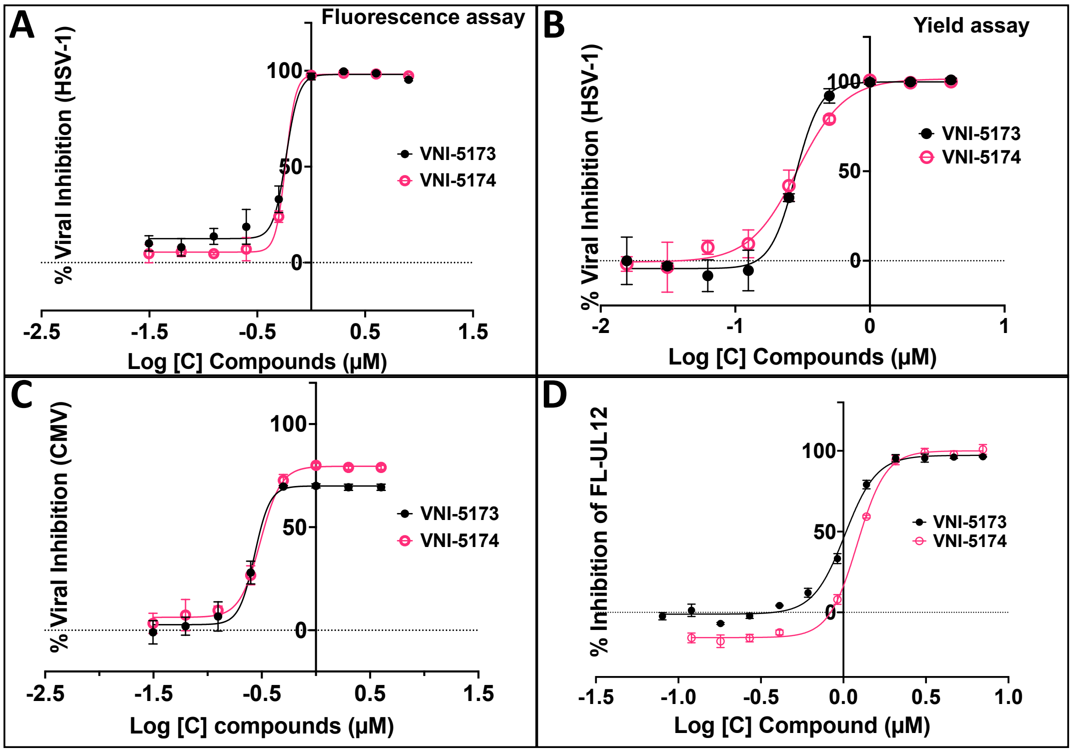


**Figure S5: Biochemical and antiviral characterization of analogs bearing a dibenzothiophene (VNI-5173) and dibenzofuran (VNI-5174) sidechain.**

Dose-response analyses for antiviral activity to determine $EC_{50}$ values for HSV-1 (A and B) and HCMV (C) in HFF cells infected with fluorescently tagged viruses in the presence of increasing concentration of the compound. Antiviral activity was evaluated using either high-content image-based assay or yield assay. (D) Dose-response analyses for enzymatic inhibition to determine $IC_{50}$ values for full-length (FL)-UL12 using the PicoGreen-based nuclease assay.

*Percentage inhibition was calculated relative to the uninhibited DMSO control. $IC_{50}$ and $EC_{50}$ values were obtained by nonlinear regression analysis in GraphPad Prism using a four-parameter logistic model (log[inhibitor] versus response, variable slope). Data represent the mean of three independent experiments, with each experiment comprising 2-3 technical replicates.*

**Table S1**: Data collection, refinement, and validation statistics for HSV-1 UL12.5 (PDB ID: 10HO). Highest-resolution shell values are shown in parentheses.

| | |
|---|---|
| Space group | P 21 21 21 |
| Unit cell (Å, °) | a = 59.00, b = 80.23, c = 127.57; α = β = γ = 90° |
| **Data collection statistics** | |
| Resolution range (Å) | 29.64–2.46 (2.56–2.46) |
| Unique reflections | 22,700 (2508) |
| Completeness (%) | 99.9 (99.2) |
| Multiplicity | 1.9 |
| Mean I/σ(I) | 12.4 (2.4) |
| $CC_{1/2}$ | 0.999 (0.885) |
| Rmerge | 0.037 |
| Rpim / Rmeas | 0.037 / 0.052 (0.316 / 0.446) |
| **MR phasing statistics** | |
| Top LLG | 317 |
| Top TFZ | 19.12 |
| **Refinement statistics** | |
| Reflections (work / free) | 21,547 / 1,104 (4.85%) |
| $R_{work}$ / $R_{free}$ | 0.181 / 0.238 |
| $CC_{work}$ | 0.9616 |
| $CC_{free}$ | 0.9342 |
| Number of non-hydrogen atoms | 3400 |
| Macromolecules | 3317 |
| Ligands | 18 |
| Solvents | 65 |
| Average B-factor ($Å^2$) | 62.0 |
| Wilson B | 49.4 |
| Macromolecules | 62.68 |
| Ligands | 79.26 |
| Water | 47.73 |
| RMSD bond lengths | 0.0107 Å |
| RMSD bond angles | 2.239° |
| **Ramachandran plot** | |
| Ramachandran favored | 95% |
| Ramachandran allowed | 5% |
| Ramachandran outliers | 0% |
| Clashscore | 3 |

### Mass spectra for compounds

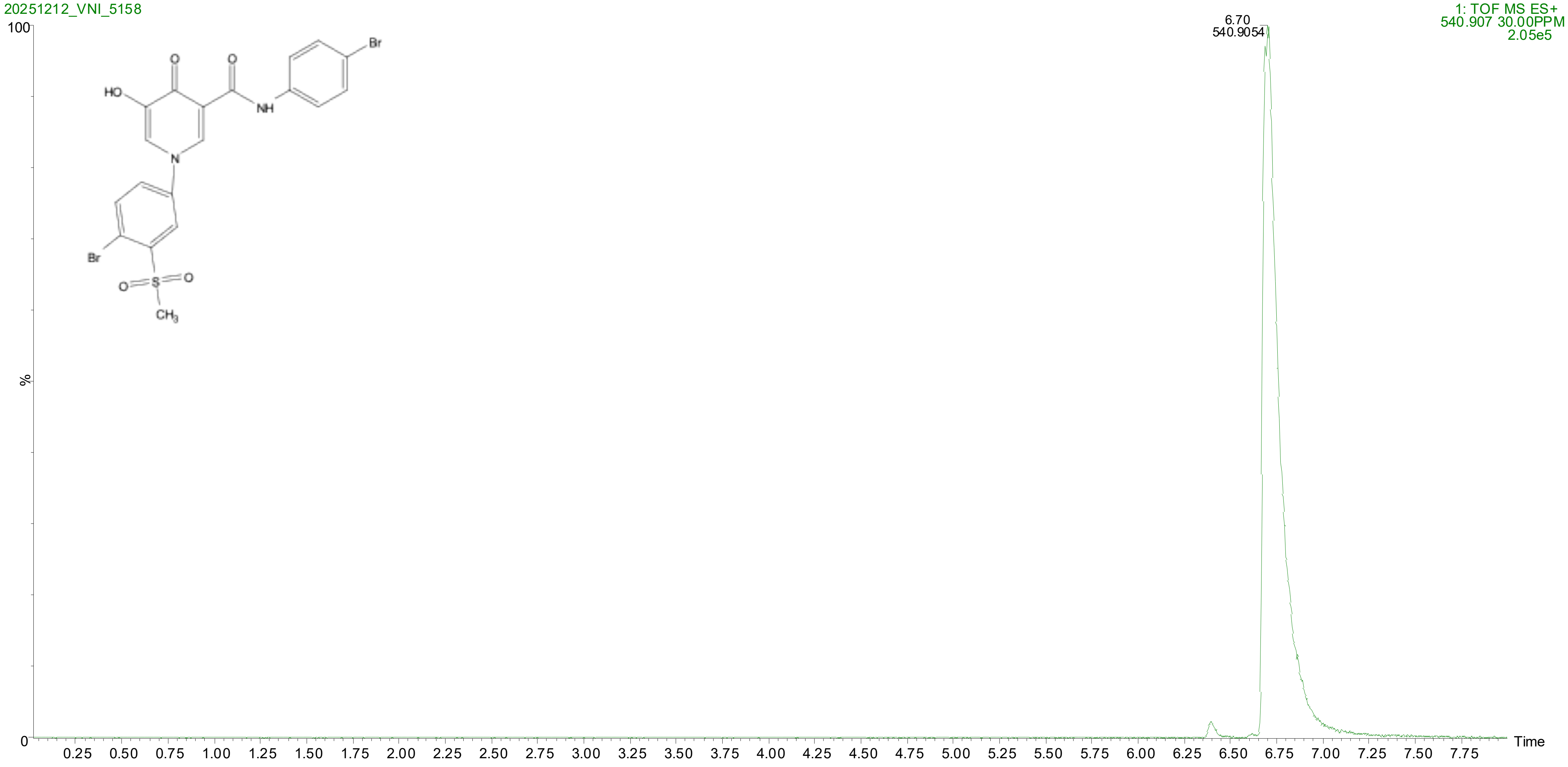
20251212_VNI_5158
1: TOF MS ES+
540.907 30.00PPM
2.05e5
6.70
540.9054
100
%
0
Time
0.25 0.50 0.75 1.00 1.25 1.50 1.75 2.00 2.25 2.50 2.75 3.00 3.25 3.50 3.75 4.00 4.25 4.50 4.75 5.00 5.25 5.50 5.75 6.00 6.25 6.50 6.75 7.00 7.25 7.50 7.75
HO
O
O
NH
Br
N
Br
O
S
O
CH3

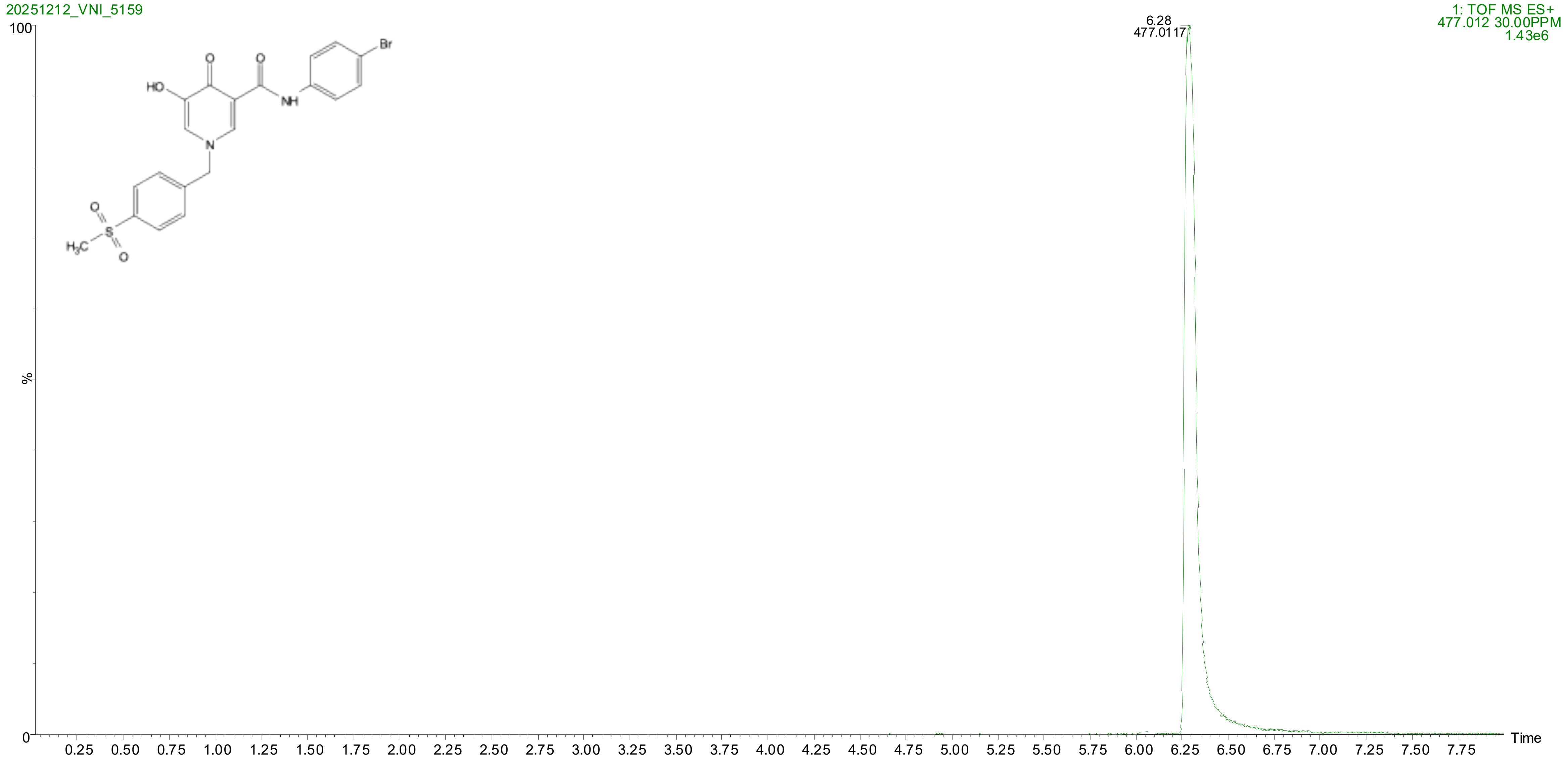

20251212_VNI_5159
1: TOF MS ES+
477.012 30.00PPM
1.43e6
6.28
477.0117
100
%
0
HO
O
O
NH
Br
N
O
S
O
H3C
0.25 0.50 0.75 1.00 1.25 1.50 1.75 2.00 2.25 2.50 2.75 3.00 3.25 3.50 3.75 4.00 4.25 4.50 4.75 5.00 5.25 5.50 5.75 6.00 6.25 6.50 6.75 7.00 7.25 7.50 7.75
Time

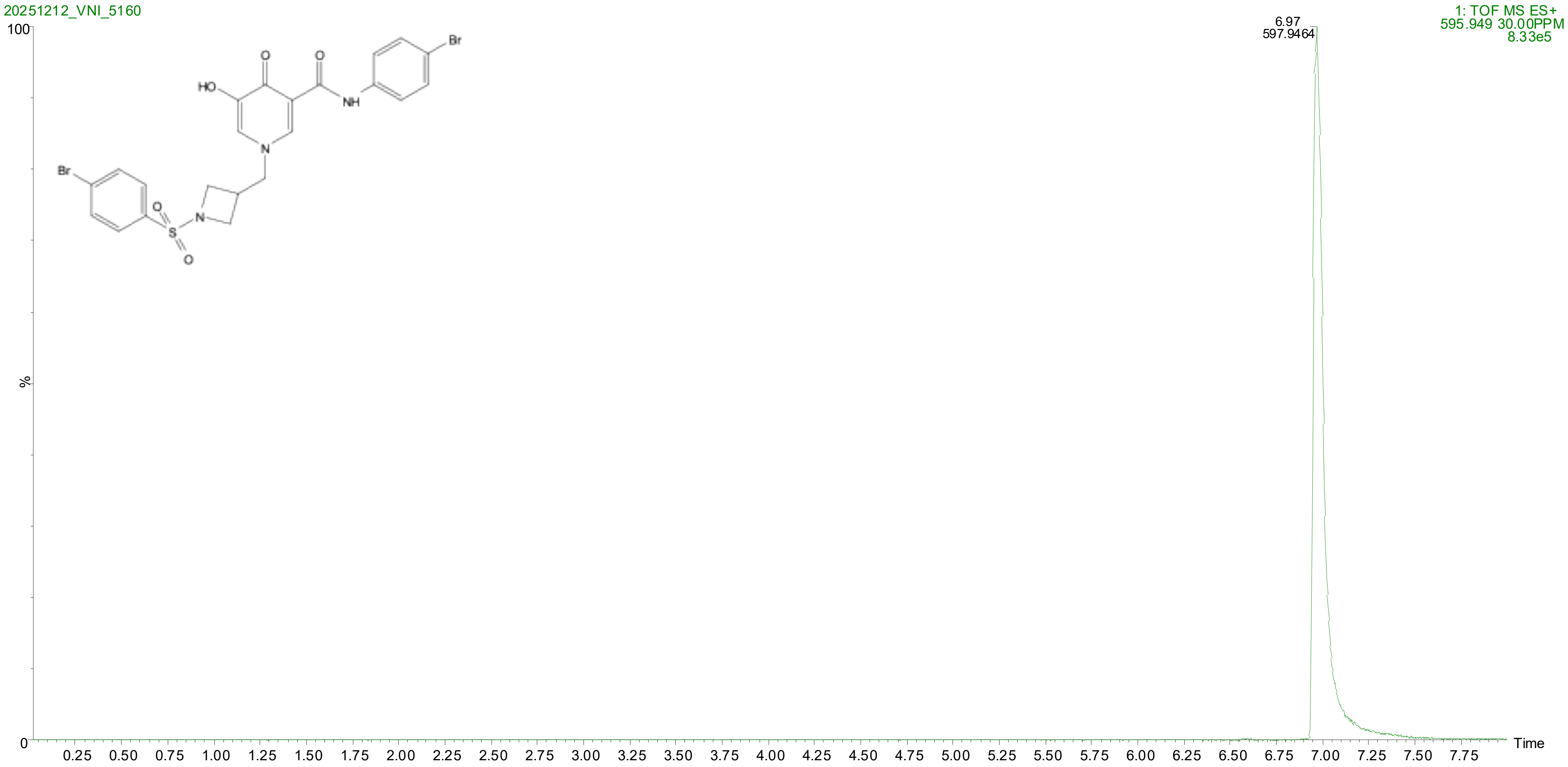
20251212_VNI_5160
1: TOF MS ES+
595.949 30.00PPM
8.33e5
6.97
597.9464
100
%
0
Time
0.25 0.50 0.75 1.00 1.25 1.50 1.75 2.00 2.25 2.50 2.75 3.00 3.25 3.50 3.75 4.00 4.25 4.50 4.75 5.00 5.25 5.50 5.75 6.00 6.25 6.50 6.75 7.00 7.25 7.50 7.75
HO
O
O
NH
Br
N
Br
O
S
O
N

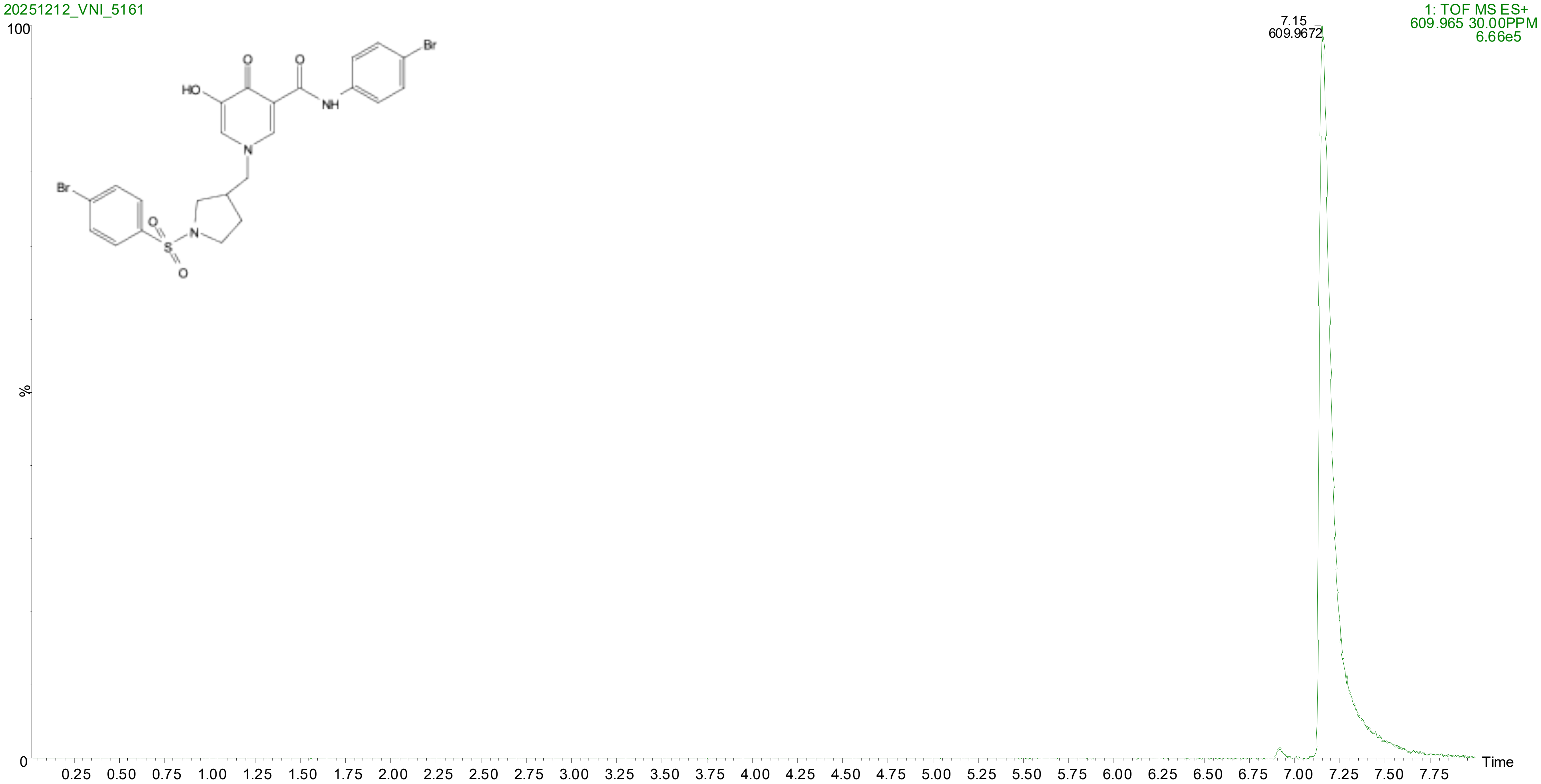
20251212_VNI_5161
1: TOF MS ES+
609.965 30.00PPM
6.66e5
7.15
609.9672
100
%
0
Time
0.25 0.50 0.75 1.00 1.25 1.50 1.75 2.00 2.25 2.50 2.75 3.00 3.25 3.50 3.75 4.00 4.25 4.50 4.75 5.00 5.25 5.50 5.75 6.00 6.25 6.50 6.75 7.00 7.25 7.50 7.75
HO
O
NH
Br
N
S

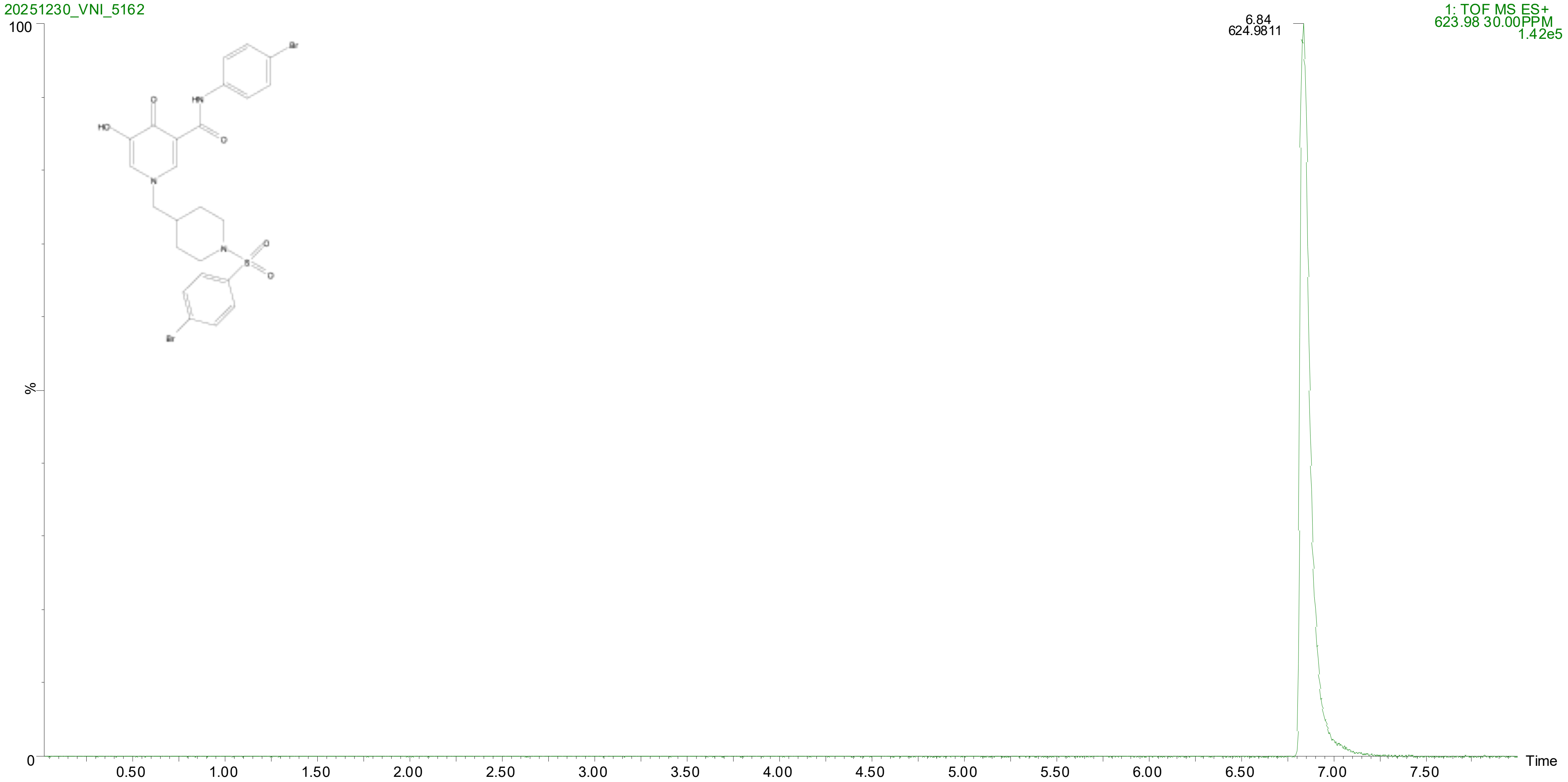
20251230_VNI_5162
1: TOF MS ES+
623.98 30.00PPM
1.42e5
6.84
624.9811
100
%
0
Time
0.50
1.00
1.50
2.00
2.50
3.00
3.50
4.00
4.50
5.00
5.50
6.00
6.50
7.00
7.50

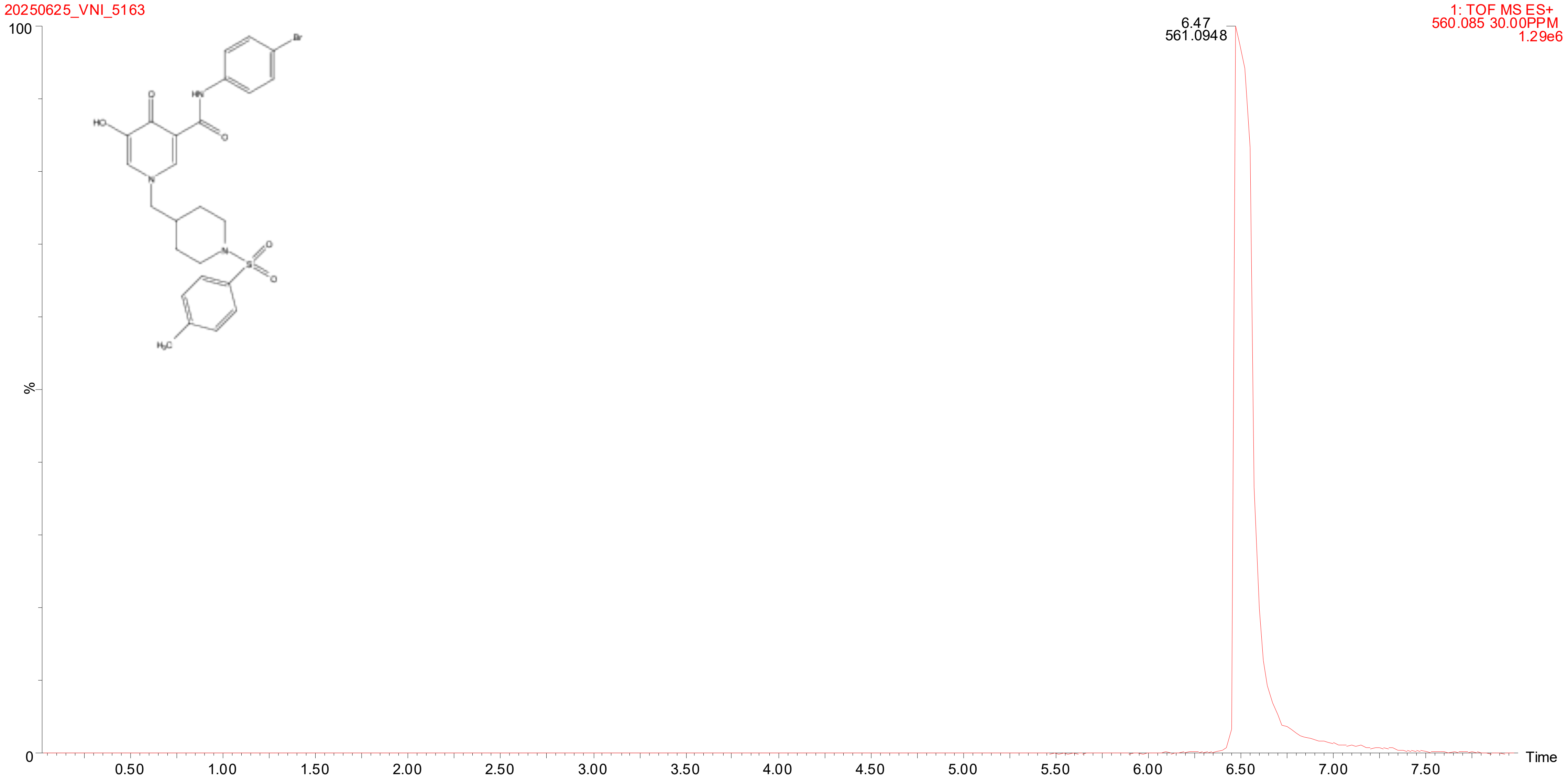
20250625_VNI_5163
1: TOF MS ES+
560.085 30.00PPM
1.29e6
6.47
561.0948
100
%
0
Time
0.50
1.00
1.50
2.00
2.50
3.00
3.50
4.00
4.50
5.00
5.50
6.00
6.50
7.00
7.50
Br
HN
O
HO
O
N
N
S
O
O
H3C

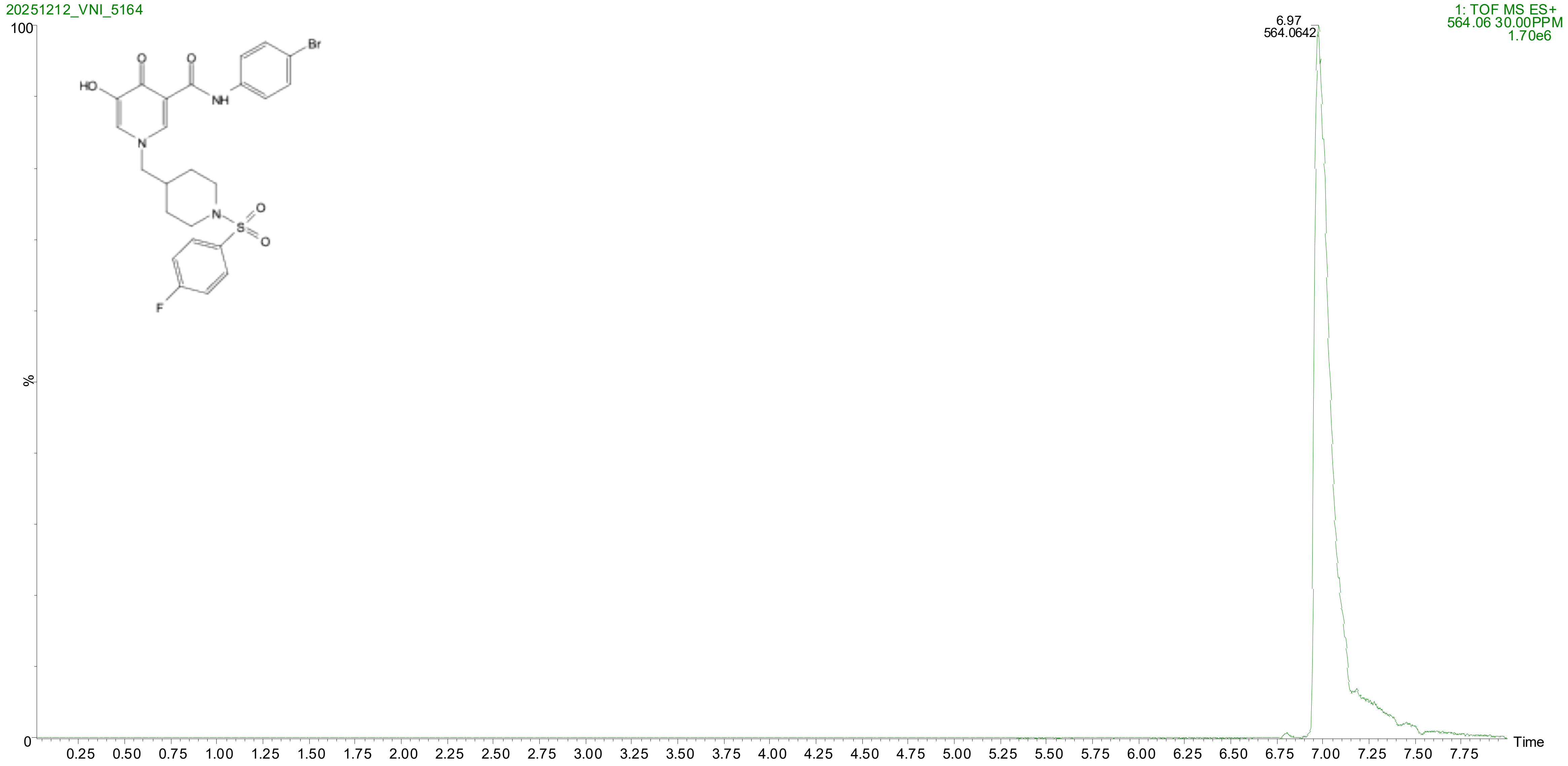
20251212_VNI_5164
1: TOF MS ES+
564.06 30.00PPM
1.70e6
6.97
564.0642
100
%
0
Time
0.25 0.50 0.75 1.00 1.25 1.50 1.75 2.00 2.25 2.50 2.75 3.00 3.25 3.50 3.75 4.00 4.25 4.50 4.75 5.00 5.25 5.50 5.75 6.00 6.25 6.50 6.75 7.00 7.25 7.50 7.75
HO
O
O
NH
Br
N
N
S
O
O
F

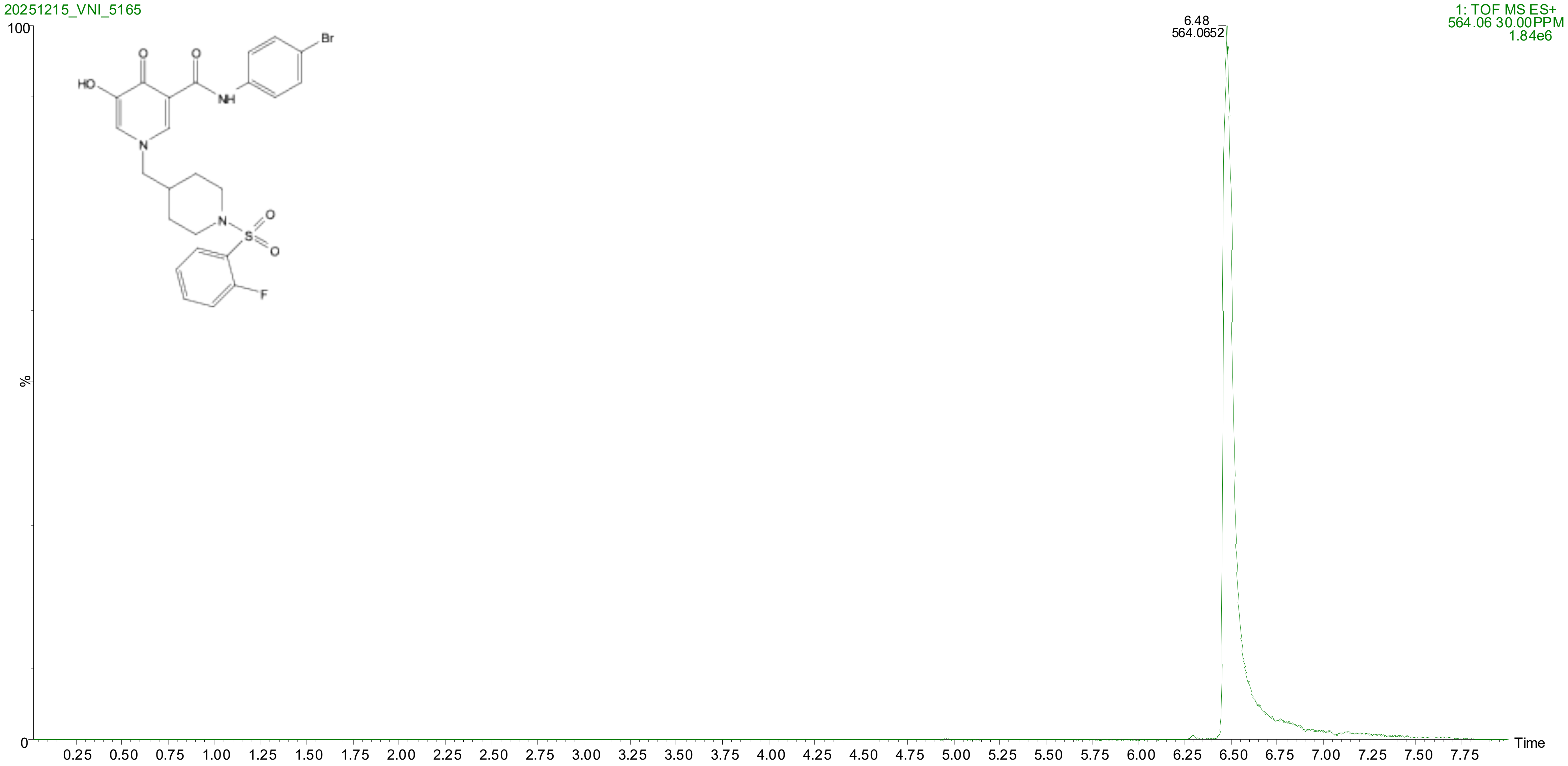
20251215_VNI_5165
1: TOF MS ES+
564.06 30.00PPM
1.84e6
6.48
564.0652
100
%
0
0.25 0.50 0.75 1.00 1.25 1.50 1.75 2.00 2.25 2.50 2.75 3.00 3.25 3.50 3.75 4.00 4.25 4.50 4.75 5.00 5.25 5.50 5.75 6.00 6.25 6.50 6.75 7.00 7.25 7.50 7.75
Time
HO
O
O
NH
Br
N
N
S
O
O
F

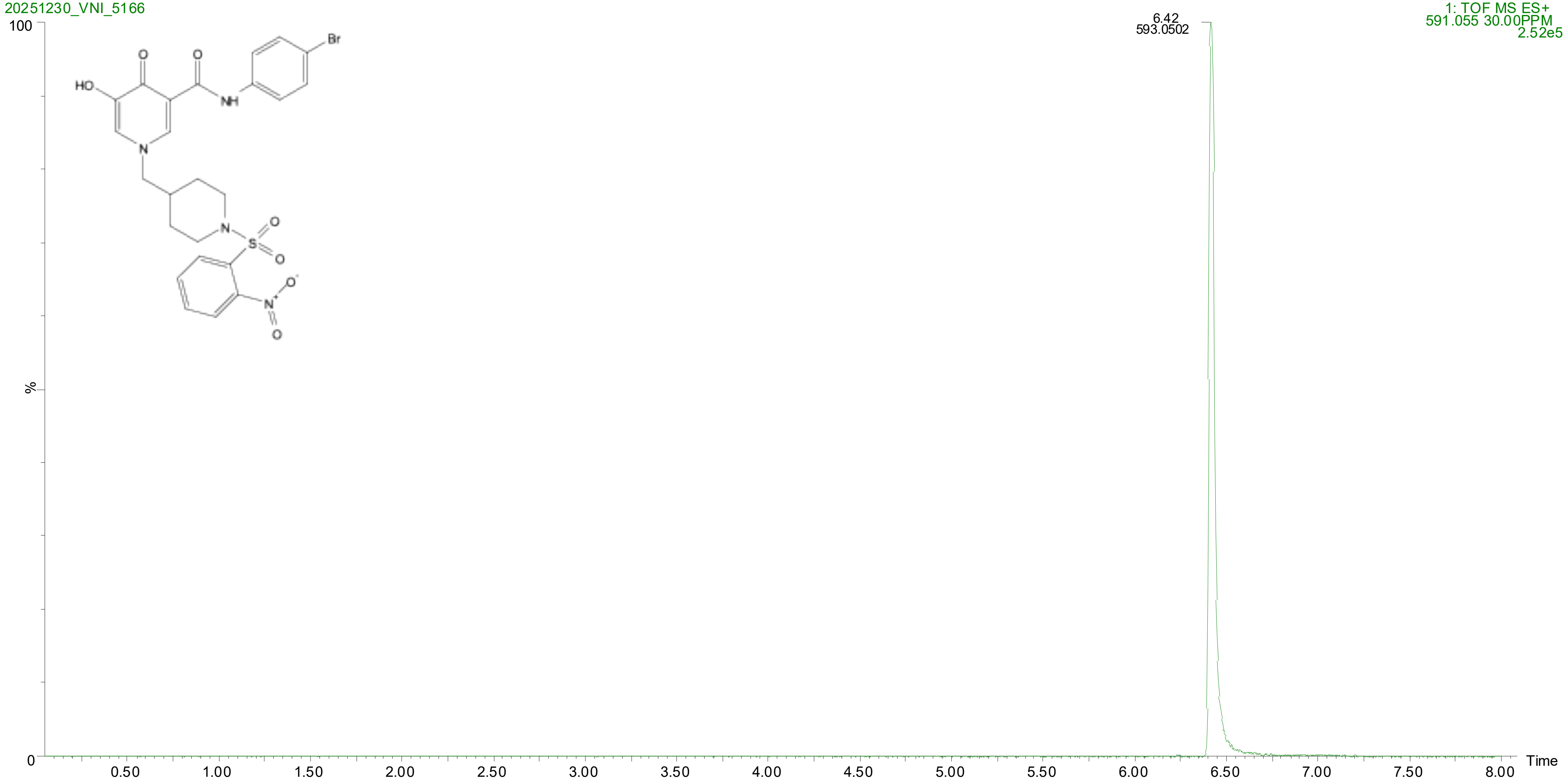

20251230_VNI_5166
1: TOF MS ES+
591.055 30.00PPM
2.52e5
6.42
593.0502
100
%
0
Time
0.50
1.00
1.50
2.00
2.50
3.00
3.50
4.00
4.50
5.00
5.50
6.00
6.50
7.00
7.50
8.00
HO
O
O
NH
Br
N
N
S
O
O
N+
O-
O

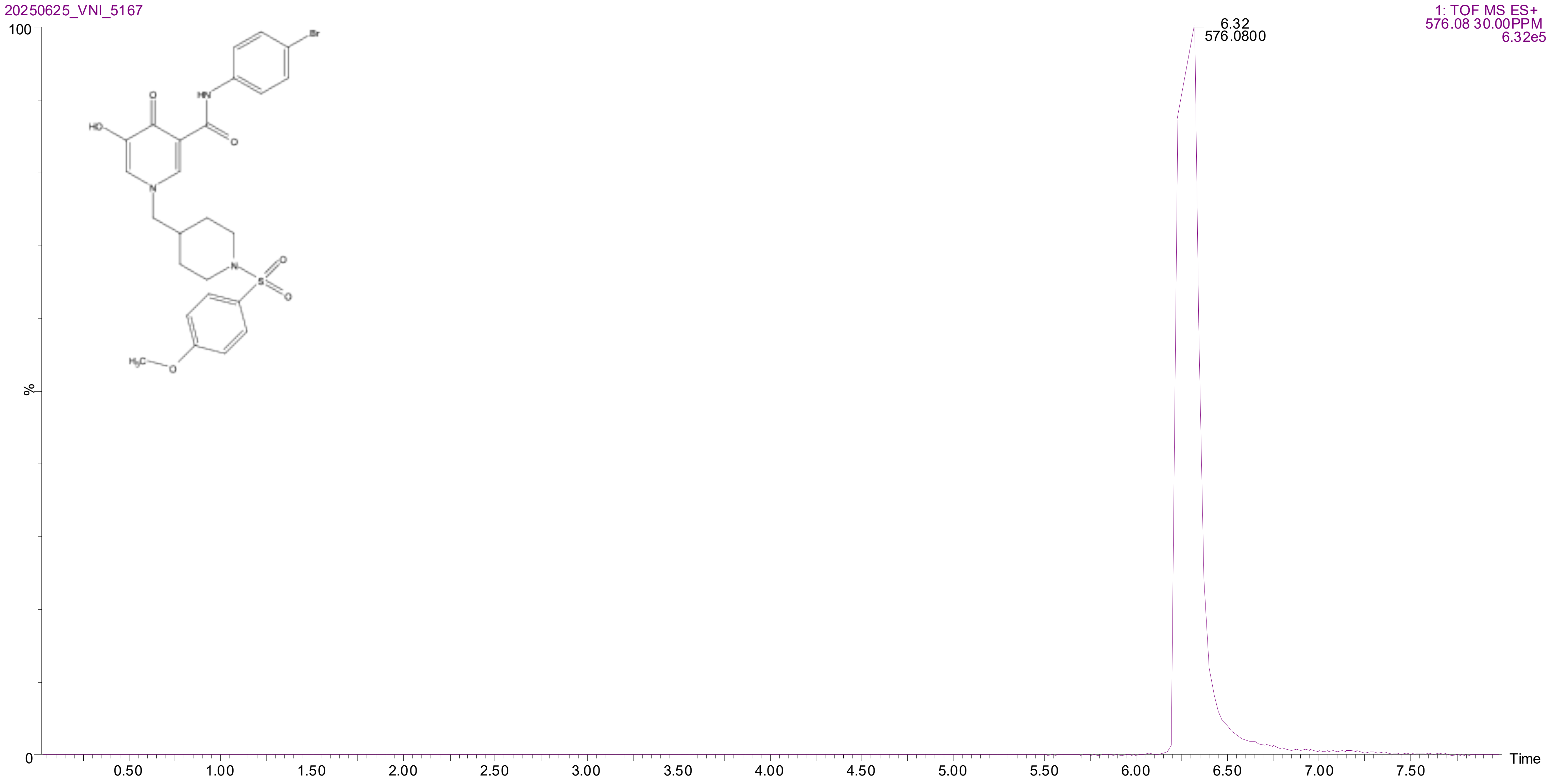
20250625_VNI_5167
1: TOF MS ES+
576.08 30.00PPM
6.32e5
6.32
576.0800
100
%
0
Time
0.50
1.00
1.50
2.00
2.50
3.00
3.50
4.00
4.50
5.00
5.50
6.00
6.50
7.00
7.50

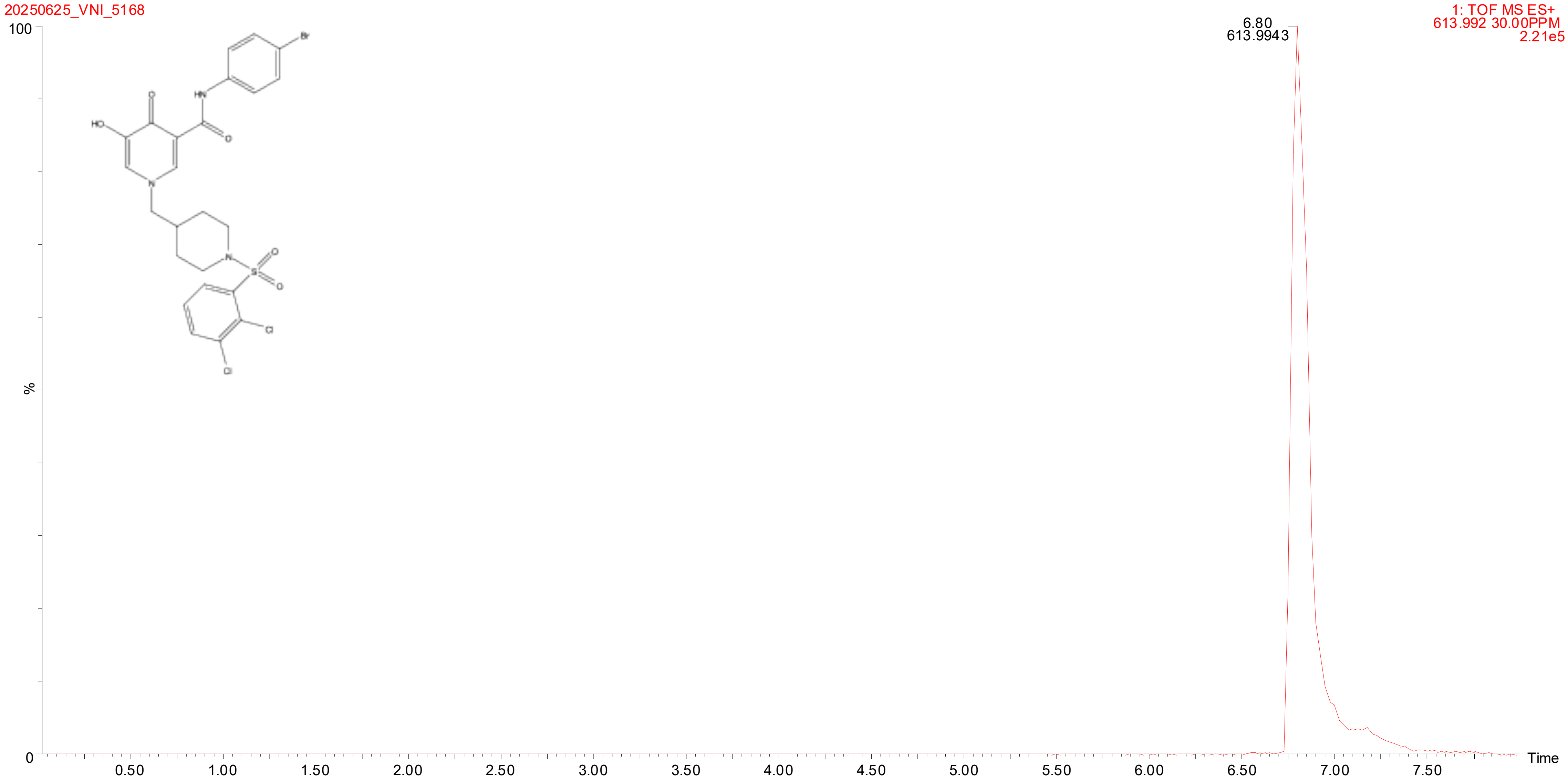
20250625_VNI_5168
1: TOF MS ES+
613.992 30.00PPM
2.21e5
6.80
613.9943
100
%
0
Time
0.50
1.00
1.50
2.00
2.50
3.00
3.50
4.00
4.50
5.00
5.50
6.00
6.50
7.00
7.50

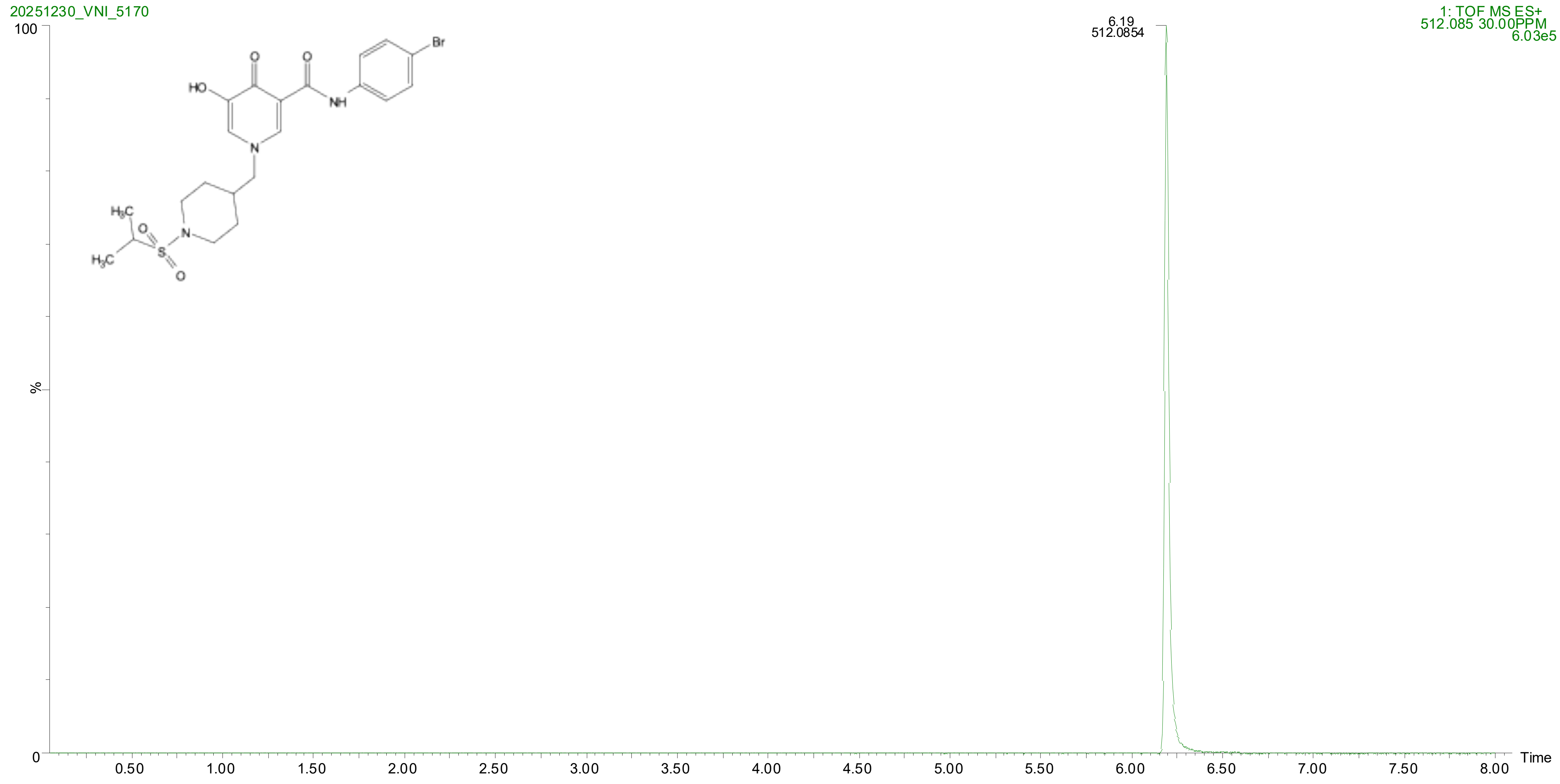

20251230_VNI_5170
1: TOF MS ES+
512.085 30.00PPM
6.03e5
6.19
512.0854
100
%
0
0.50
1.00
1.50
2.00
2.50
3.00
3.50
4.00
4.50
5.00
5.50
6.00
6.50
7.00
7.50
8.00
Time

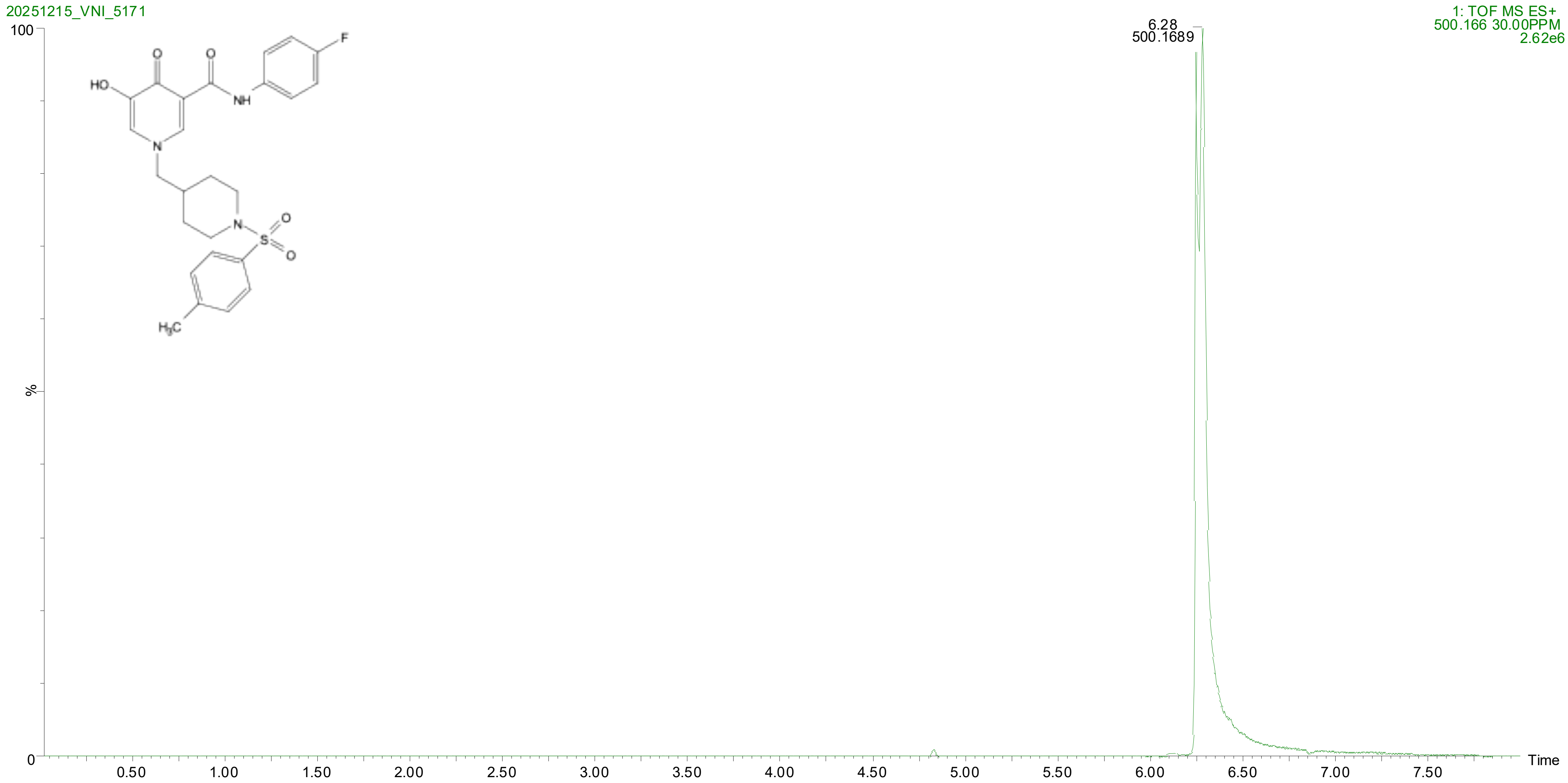

20251215_VNI_5171
1: TOF MS ES+
500.166 30.00PPM
2.62e6
6.28
500.1689
100
%
0
Time
0.50
1.00
1.50
2.00
2.50
3.00
3.50
4.00
4.50
5.00
5.50
6.00
6.50
7.00
7.50
HO
O
O
NH
F
N
N
S
O
O
H3C

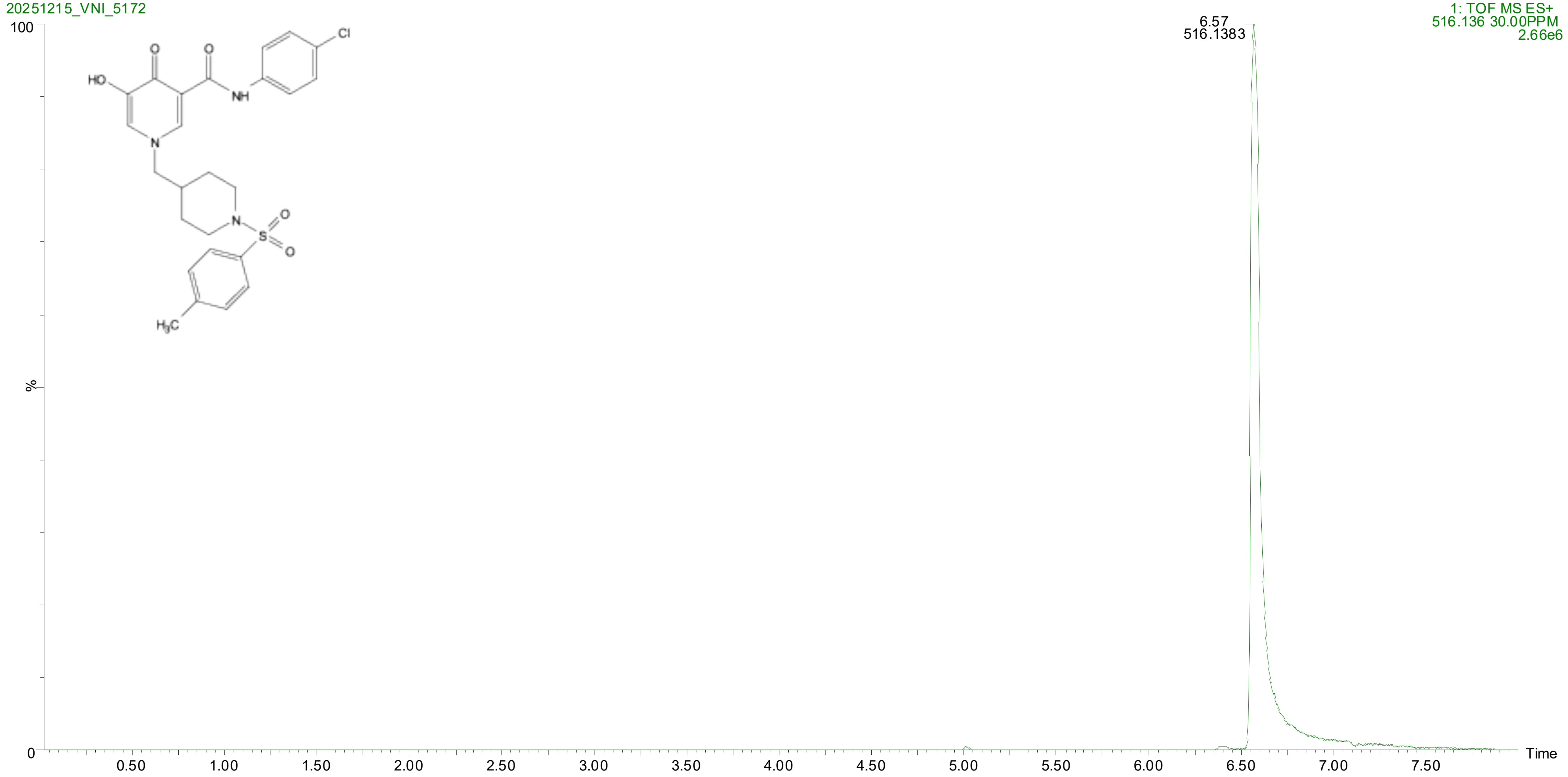
20251215_VNI_5172
1: TOF MS ES+
516.136 30.00PPM
2.66e6
6.57
516.1383
100
%
0
Time
0.50
1.00
1.50
2.00
2.50
3.00
3.50
4.00
4.50
5.00
5.50
6.00
6.50
7.00
7.50
HO
O
O
NH
Cl
N
N
S
O
O
H3C

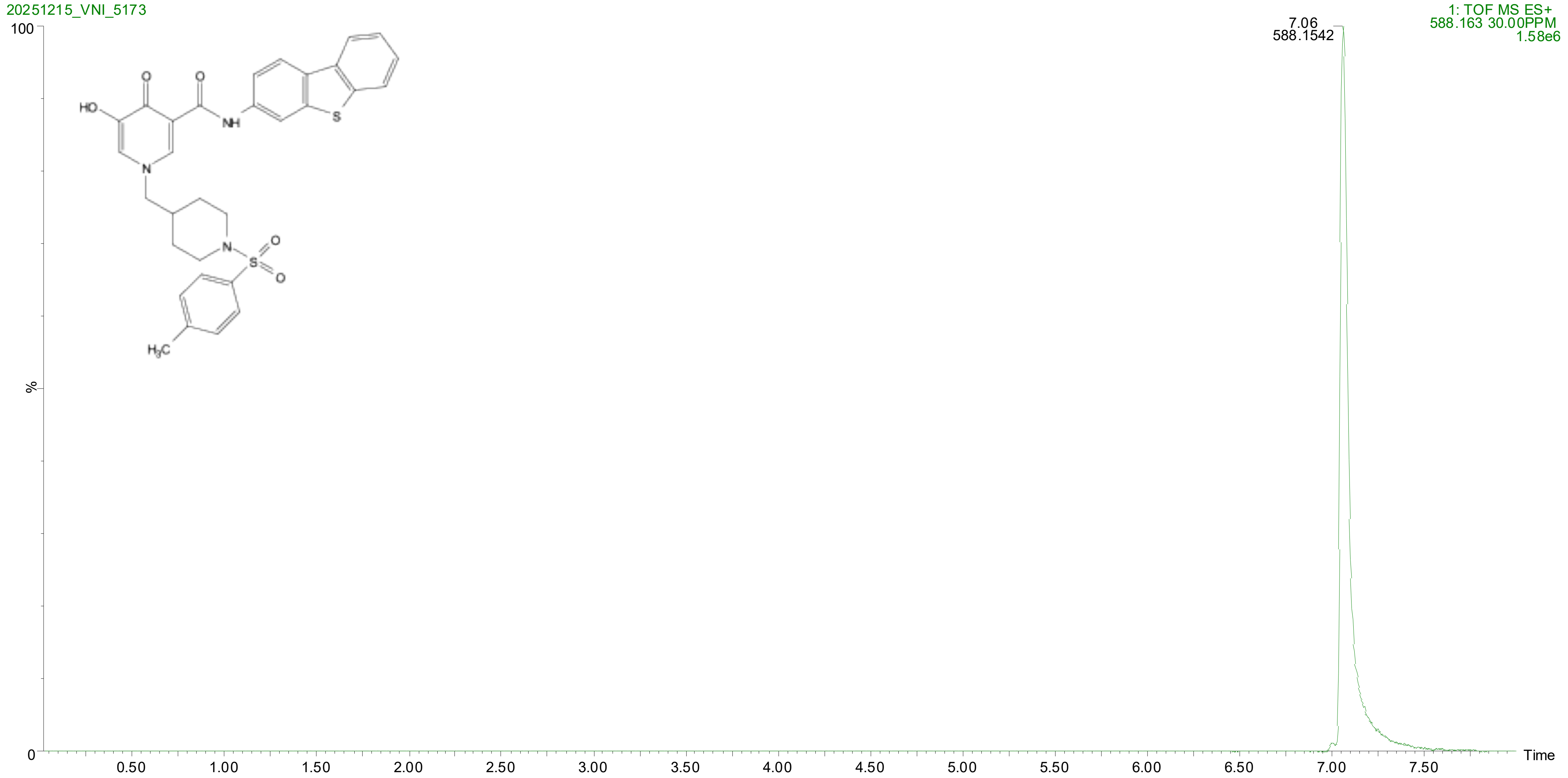

20251215_VNI_5173
1: TOF MS ES+
588.163 30.00PPM
1.58e6
7.06
588.1542
100
%
0
Time
0.50
1.00
1.50
2.00
2.50
3.00
3.50
4.00
4.50
5.00
5.50
6.00
6.50
7.00
7.50
HO
O
O
NH
S
N
N
S
O
O
H3C

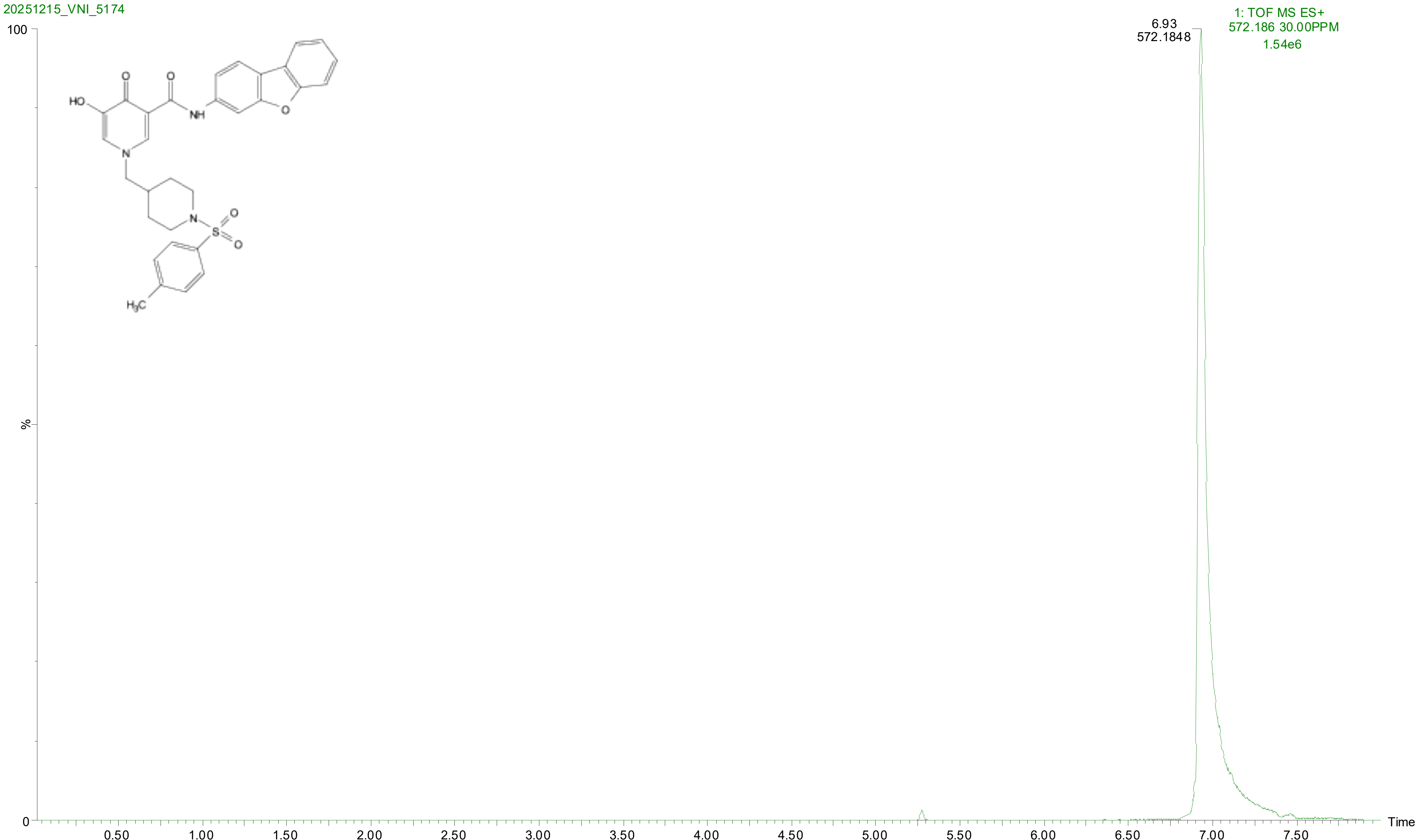
20251215_VNI_5174
1: TOF MS ES+
572.186 30.00PPM
1.54e6
6.93
572.1848
100
%
0
HO
O
O
NH
O
N
N
S
O
O
H3C
0.50
1.00
1.50
2.00
2.50
3.00
3.50
4.00
4.50
5.00
5.50
6.00
6.50
7.00
7.50
Time

HPLC data for compounds

**Default file**

VNI-5162_10 mM Sm (Mn, 2x3)

Diode Array
Range: 5.667e+1
Area

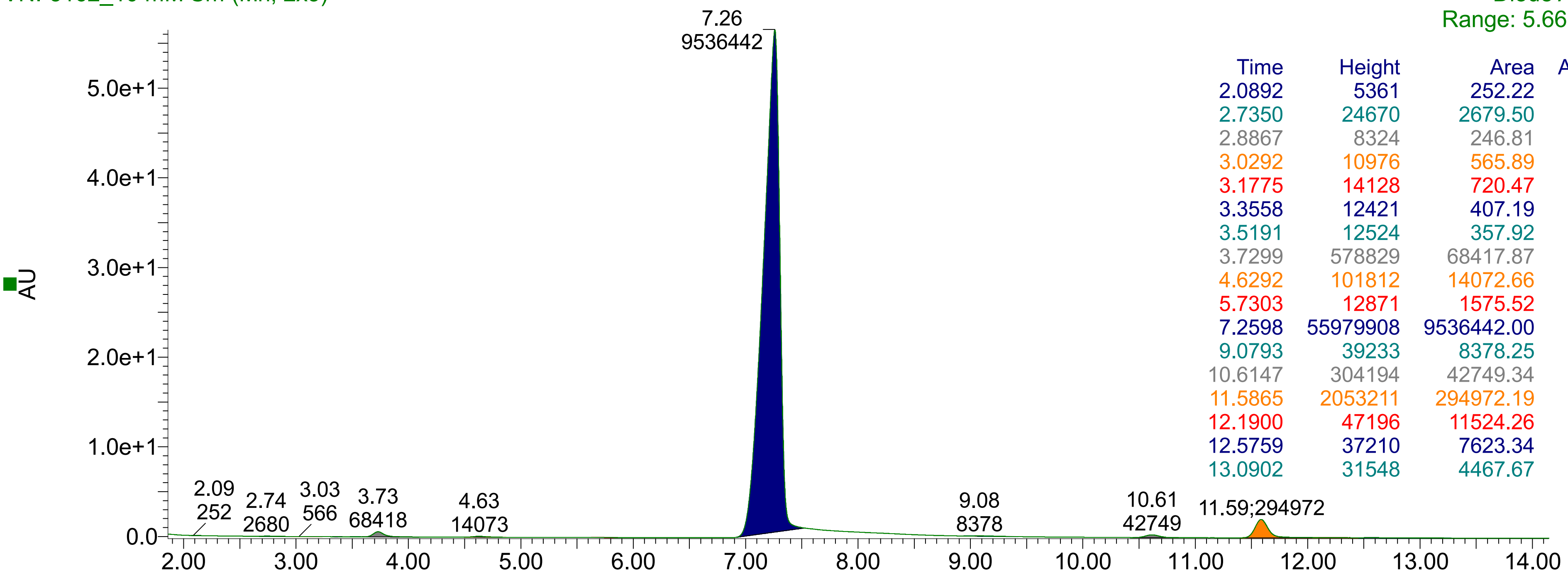


| Time | Height | Area | Area% |
|---|---|---|---|
| 2.0892 | 5361 | 252.22 | 0.00 |
| 2.7350 | 24670 | 2679.50 | 0.03 |
| 2.8867 | 8324 | 246.81 | 0.00 |
| 3.0292 | 10976 | 565.89 | 0.01 |
| 3.1775 | 14128 | 720.47 | 0.01 |
| 3.3558 | 12421 | 407.19 | 0.00 |
| 3.5191 | 12524 | 357.92 | 0.00 |
| 3.7299 | 578829 | 68417.87 | 0.68 |
| 4.6292 | 101812 | 14072.66 | 0.14 |
| 5.7303 | 12871 | 1575.52 | 0.02 |
| 7.2598 | 55979908 | 9536442.00 | 95.41 |
| 9.0793 | 39233 | 8378.25 | 0.08 |
| 10.6147 | 304194 | 42749.34 | 0.43 |
| 11.5865 | 2053211 | 294972.19 | 2.95 |
| 12.1900 | 47196 | 11524.26 | 0.12 |
| 12.5759 | 37210 | 7623.34 | 0.08 |
| 13.0902 | 31548 | 4467.67 | 0.04 |

VNI-5162_10 mM

Diode Array
Range: 5.673e+1

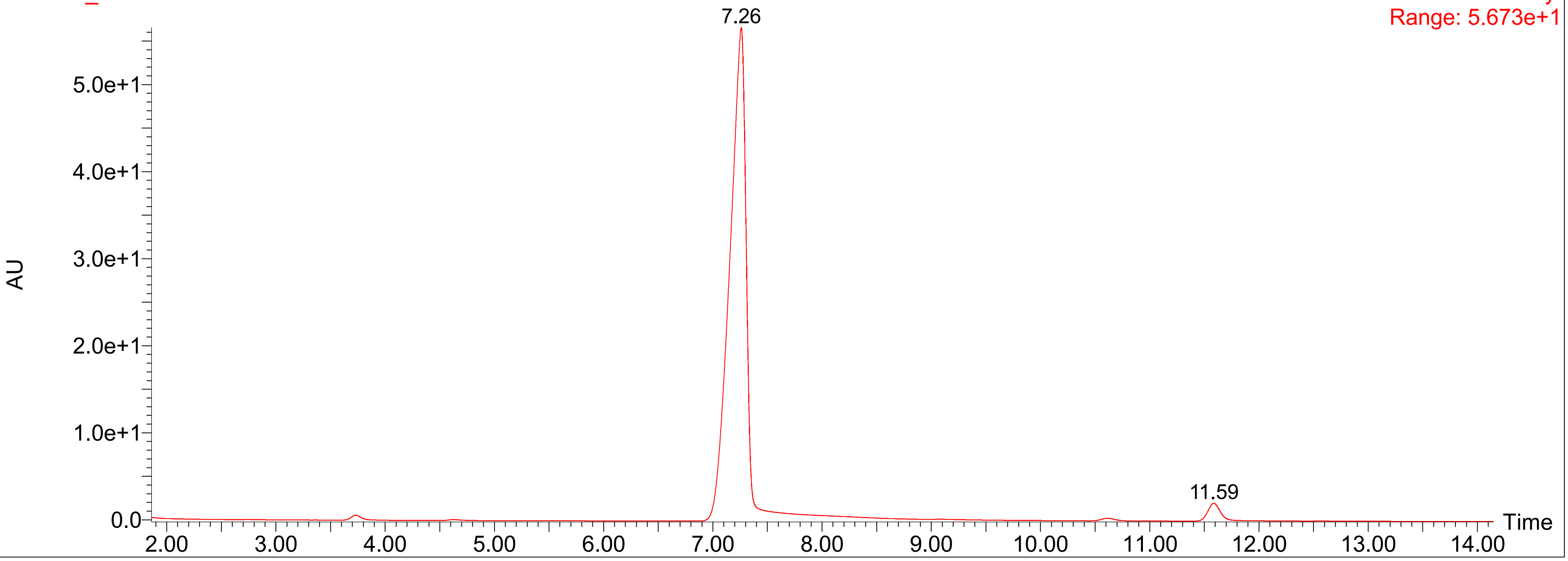

**Default file**

VNI-5163_10 mM Sm (Mn, 2x3)

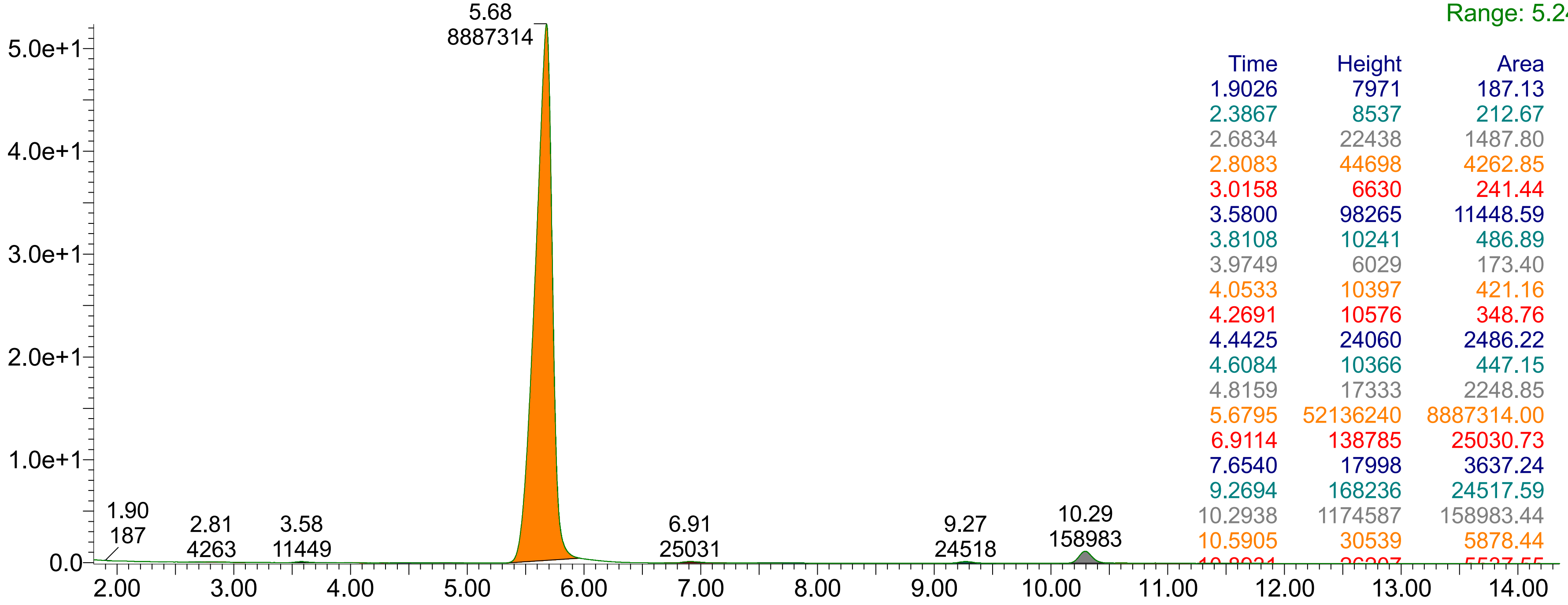


| Time | Height | Area | Area% |
|---|---|---|---|
| 1.9026 | 7971 | 187.13 | 0.00 |
| 2.3867 | 8537 | 212.67 | 0.00 |
| 2.6834 | 22438 | 1487.80 | 0.02 |
| 2.8083 | 44698 | 4262.85 | 0.05 |
| 3.0158 | 6630 | 241.44 | 0.00 |
| 3.5800 | 98265 | 11448.59 | 0.13 |
| 3.8108 | 10241 | 486.89 | 0.01 |
| 3.9749 | 6029 | 173.40 | 0.00 |
| 4.0533 | 10397 | 421.16 | 0.00 |
| 4.2691 | 10576 | 348.76 | 0.00 |
| 4.4425 | 24060 | 2486.22 | 0.03 |
| 4.6084 | 10366 | 447.15 | 0.00 |
| 4.8159 | 17333 | 2248.85 | 0.02 |
| 5.6795 | 52136240 | 8887314.00 | 97.27 |
| 6.9114 | 138785 | 25030.73 | 0.27 |
| 7.6540 | 17998 | 3637.24 | 0.04 |
| 9.2694 | 168236 | 24517.59 | 0.27 |
| 10.2938 | 1174587 | 158983.44 | 1.74 |
| 10.5905 | 30539 | 5878.44 | 0.06 |
| [illegible] | [illegible] | [illegible] | [illegible] |

VNI-5163_10 mM

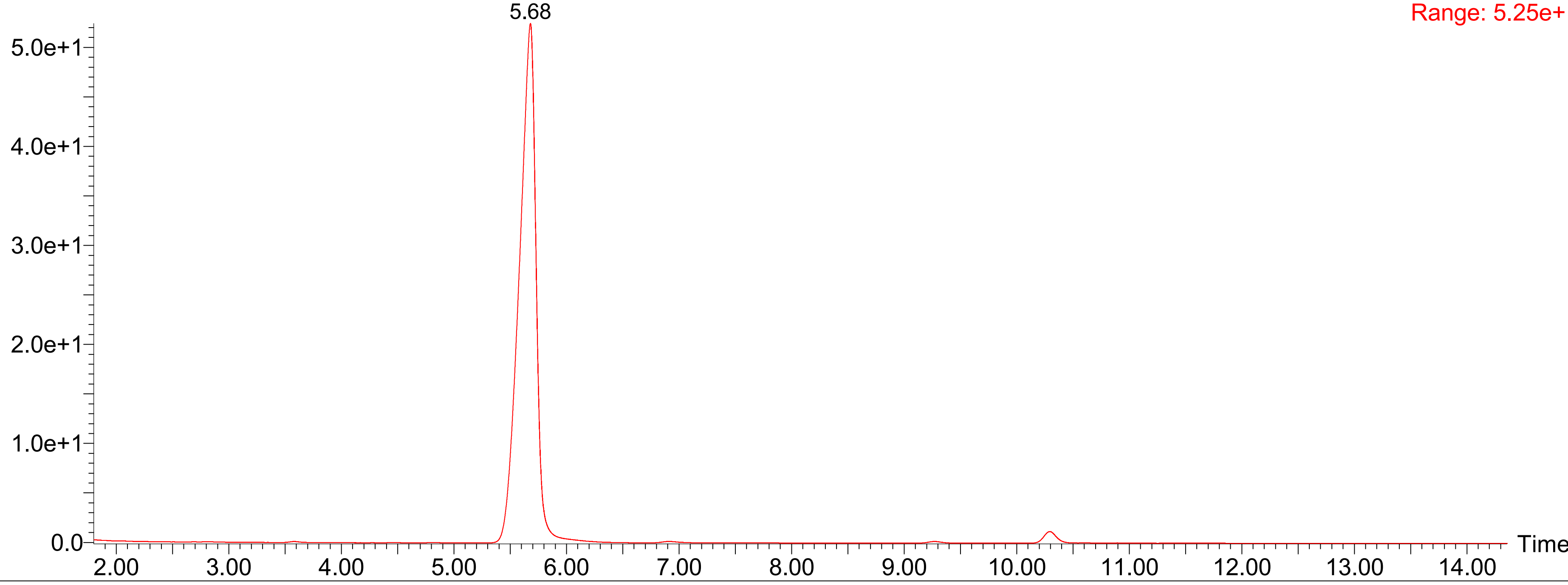

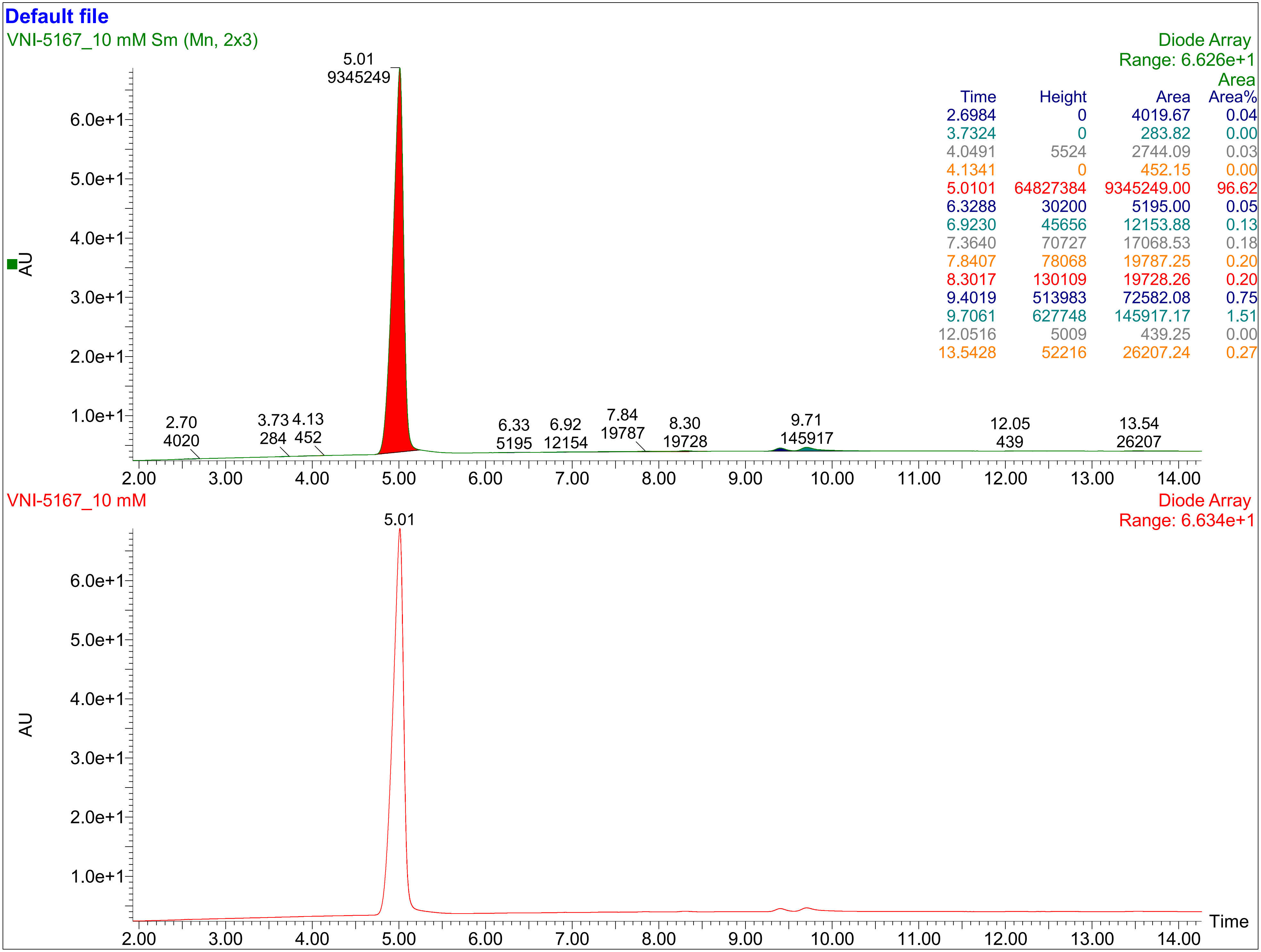


| Time | Height | Area | Area Area% |
|---|---|---|---|
| 2.6984 | 0 | 4019.67 | 0.04 |
| 3.7324 | 0 | 283.82 | 0.00 |
| 4.0491 | 5524 | 2744.09 | 0.03 |
| 4.1341 | 0 | 452.15 | 0.00 |
| 5.0101 | 64827384 | 9345249.00 | 96.62 |
| 6.3288 | 30200 | 5195.00 | 0.05 |
| 6.9230 | 45656 | 12153.88 | 0.13 |
| 7.3640 | 70727 | 17068.53 | 0.18 |
| 7.8407 | 78068 | 19787.25 | 0.20 |
| 8.3017 | 130109 | 19728.26 | 0.20 |
| 9.4019 | 513983 | 72582.08 | 0.75 |
| 9.7061 | 627748 | 145917.17 | 1.51 |
| 12.0516 | 5009 | 439.25 | 0.00 |
| 13.5428 | 52216 | 26207.24 | 0.27 |

**Default file**

VNI-5168_10 mM Sm (Mn, 2x3)

Diode Array
Range: 3.933e+1

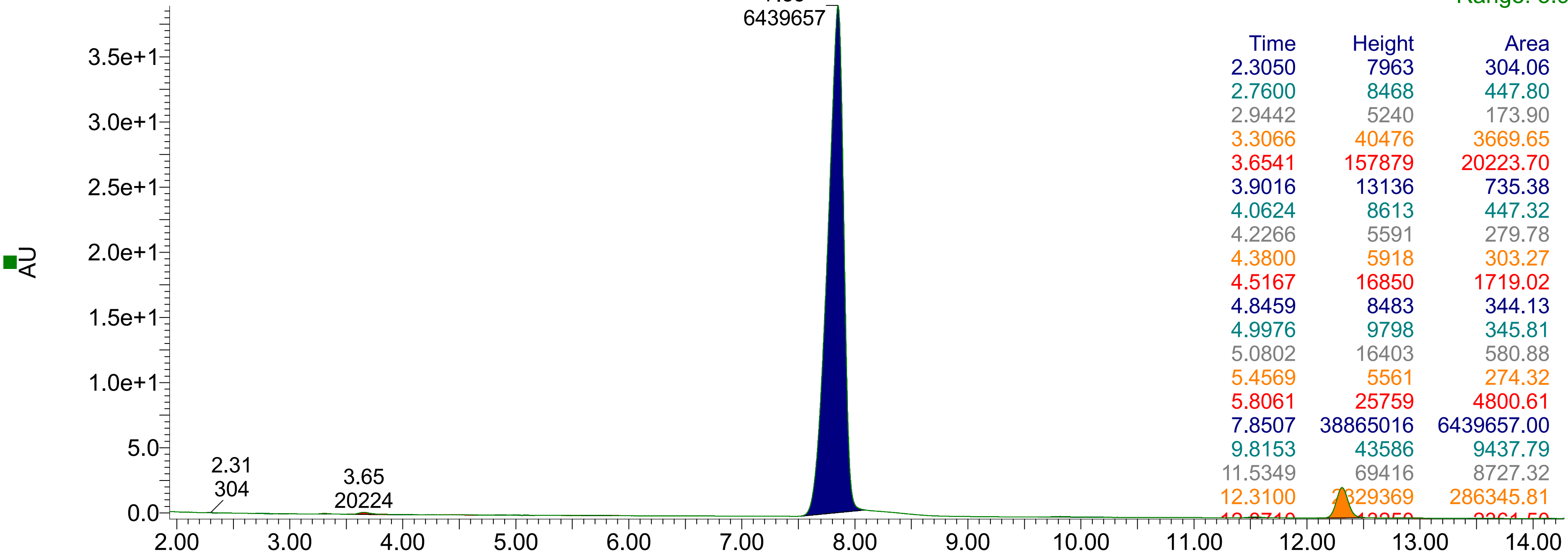


| Time | Height | Area | Area Area% |
|---|---|---|---|
| 2.3050 | 7963 | 304.06 | 0.00 |
| 2.7600 | 8468 | 447.80 | 0.01 |
| 2.9442 | 5240 | 173.90 | 0.00 |
| 3.3066 | 40476 | 3669.65 | 0.05 |
| 3.6541 | 157879 | 20223.70 | 0.30 |
| 3.9016 | 13136 | 735.38 | 0.01 |
| 4.0624 | 8613 | 447.32 | 0.01 |
| 4.2266 | 5591 | 279.78 | 0.00 |
| 4.3800 | 5918 | 303.27 | 0.00 |
| 4.5167 | 16850 | 1719.02 | 0.03 |
| 4.8459 | 8483 | 344.13 | 0.01 |
| 4.9976 | 9798 | 345.81 | 0.01 |
| 5.0802 | 16403 | 580.88 | 0.01 |
| 5.4569 | 5561 | 274.32 | 0.00 |
| 5.8061 | 25759 | 4800.61 | 0.07 |
| 7.8507 | 38865016 | 6439657.00 | 94.91 |
| 9.8153 | 43586 | 9437.79 | 0.14 |
| 11.5349 | 69416 | 8727.32 | 0.13 |
| 12.3100 | 2329369 | 286345.81 | 4.22 |
| [illegible] | [illegible] | [illegible] | [illegible] |

VNI-5168_10 mM

Diode Array
Range: 3.937e+1

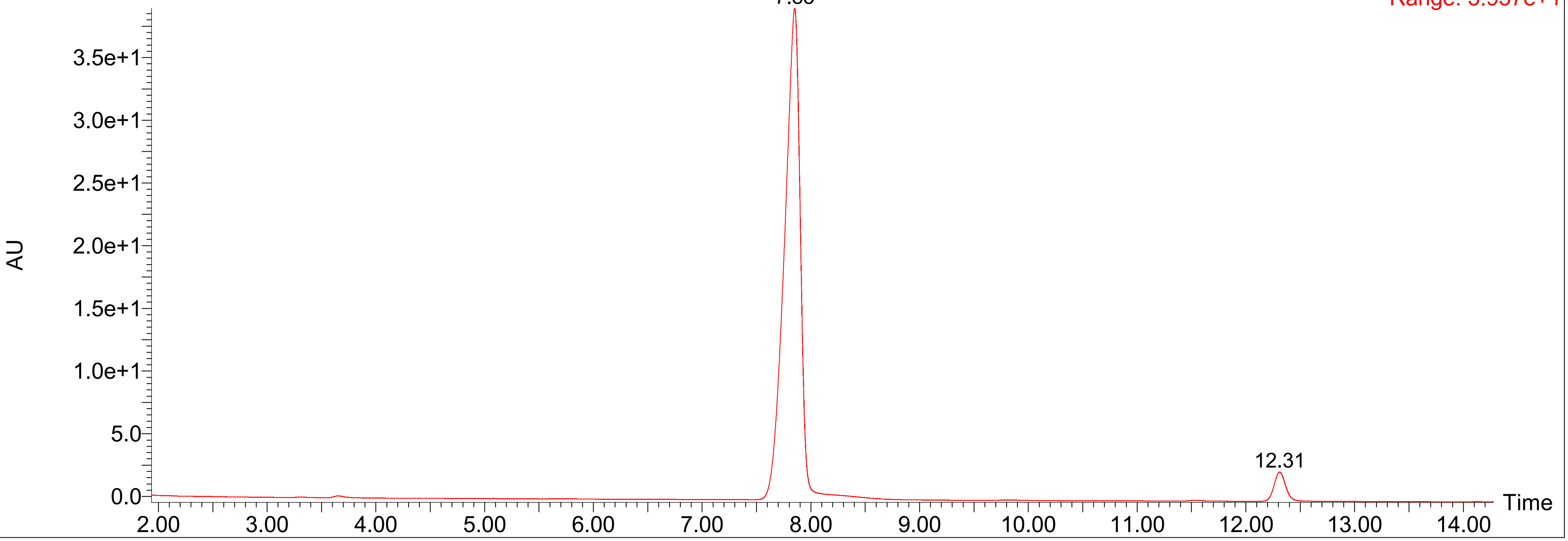

### $^{1}H$ NMR and $^{13}C$ NMR spectra

2

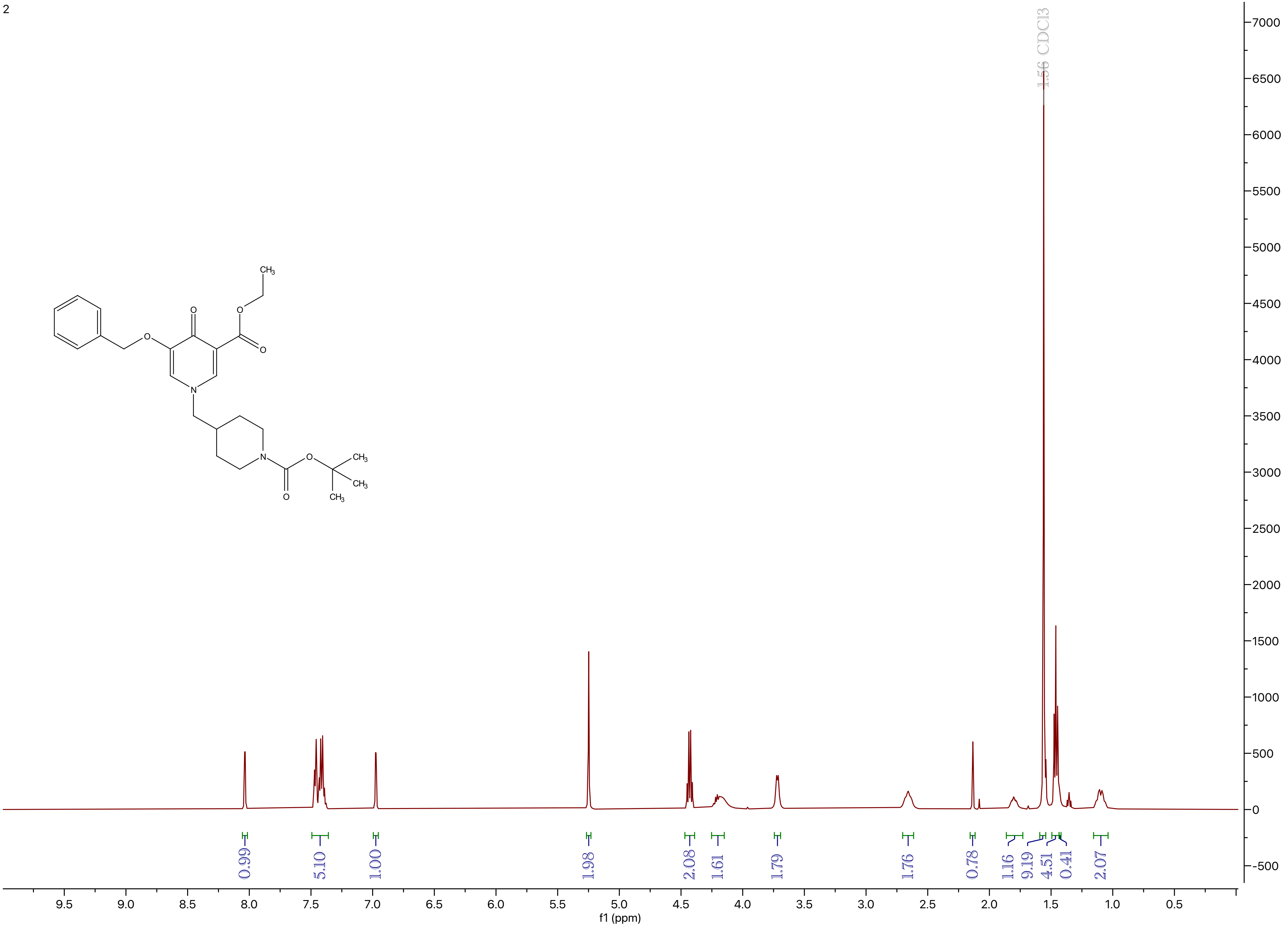

CH3
O
O
O
O
N
N
O
O
CH3
CH3
CH3
1.56 CDCl3
0.99
5.10
1.00
1.98
2.08
1.61
1.79
1.76
0.78
1.16
9.19
4.51
0.41
2.07
9.5
9.0
8.5
8.0
7.5
7.0
6.5
6.0
5.5
5.0
4.5
4.0
3.5
3.0
2.5
2.0
1.5
1.0
0.5
f1 (ppm)
7000
6500
6000
5500
5000
4500
4000
3500
3000
2500
2000
1500
1000
500
0
-500

3

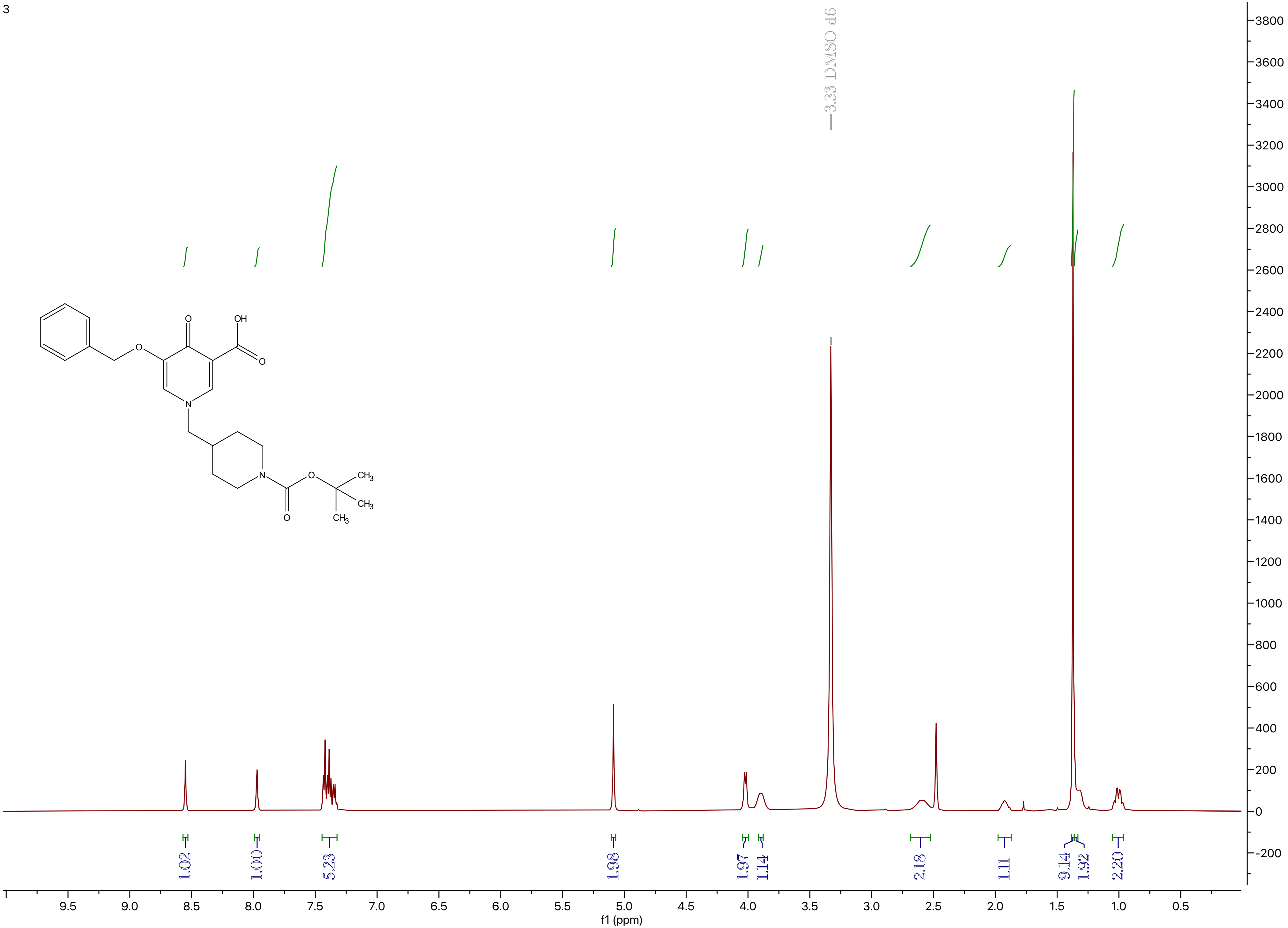
OH
O
O
O
N
N
O
O
CH3
CH3
CH3
—3.33 DMSO-d6
1.02
1.00
5.23
1.98
1.97
1.14
2.18
1.11
9.14
1.92
2.20
9.5
9.0
8.5
8.0
7.5
7.0
6.5
6.0
5.5
5.0
4.5
4.0
3.5
3.0
2.5
2.0
1.5
1.0
0.5
f1 (ppm)
3800
3600
3400
3200
3000
2800
2600
2400
2200
2000
1800
1600
1400
1200
1000
800
600
400
200
0
-200

4

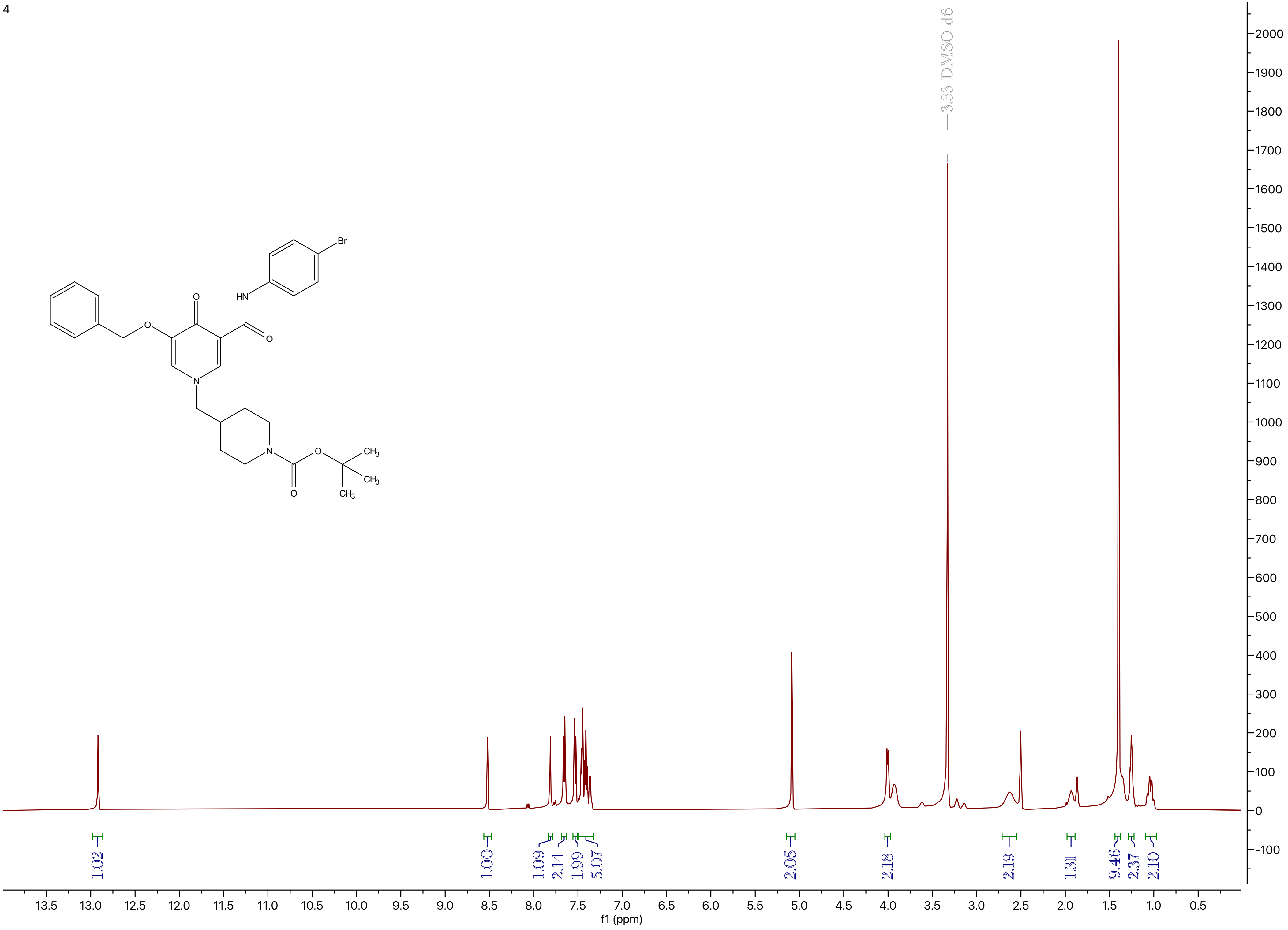

Br
HN
O
O
N
O
N
O
O
CH3
CH3
CH3
3.33 DMSO-d6
1.02
1.00
1.09
2.14
1.99
5.07
2.05
2.18
2.19
1.31
9.46
2.37
2.10
13.5 13.0 12.5 12.0 11.5 11.0 10.5 10.0 9.5 9.0 8.5 8.0 7.5 7.0 6.5 6.0 5.5 5.0 4.5 4.0 3.5 3.0 2.5 2.0 1.5 1.0 0.5
f1 (ppm)
2000 1900 1800 1700 1600 1500 1400 1300 1200 1100 1000 900 800 700 600 500 400 300 200 100 0 -100

5

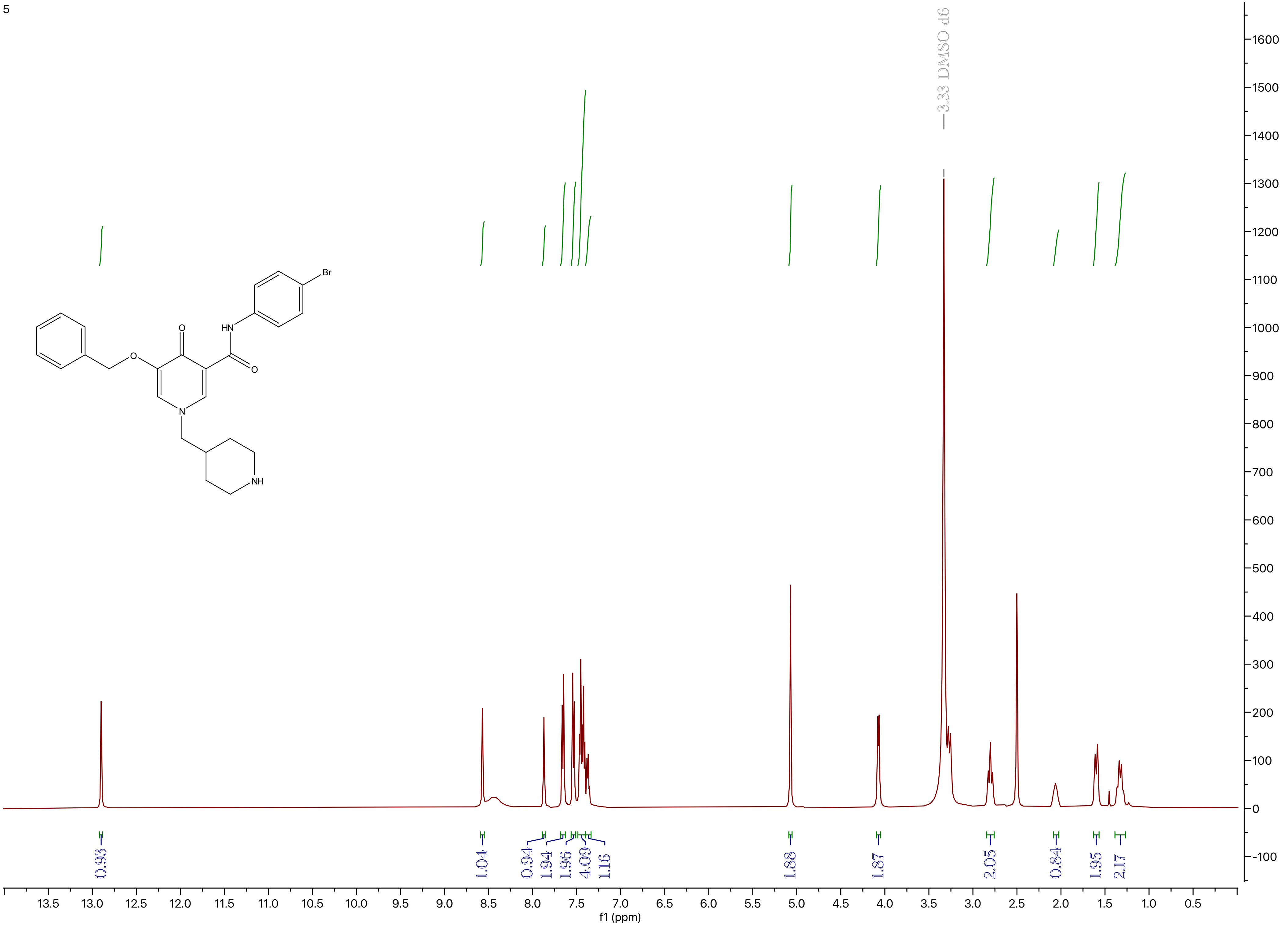

3.33 DMSO-d6
Br
HN
O
O
O
N
NH
0.93
1.04
0.94
1.94
1.96
4.09
1.16
1.88
1.87
2.05
0.84
1.95
2.17
1600
1500
1400
1300
1200
1100
1000
900
800
700
600
500
400
300
200
100
0
-100
13.5
13.0
12.5
12.0
11.5
11.0
10.5
10.0
9.5
9.0
8.5
8.0
7.5
7.0
6.5
6.0
5.5
5.0
4.5
4.0
3.5
3.0
2.5
2.0
1.5
1.0
0.5
f1 (ppm)

VNI-5158

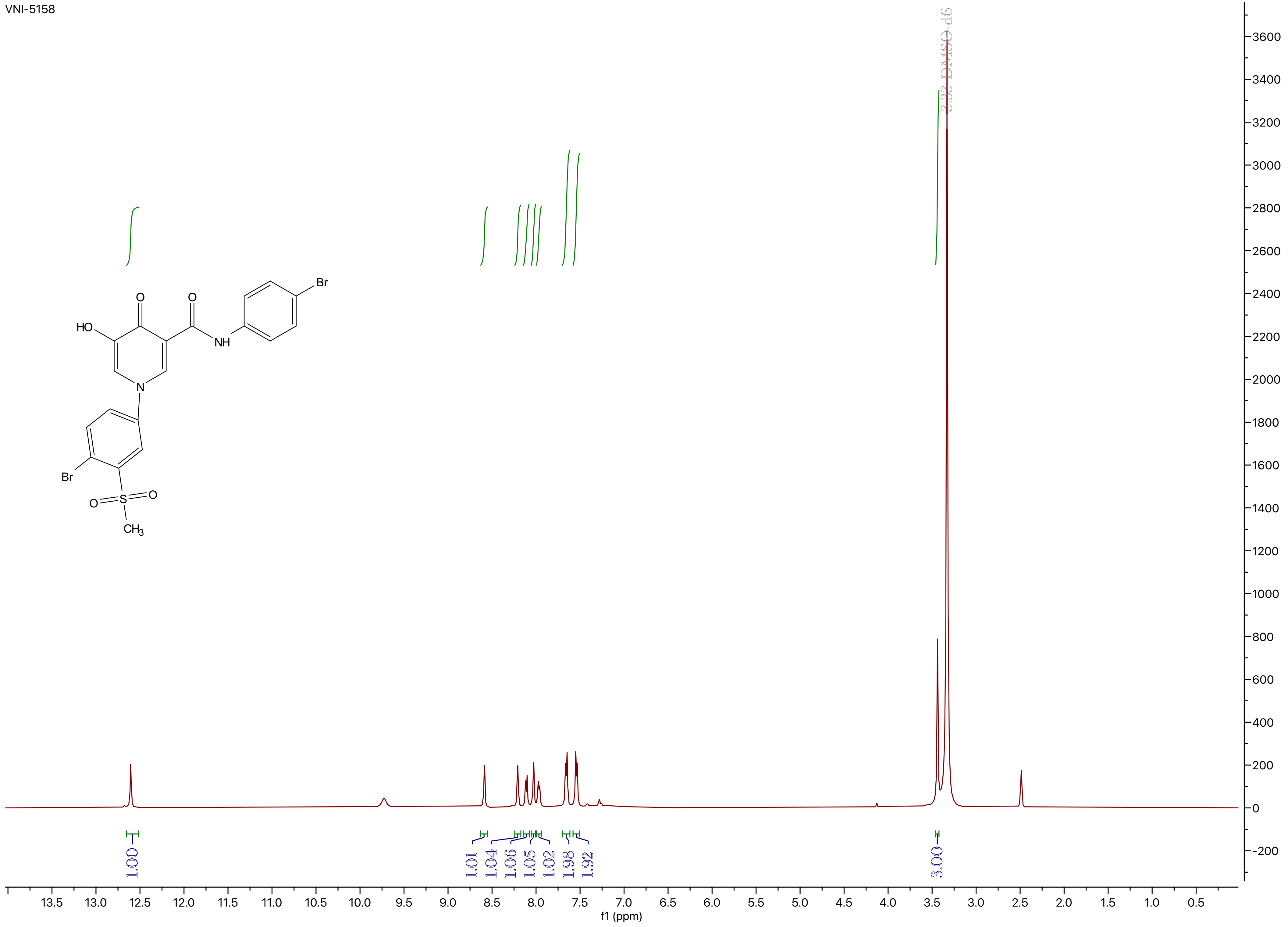

3.33 DMSO-d6
1.00
1.01
1.04
1.06
1.05
1.02
1.98
1.92
3.00
f1 (ppm)
HO
O
O
NH
Br
N
Br
O
S
O
CH3

VNI-5158

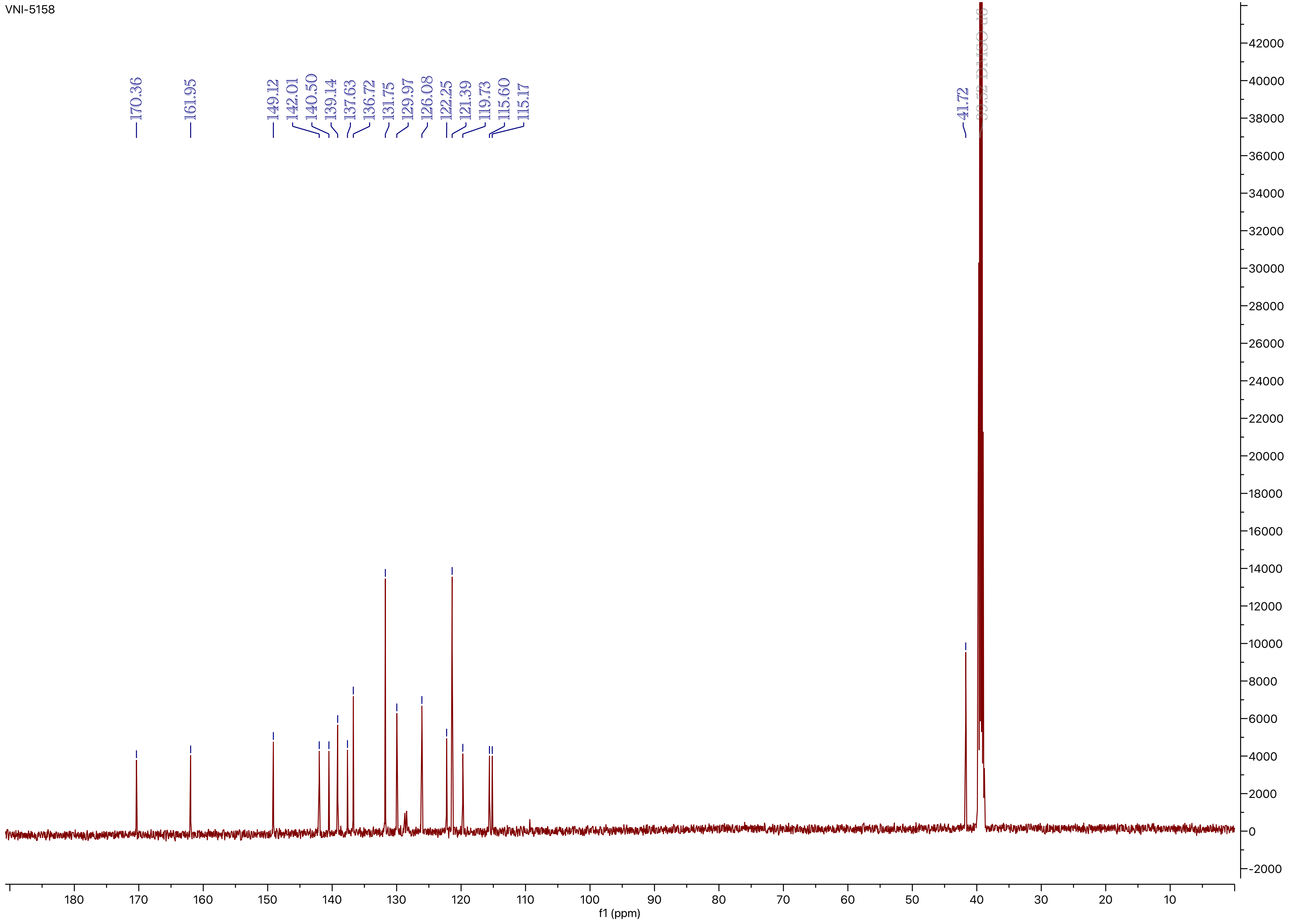
170.36
161.95
149.12
142.01
140.50
139.14
137.63
136.72
131.75
129.97
126.08
122.25
121.39
119.73
115.60
115.17
41.72
39.52 DMSO-d6
f1 (ppm)

VNI-5159

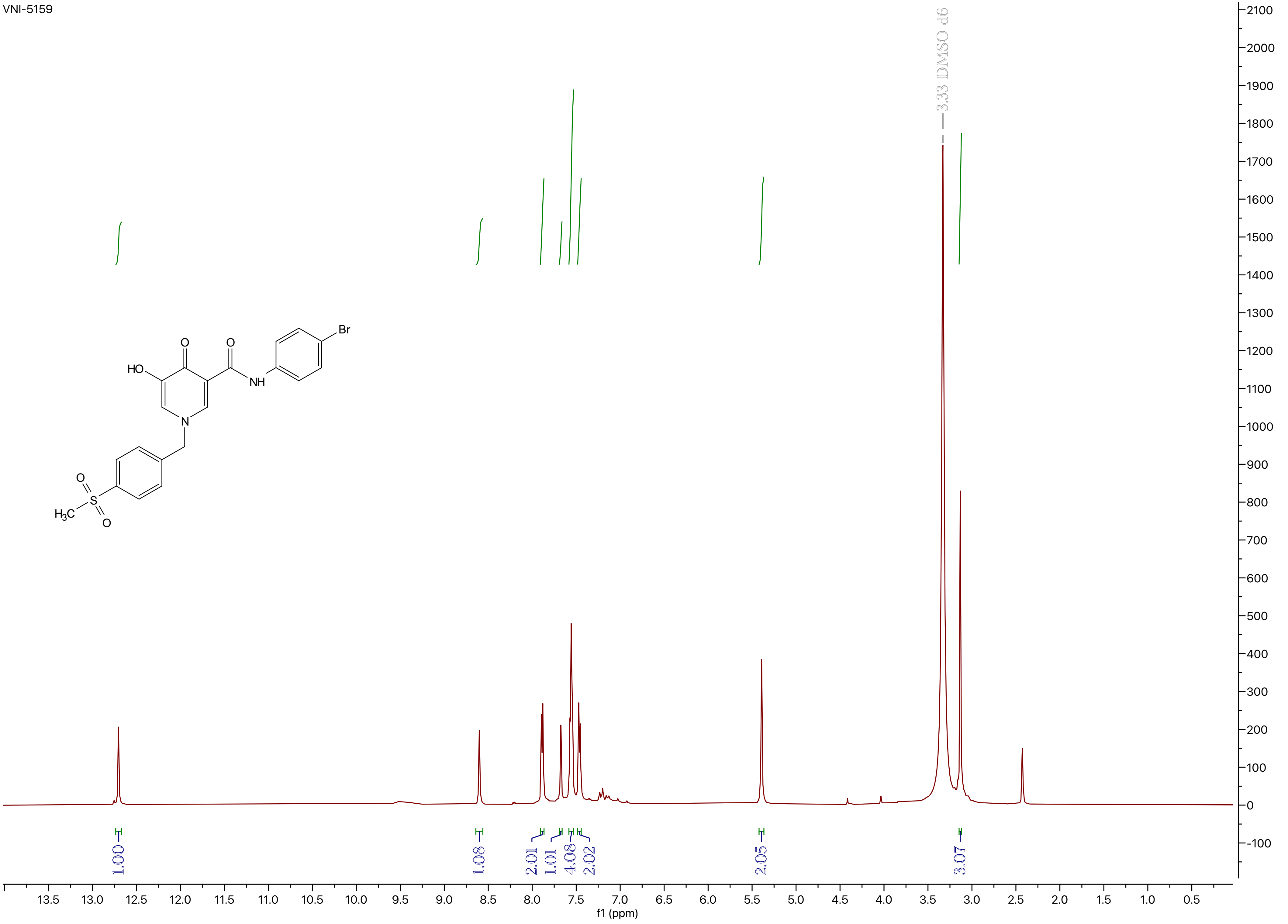

3.33 DMSO-d6
HO
O
O
NH
Br
N
O
S
O
H3C
1.00
1.08
2.01
1.01
4.08
2.02
2.05
3.07
f1 (ppm)

VNI-5159

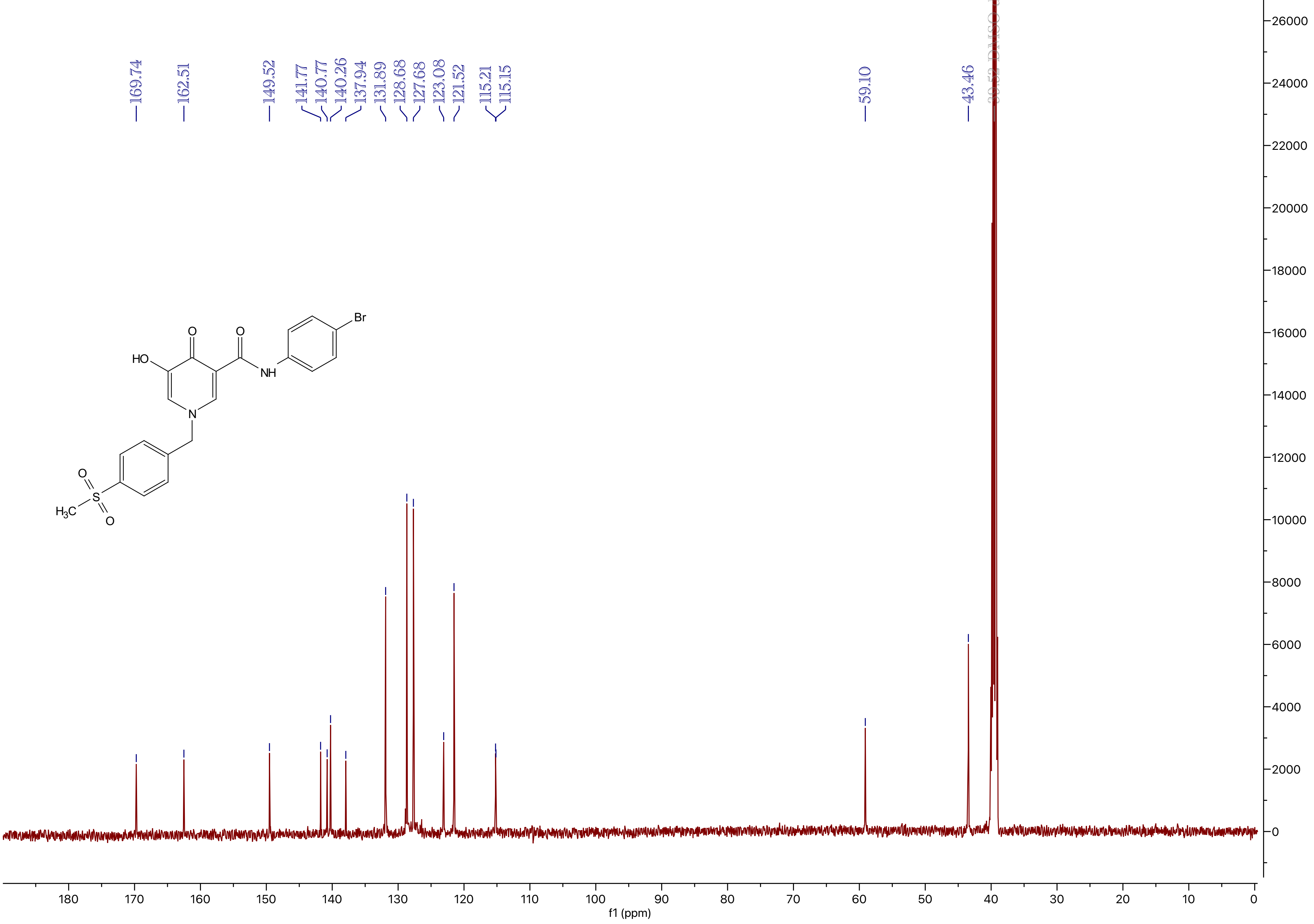
169.74
162.51
149.52
141.77
140.77
140.26
137.94
131.89
128.68
127.68
123.08
121.52
115.21
115.15
59.10
43.46
HO
O
O
NH
Br
N
O
S
O
H3C
f1 (ppm)

VNI-5160

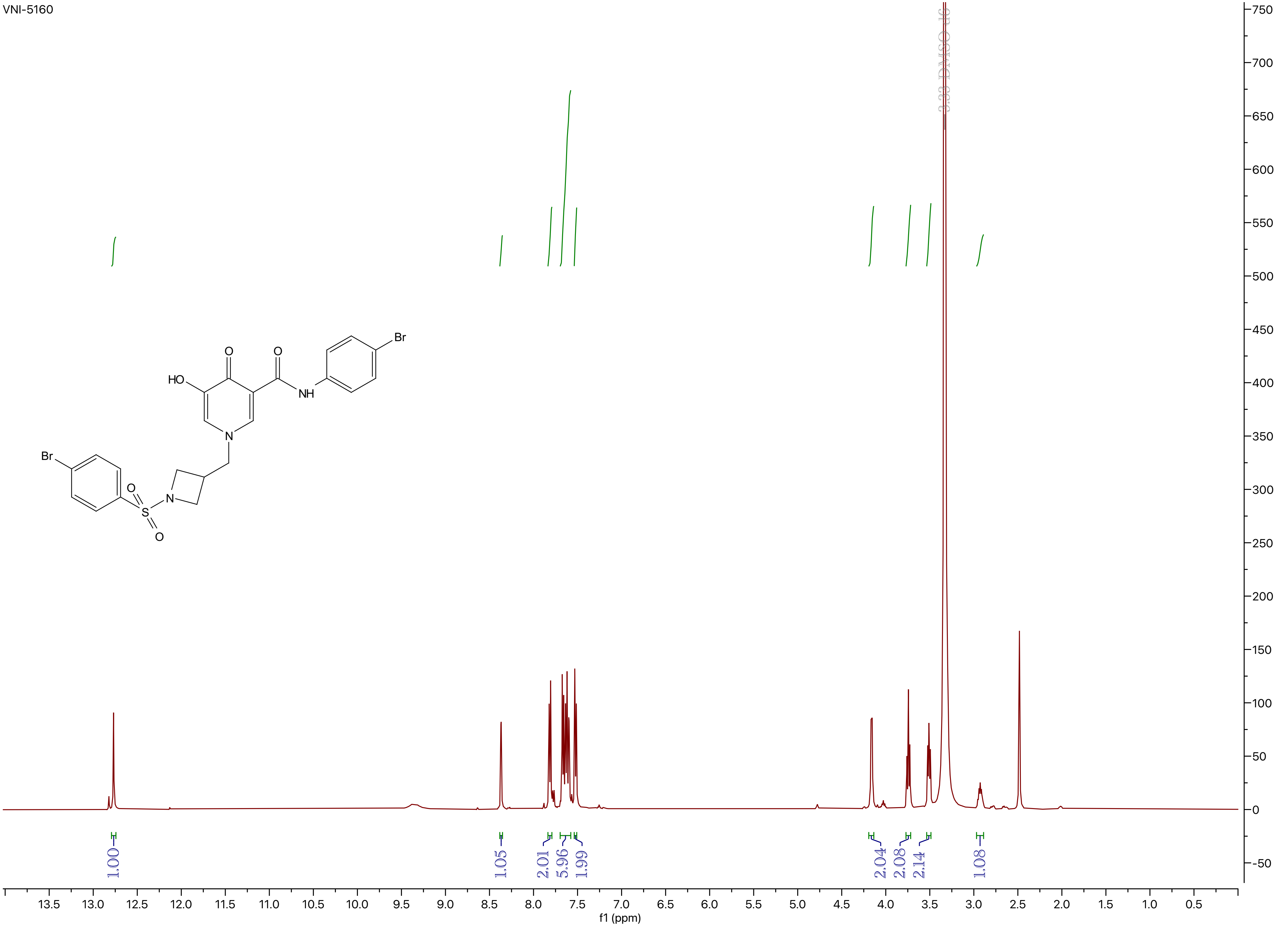
3.33 DMSO-d6
1.00
1.05
2.01
5.96
1.99
2.04
2.08
2.14
1.08
f1 (ppm)
HO
O
O
NH
Br
N
Br
O
S
O
N

VNI-5160

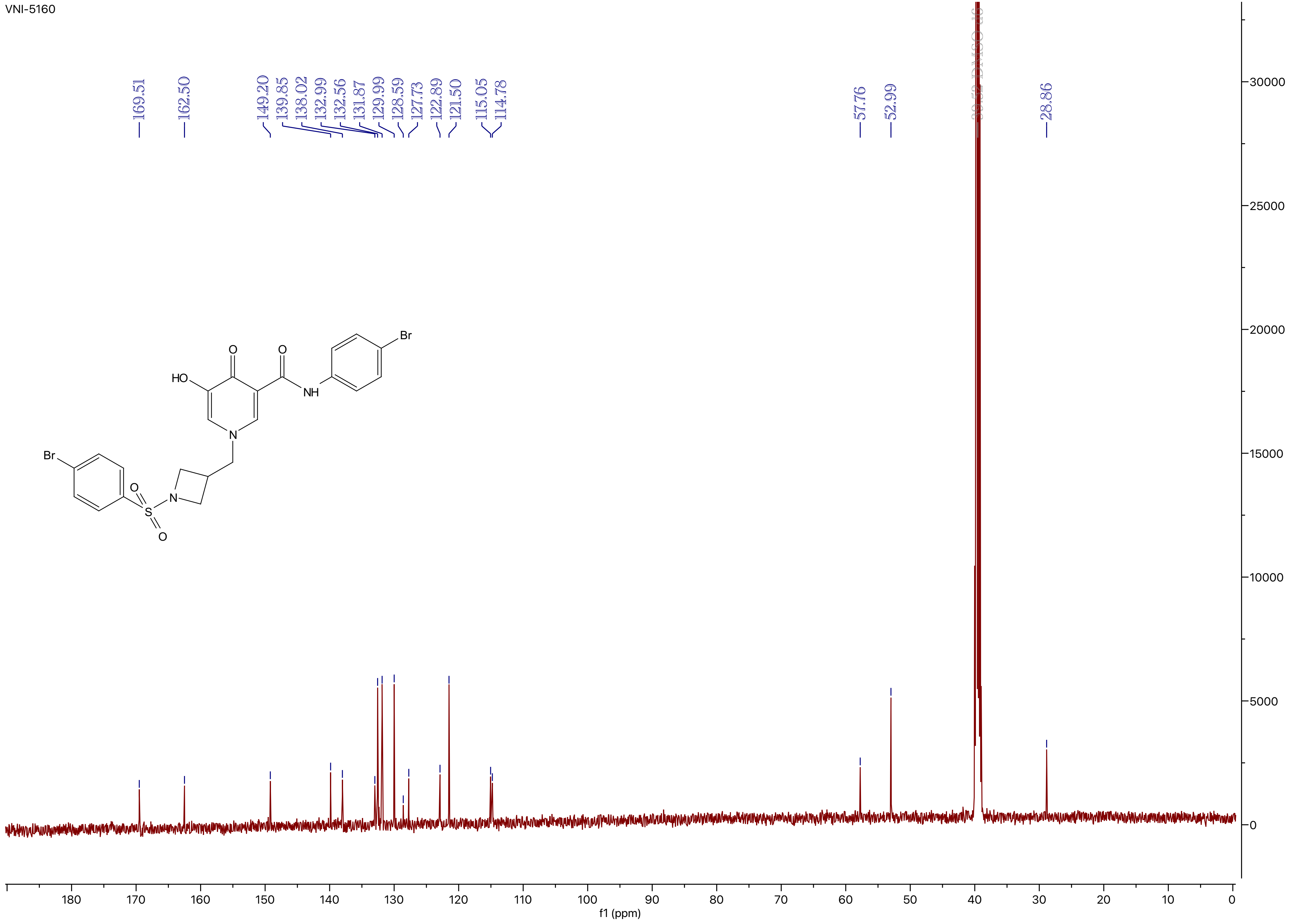
169.51
162.50
149.20
139.85
138.02
132.99
132.56
131.87
129.99
128.59
127.73
122.89
121.50
115.05
114.78
57.76
52.99
39.52 DMSO-d6
28.86
30000
25000
20000
15000
10000
5000
0
180
170
160
150
140
130
120
110
100
90
80
70
60
50
40
30
20
10
0
f1 (ppm)
HO
O
O
NH
Br
N
Br
O
S
O
N

VNI-5161

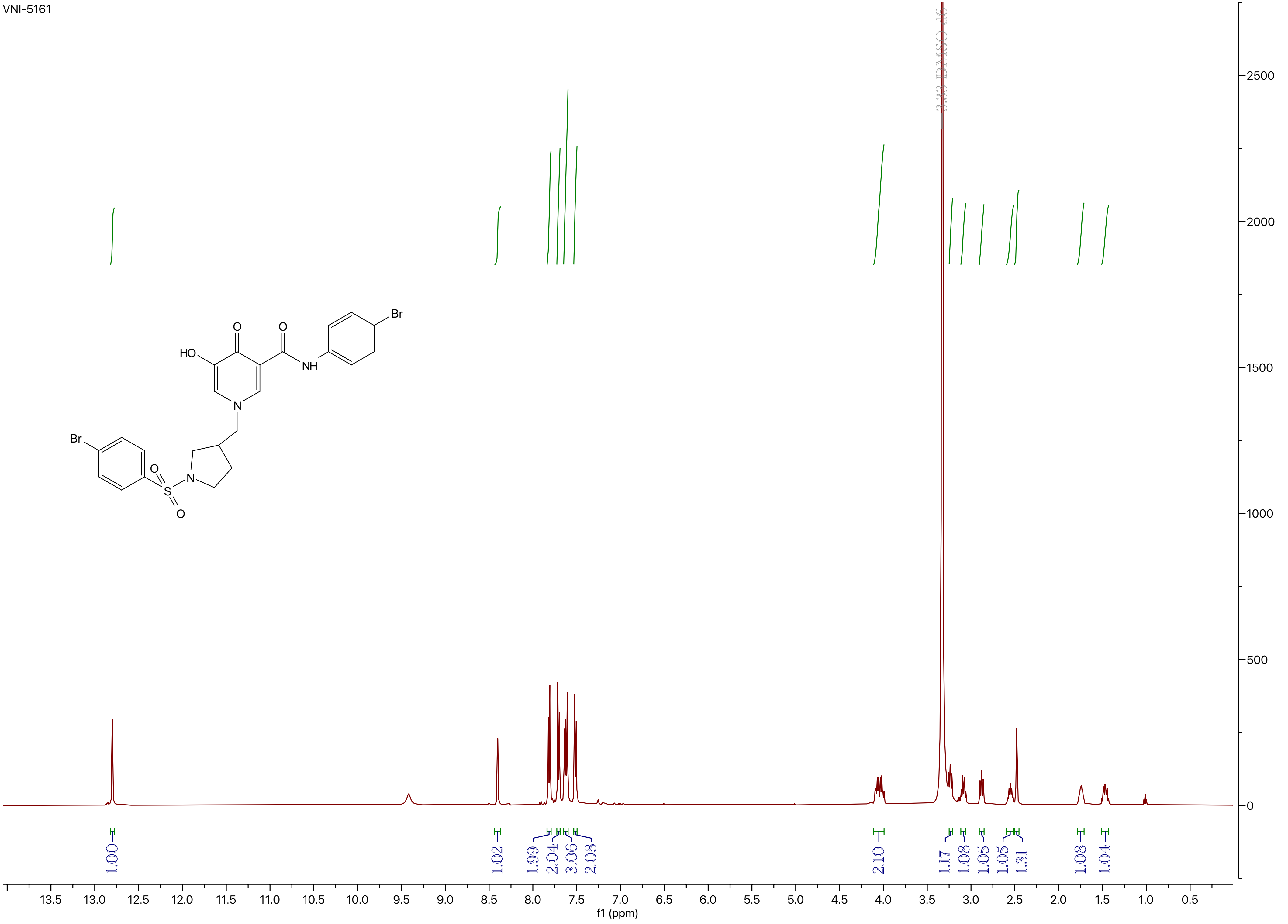
3.33 DMSO-d6
Br
O
O
HO
NH
N
Br
O
S
N
O
1.00
1.02
1.99
2.04
3.06
2.08
2.10
1.17
1.08
1.05
1.05
1.31
1.08
1.04
13.5 13.0 12.5 12.0 11.5 11.0 10.5 10.0 9.5 9.0 8.5 8.0 7.5 7.0 6.5 6.0 5.5 5.0 4.5 4.0 3.5 3.0 2.5 2.0 1.5 1.0 0.5
f1 (ppm)
2500
2000
1500
1000
500
0

VNI-5161

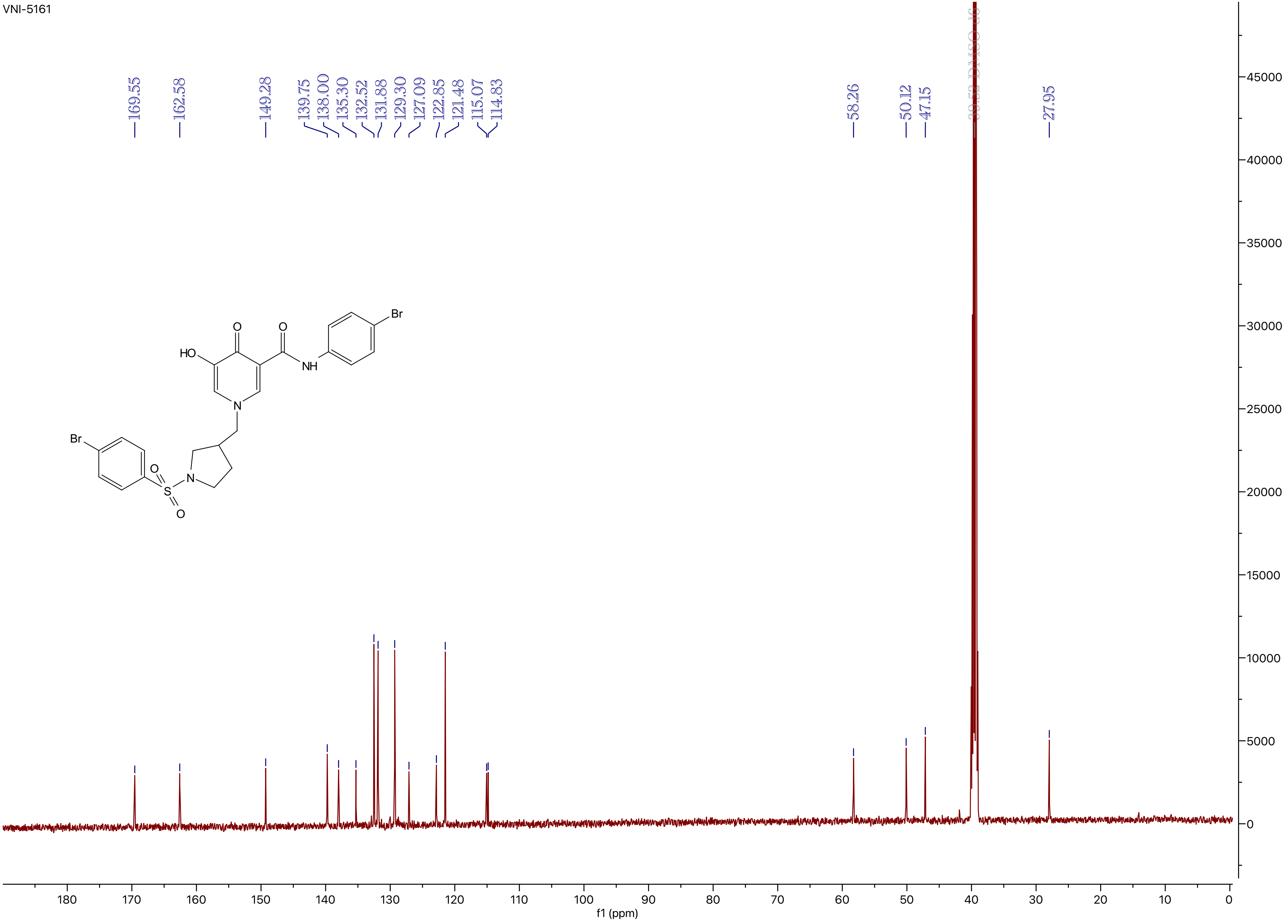

169.55
162.58
149.28
139.75
138.00
135.30
132.52
131.88
129.30
127.09
122.85
121.48
115.07
114.83
58.26
50.12
47.15
39.52 DMSO-d6
27.95
HO
O
O
NH
Br
N
Br
O
S
O
N
45000
40000
35000
30000
25000
20000
15000
10000
5000
0
180
170
160
150
140
130
120
110
100
90
80
70
60
50
40
30
20
10
0
f1 (ppm)

VNI-5162

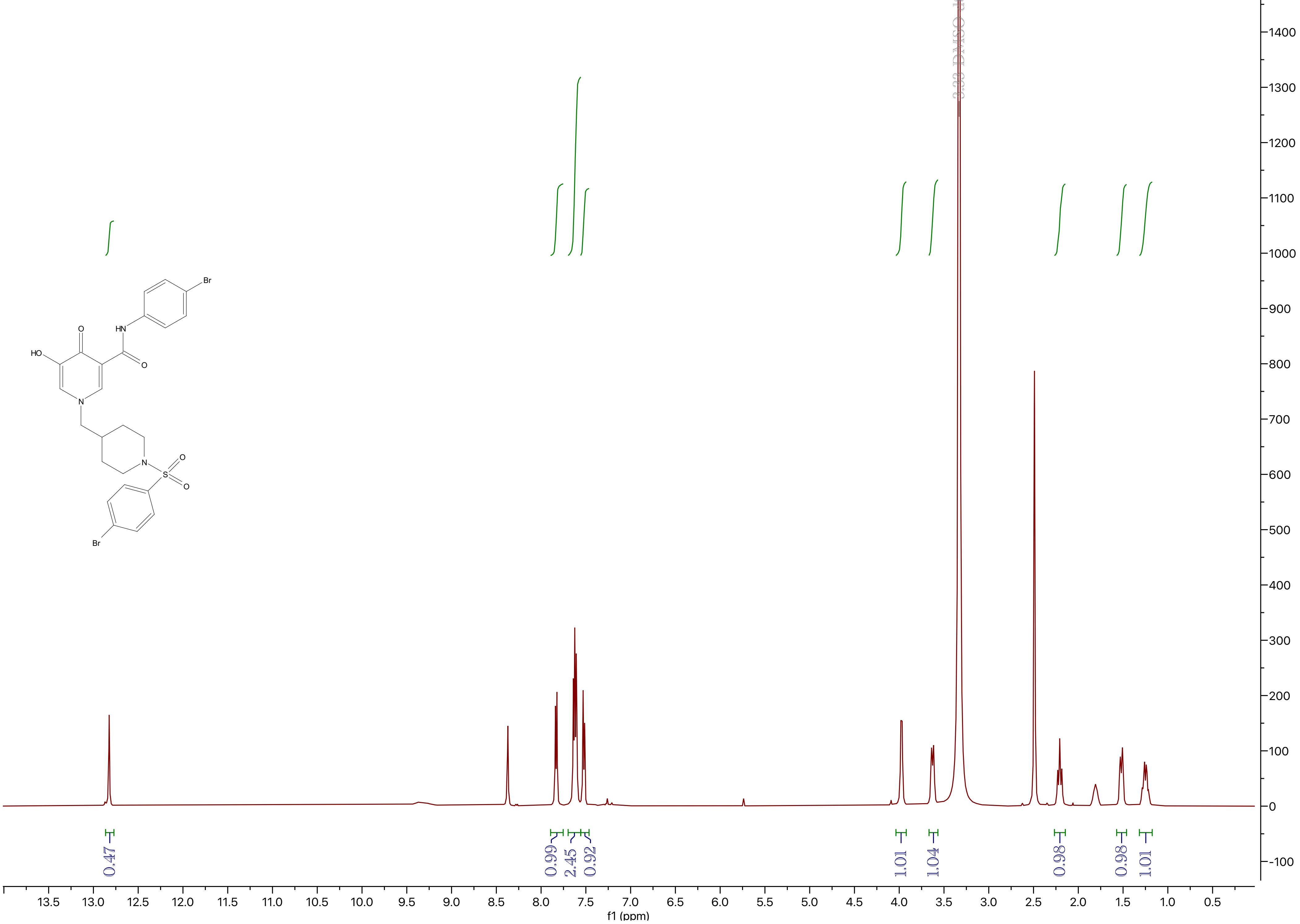
3.33 DMSO-d6
Br
HN
O
HO
N
S
Br
0.47
0.99
2.45
0.92
1.01
1.04
0.98
0.98
1.01
13.5
13.0
12.5
12.0
11.5
11.0
10.5
10.0
9.5
9.0
8.5
8.0
7.5
7.0
6.5
6.0
5.5
5.0
4.5
4.0
3.5
3.0
2.5
2.0
1.5
1.0
0.5
f1 (ppm)
1400
1300
1200
1100
1000
900
800
700
600
500
400
300
200
100
0
-100

VNI-5162

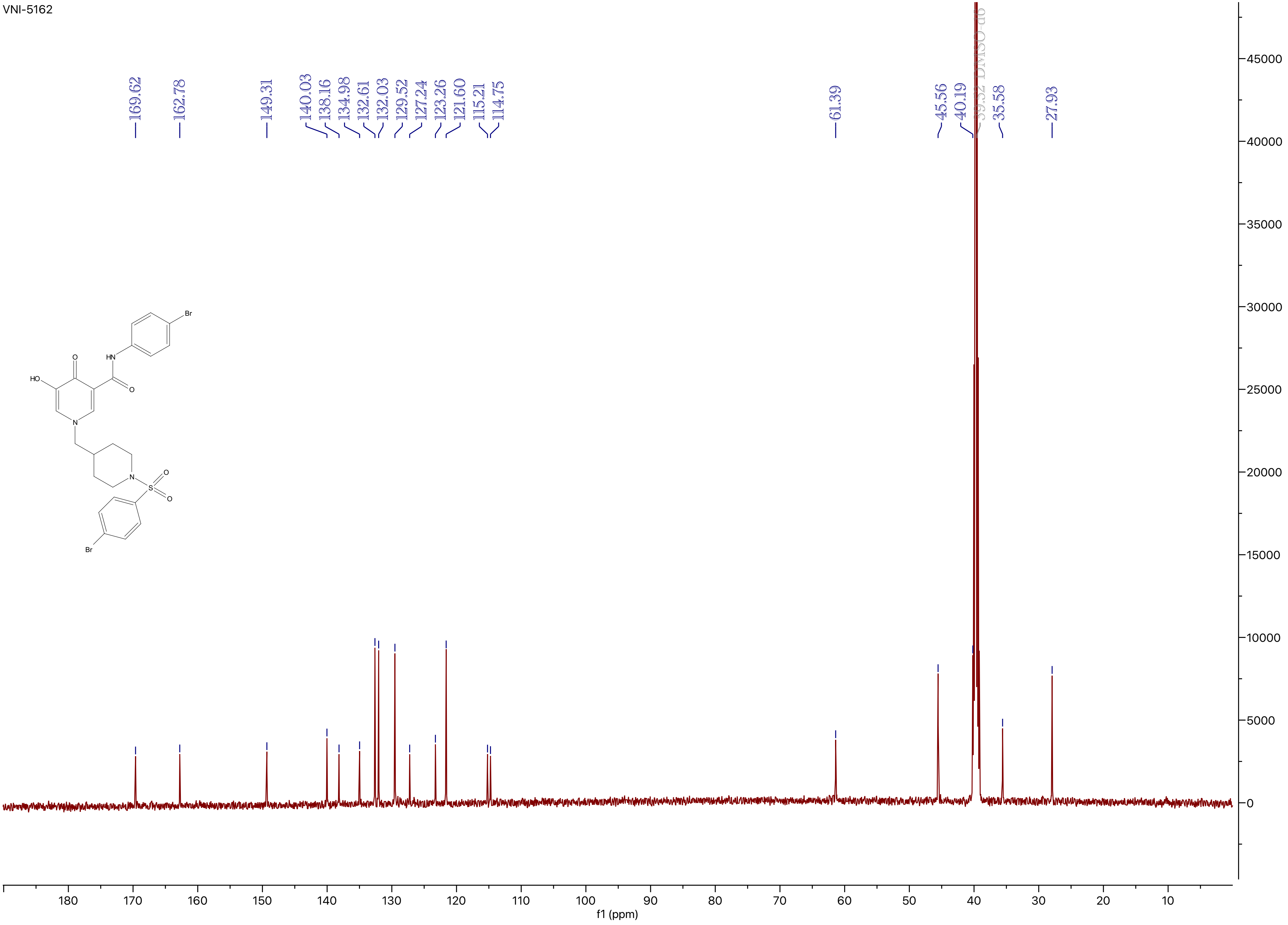
169.62
162.78
149.31
140.03
138.16
134.98
132.61
132.03
129.52
127.24
123.26
121.60
115.21
114.75
61.39
45.56
40.19
39.52 DMSO-d6
35.58
27.93
f1 (ppm)
HO
O
HN
Br
O
N
N
S
O
O
Br

VNI-5163

$^1$H NMR (500 MHz, DMSO-$d_6$) δ 2.410 (s, 1H).

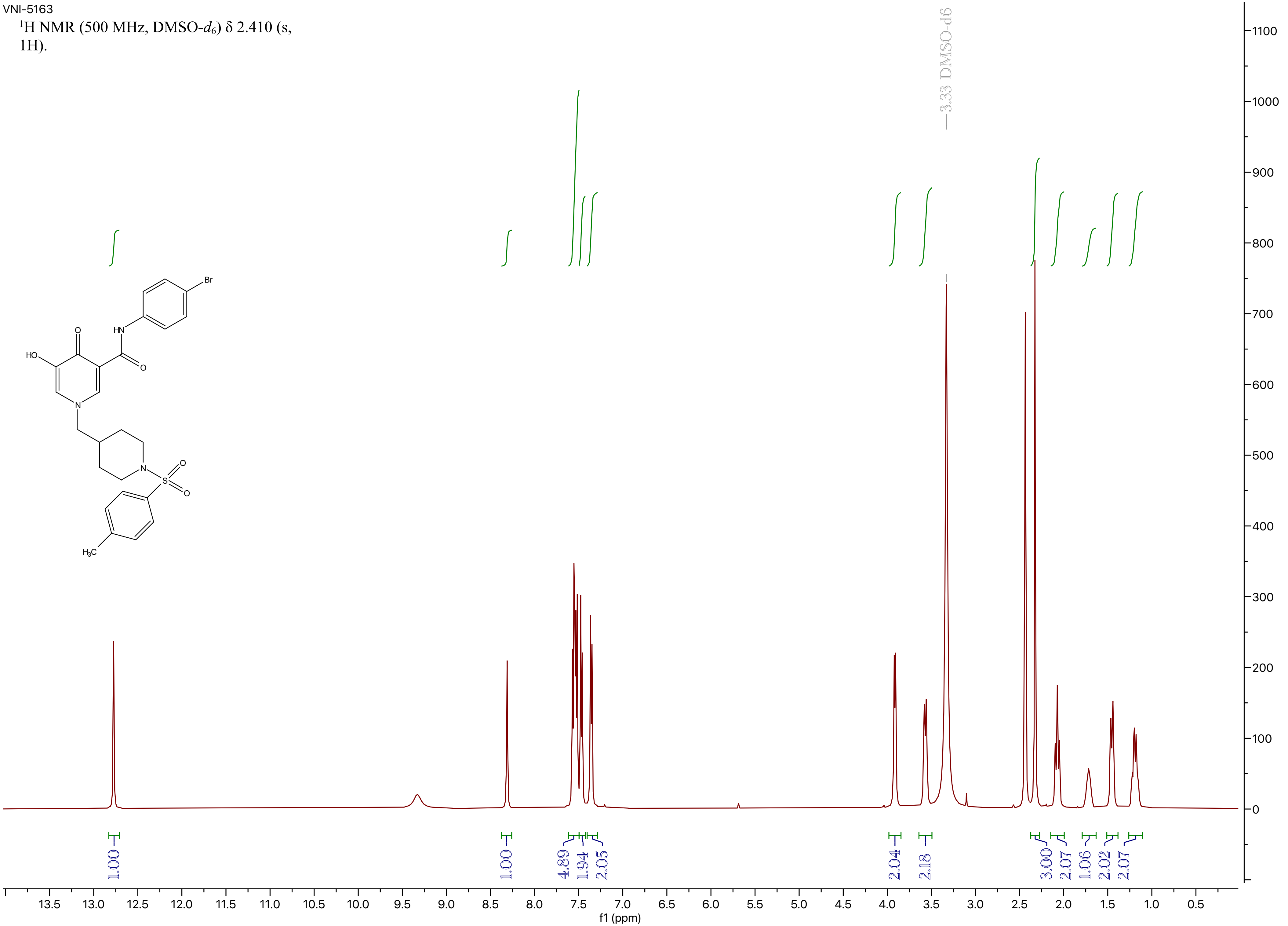

VNI-5163

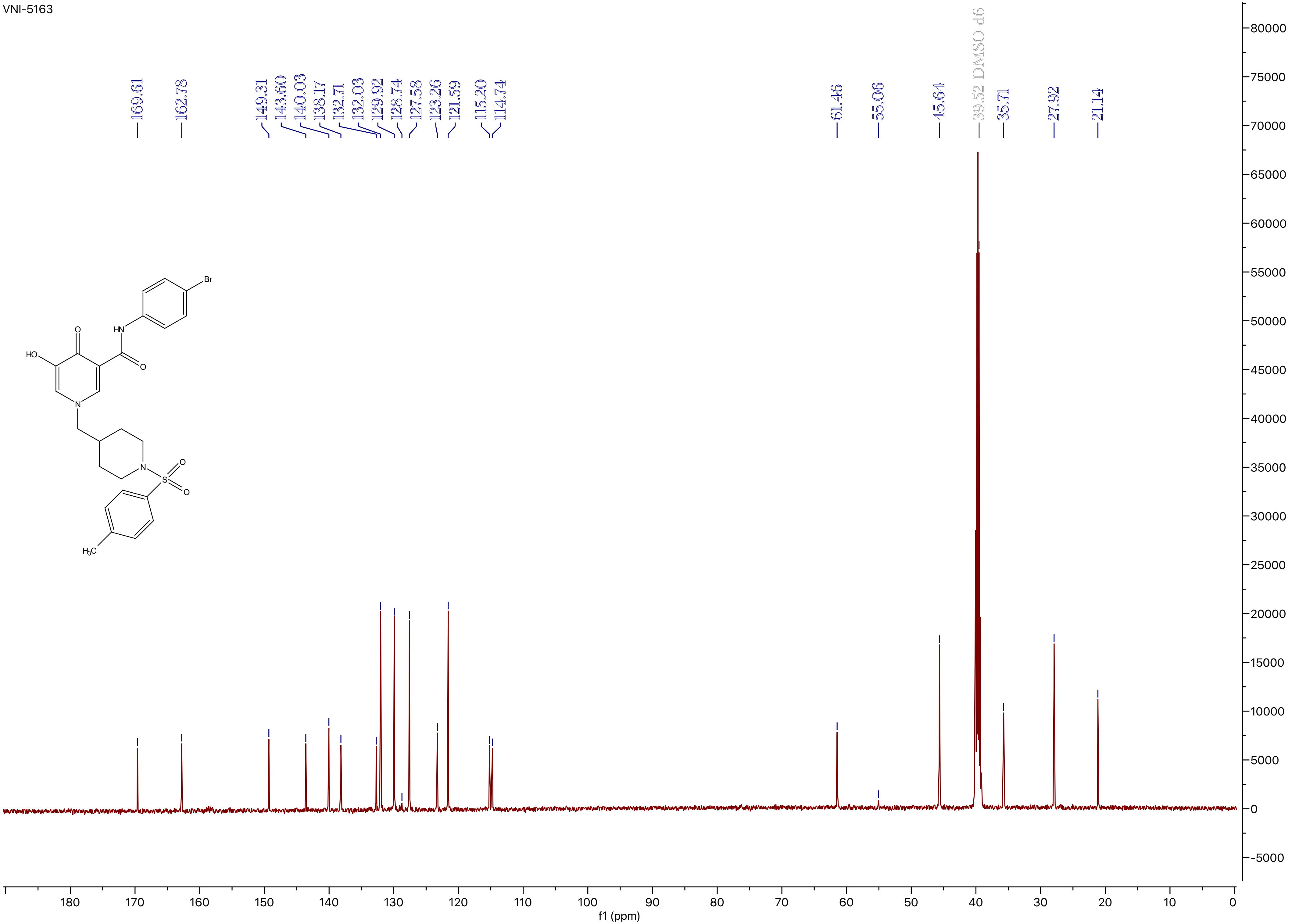

169.61
162.78
149.31
143.60
140.03
138.17
132.71
132.03
129.92
128.74
127.58
123.26
121.59
115.20
114.74
61.46
55.06
45.64
39.52 DMSO-d6
35.71
27.92
21.14
f1 (ppm)
Br
HN
O
HO
O
N
N
S
O
O
H3C

VNI-5164

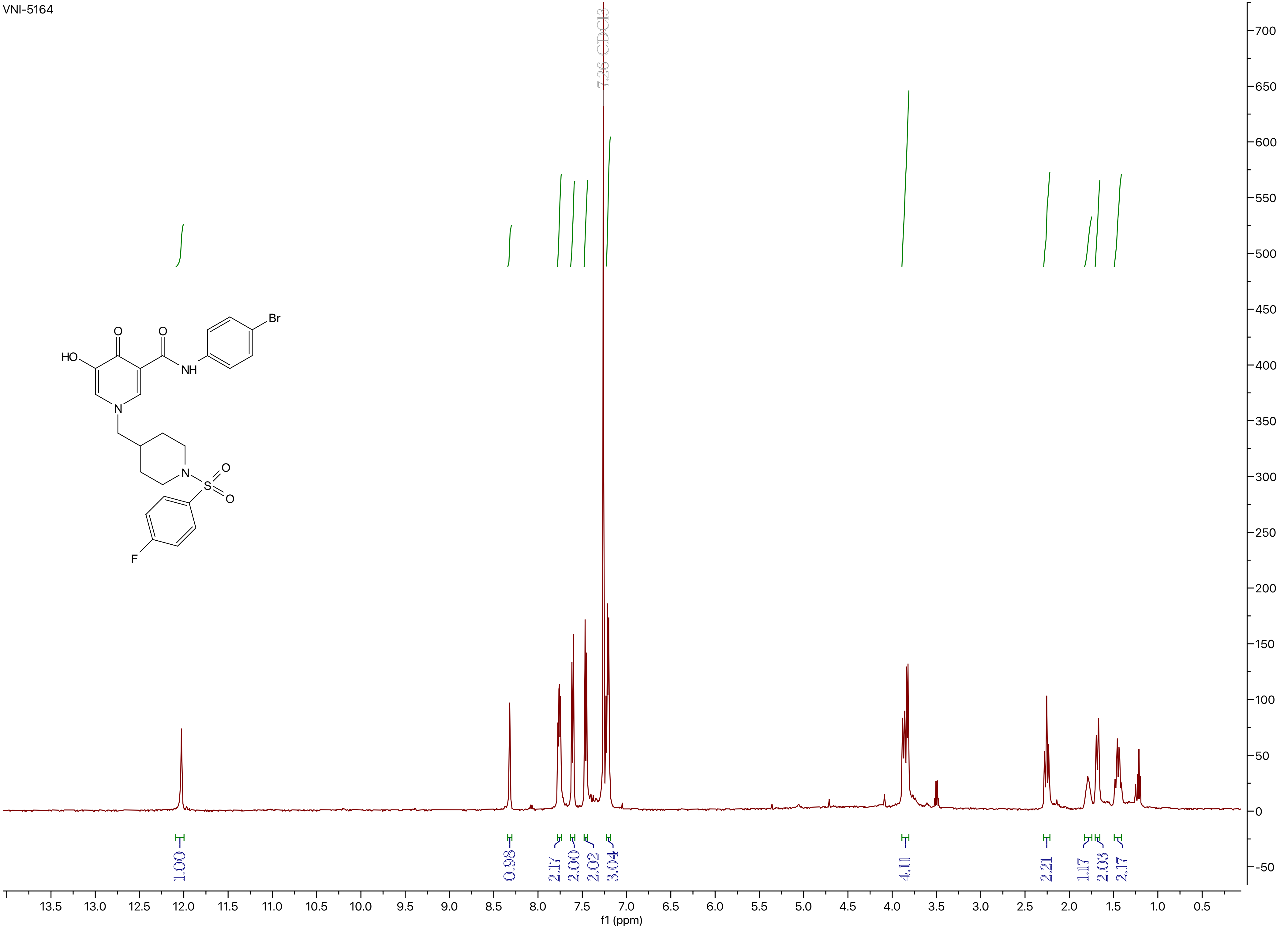
7.26 CDCl3
HO
O
O
NH
Br
N
N
S
O
O
F
1.00
0.98
2.17
2.00
2.02
3.04
4.11
2.21
1.17
2.03
2.17
f1 (ppm)

VNI-5164

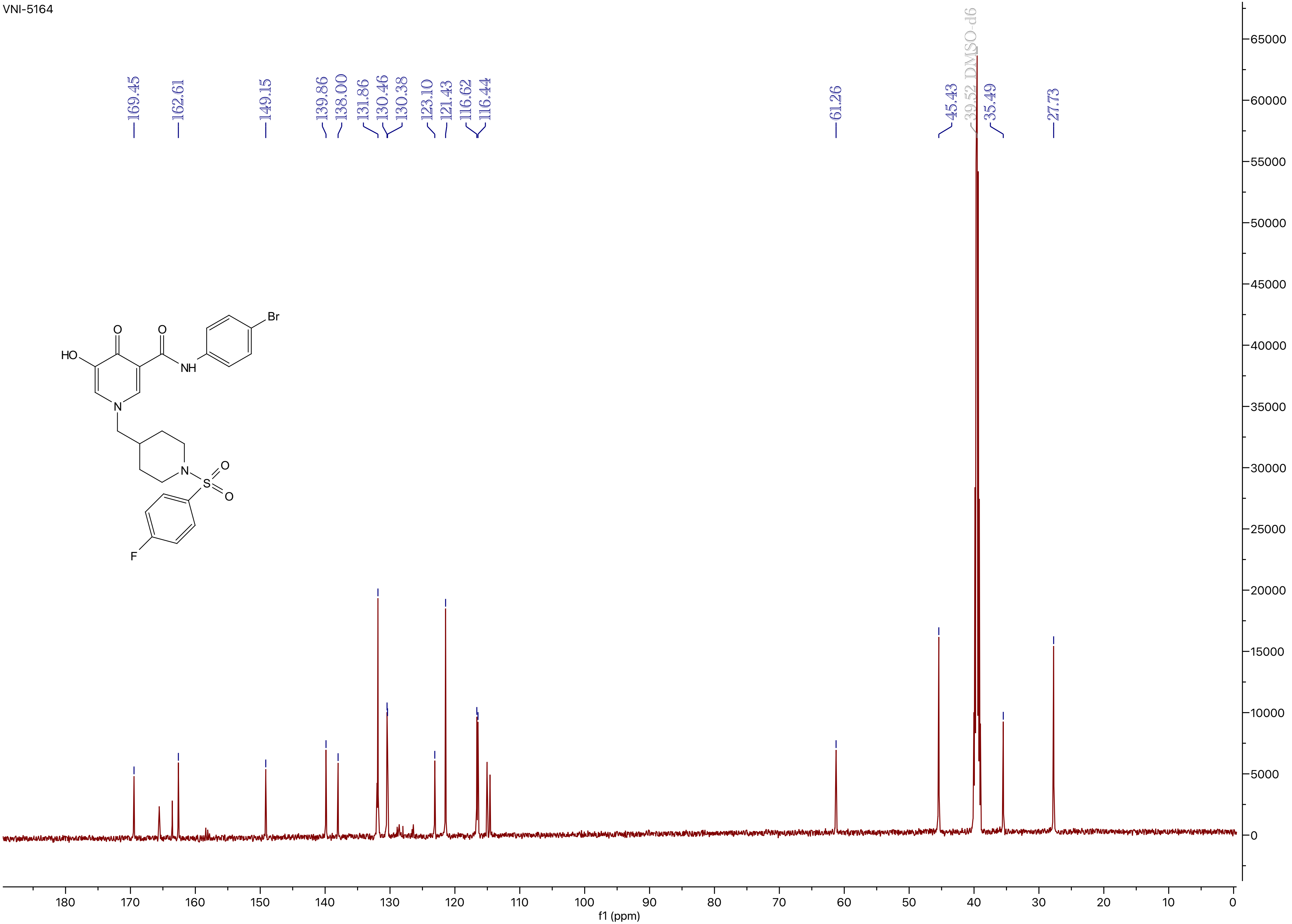
169.45
162.61
149.15
139.86
138.00
131.86
130.46
130.38
123.10
121.43
116.62
116.44
61.26
45.43
39.52 DMSO-d6
35.49
27.73
HO
O
O
NH
Br
N
N
S
O
O
F
65000
60000
55000
50000
45000
40000
35000
30000
25000
20000
15000
10000
5000
0
180
170
160
150
140
130
120
110
100
90
80
70
60
50
40
30
20
10
0
f1 (ppm)

VNI-5165

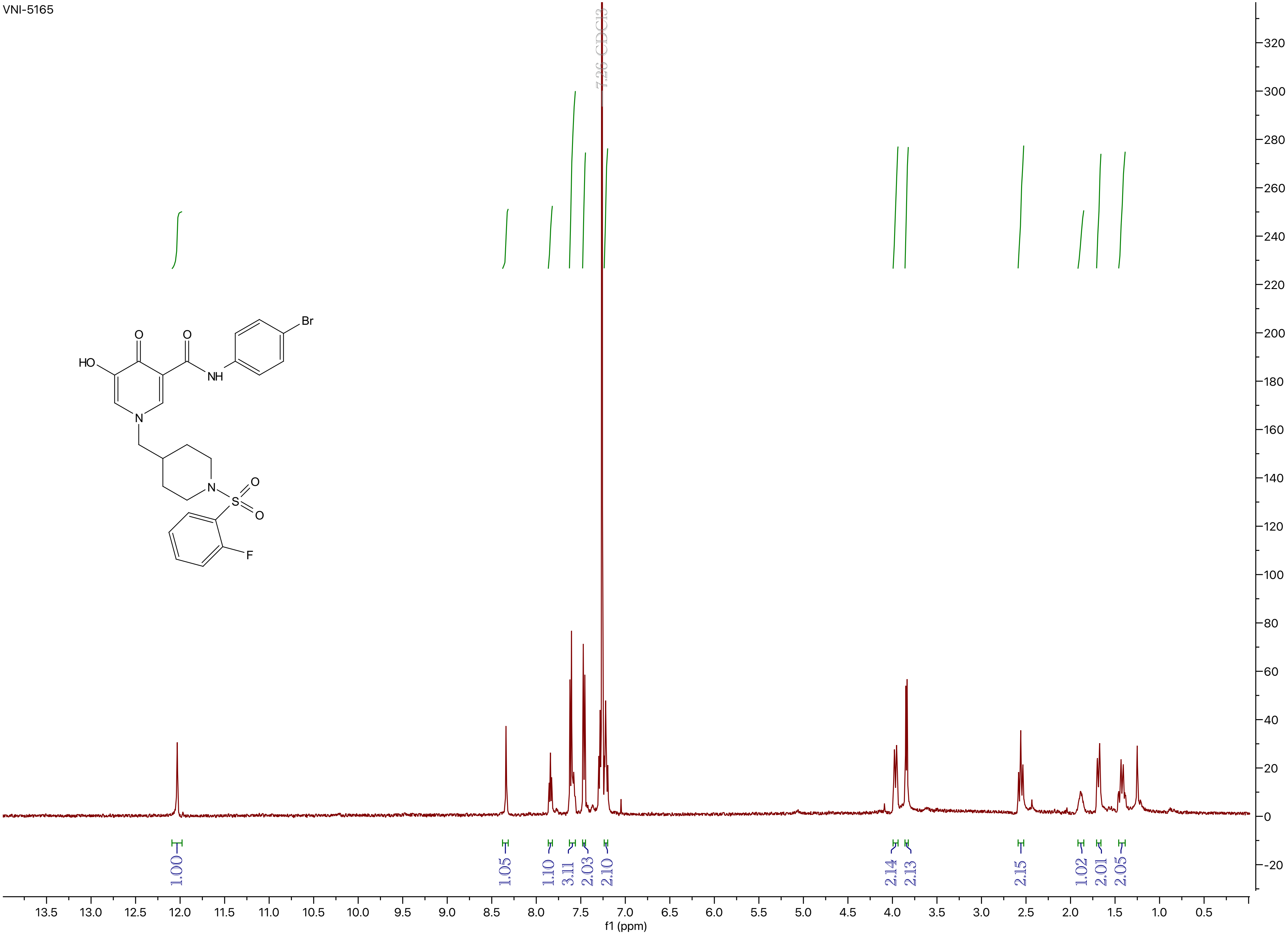
7.26 CDCl3
1.00
1.05
1.10
3.11
2.03
2.10
2.14
2.13
2.15
1.02
2.01
2.05
f1 (ppm)
HO
O
O
NH
Br
N
N
S
O
O
F

VNI-5165

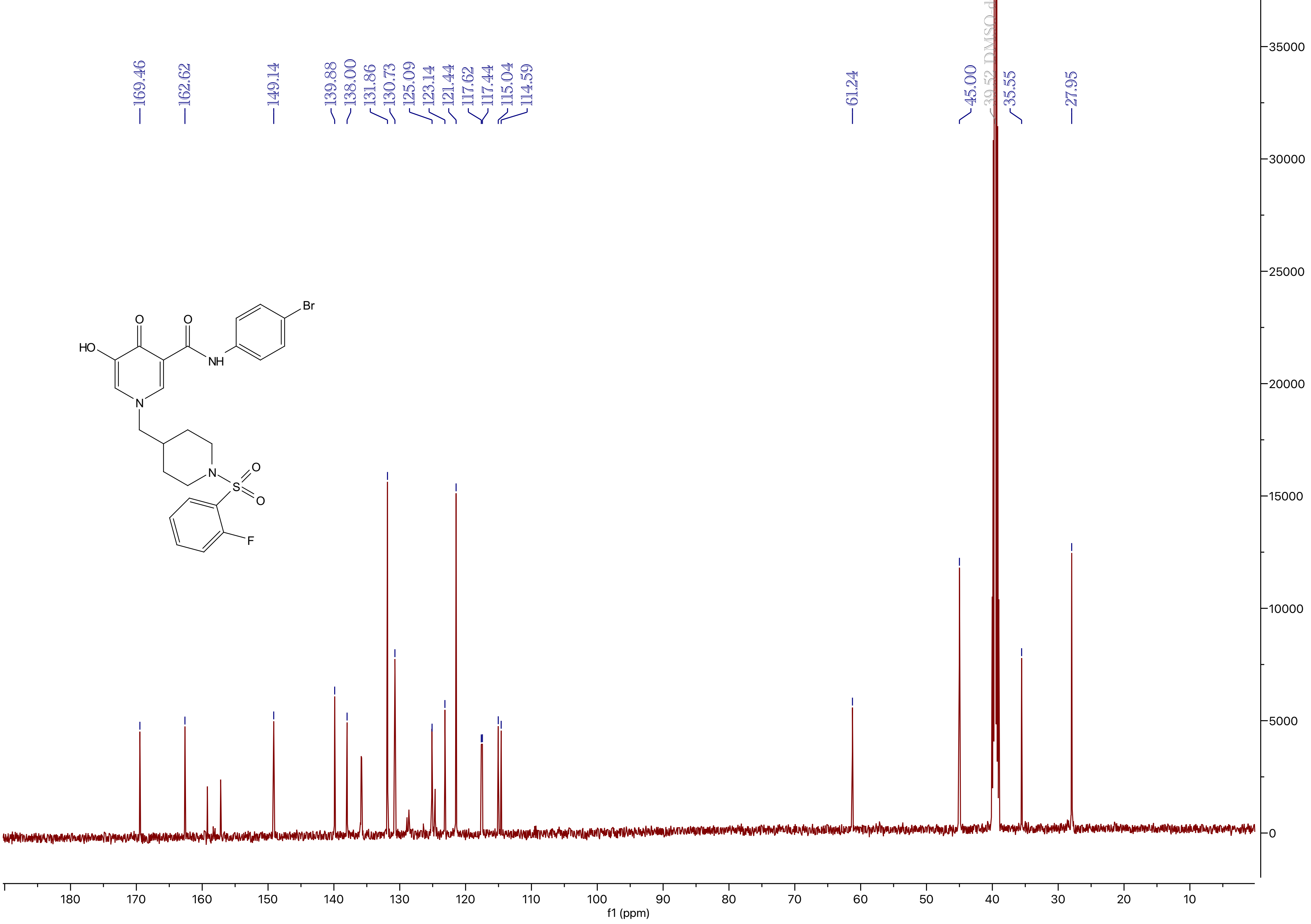
169.46
162.62
149.14
139.88
138.00
131.86
130.73
125.09
123.14
121.44
117.62
117.44
115.04
114.59
61.24
45.00
39.52 DMSO-d6
35.55
27.95
HO
O
O
NH
Br
N
N
S
O
O
F
35000
30000
25000
20000
15000
10000
5000
0
180
170
160
150
140
130
120
110
100
90
80
70
60
50
40
30
20
10
f1 (ppm)

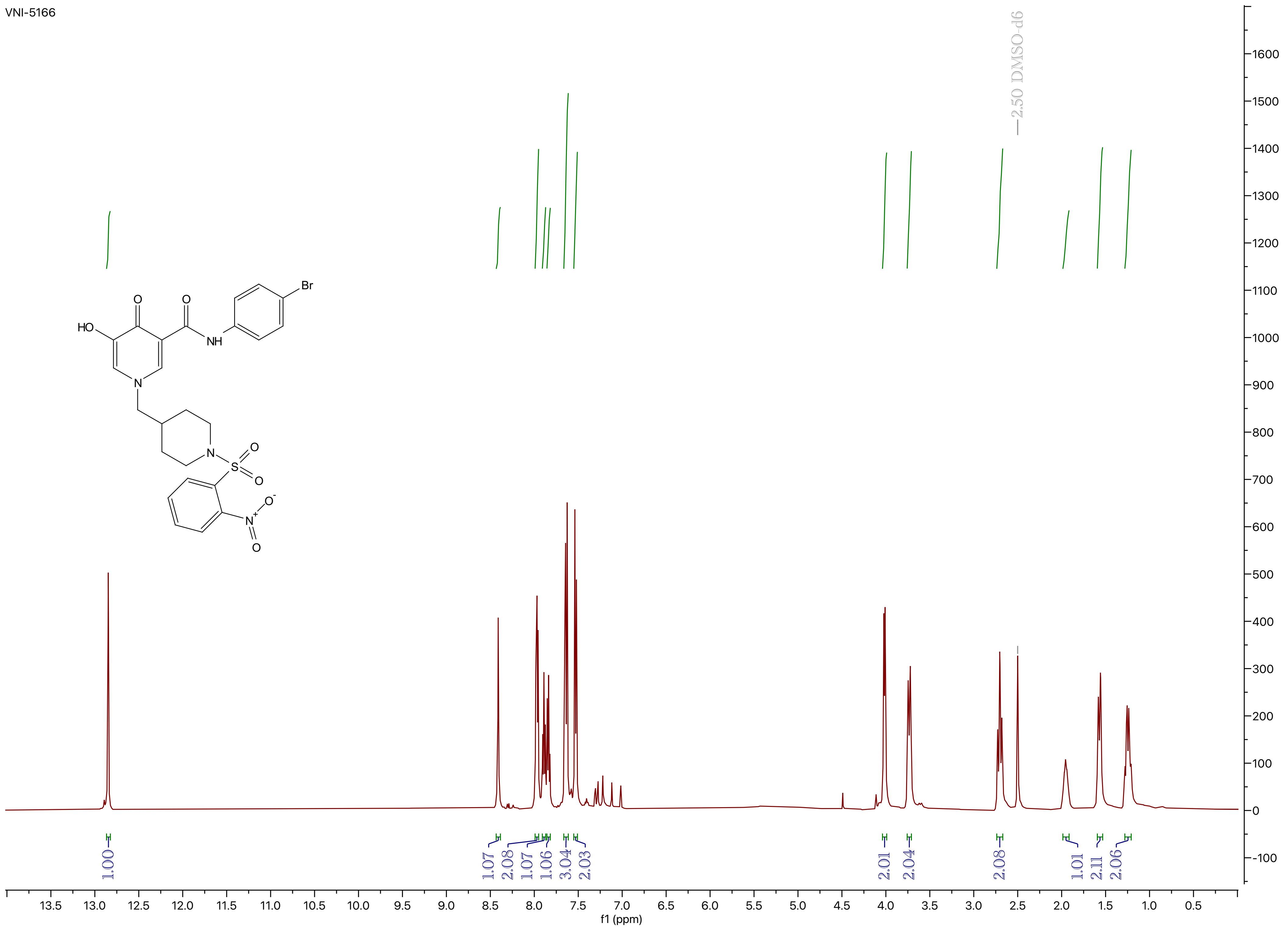
VNI-5166
2.50 DMSO-d6
1.00
1.07
2.08
1.07
1.06
3.04
2.03
2.01
2.04
2.08
1.01
2.11
2.06
f1 (ppm)
HO
O
O
NH
Br
N
N
S
O
O
N+
O-
O

VNI-5166

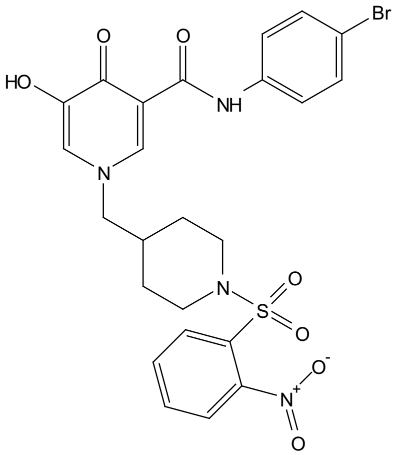

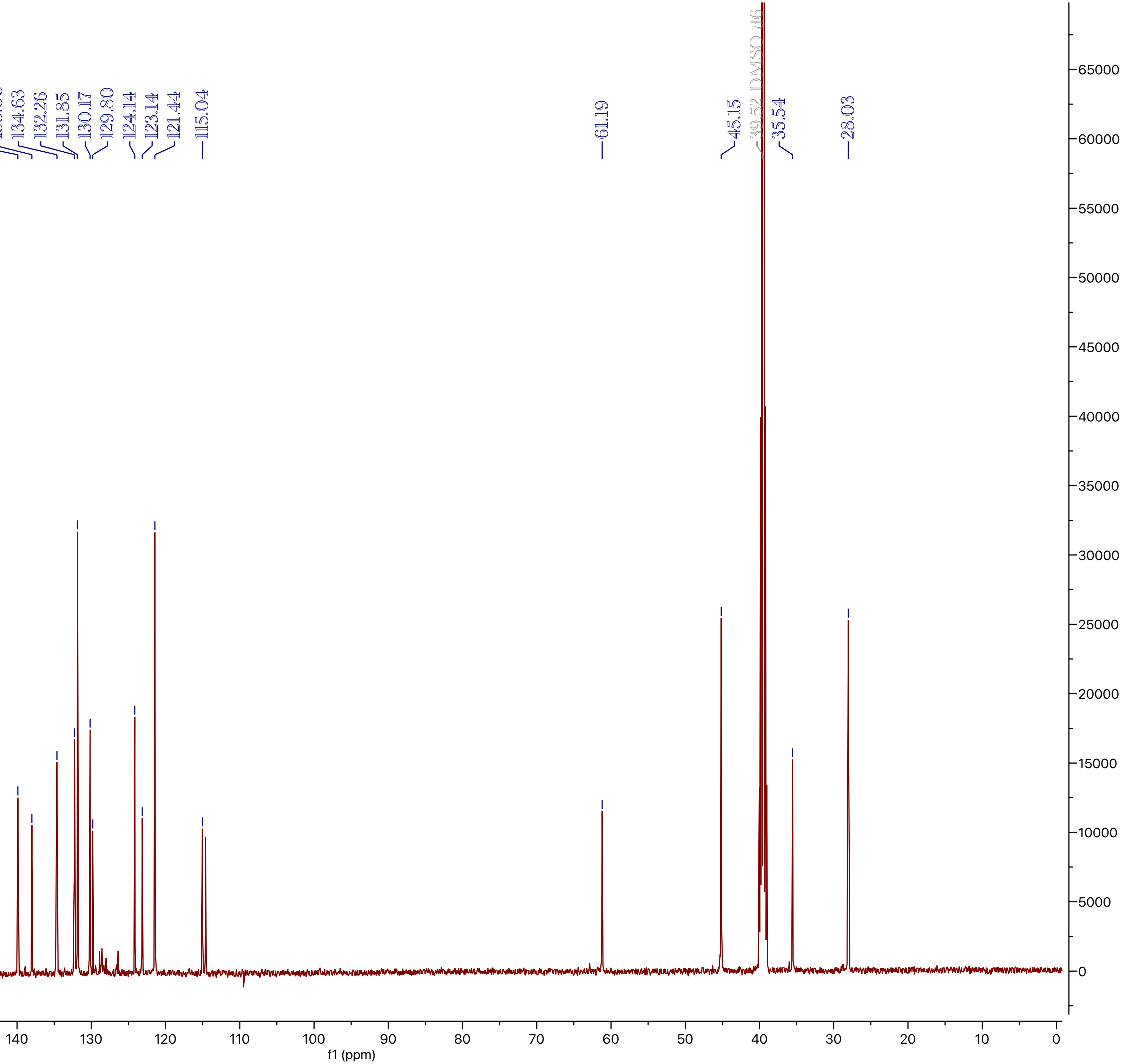
169.47
162.62
149.16
147.78
139.89
138.00
134.63
132.26
131.85
130.17
129.80
124.14
123.14
121.44
115.04
61.19
45.15
39.52 DMSO-d6
35.54
28.03
f1 (ppm)

VNI-5167

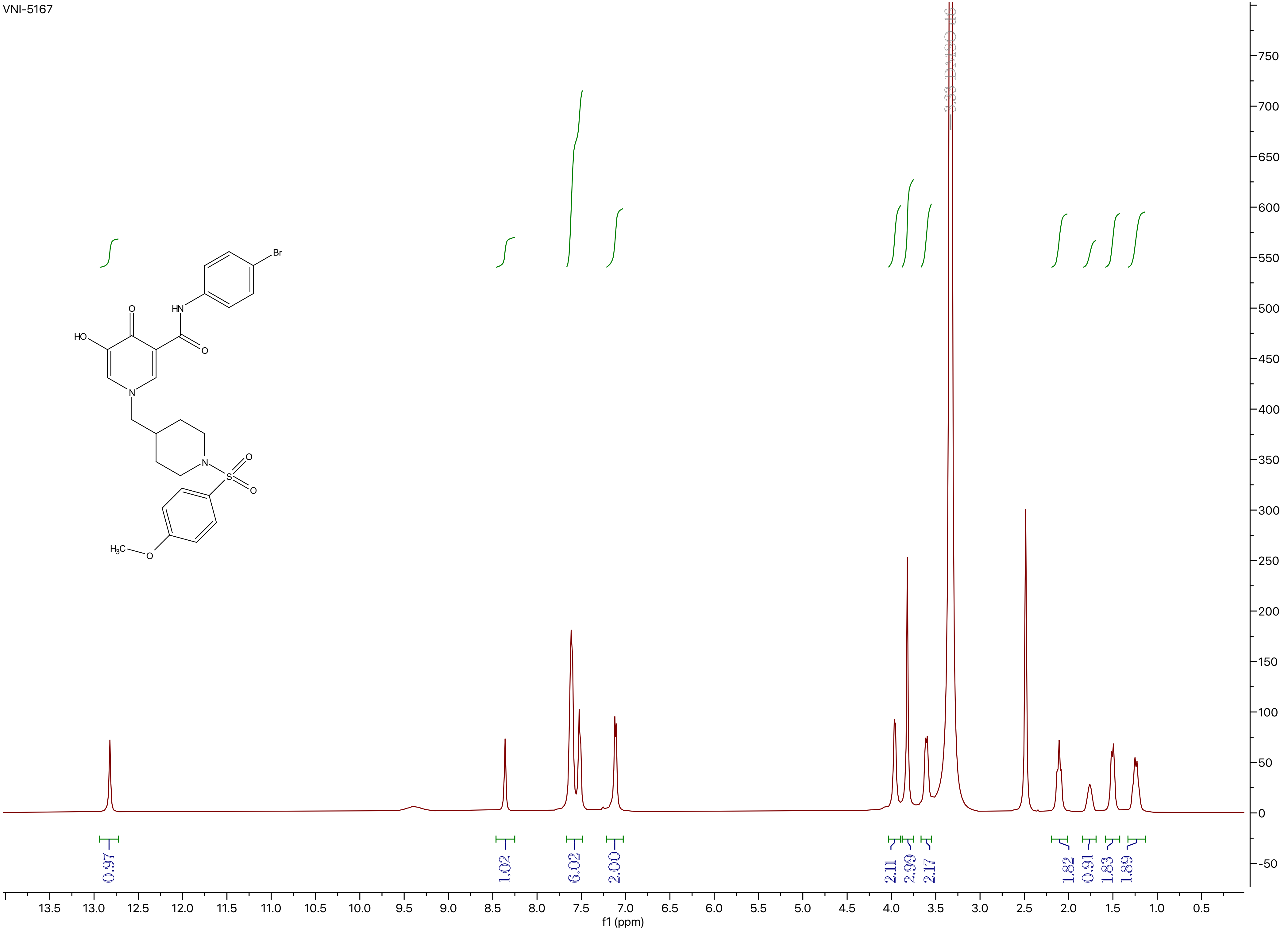

3.33 DMSO-d6
0.97
1.02
6.02
2.00
2.11
2.99
2.17
1.82
0.91
1.83
1.89
f1 (ppm)
HO
O
HN
Br
O
N
N
S
O
O
H3C
O

VNI-5167

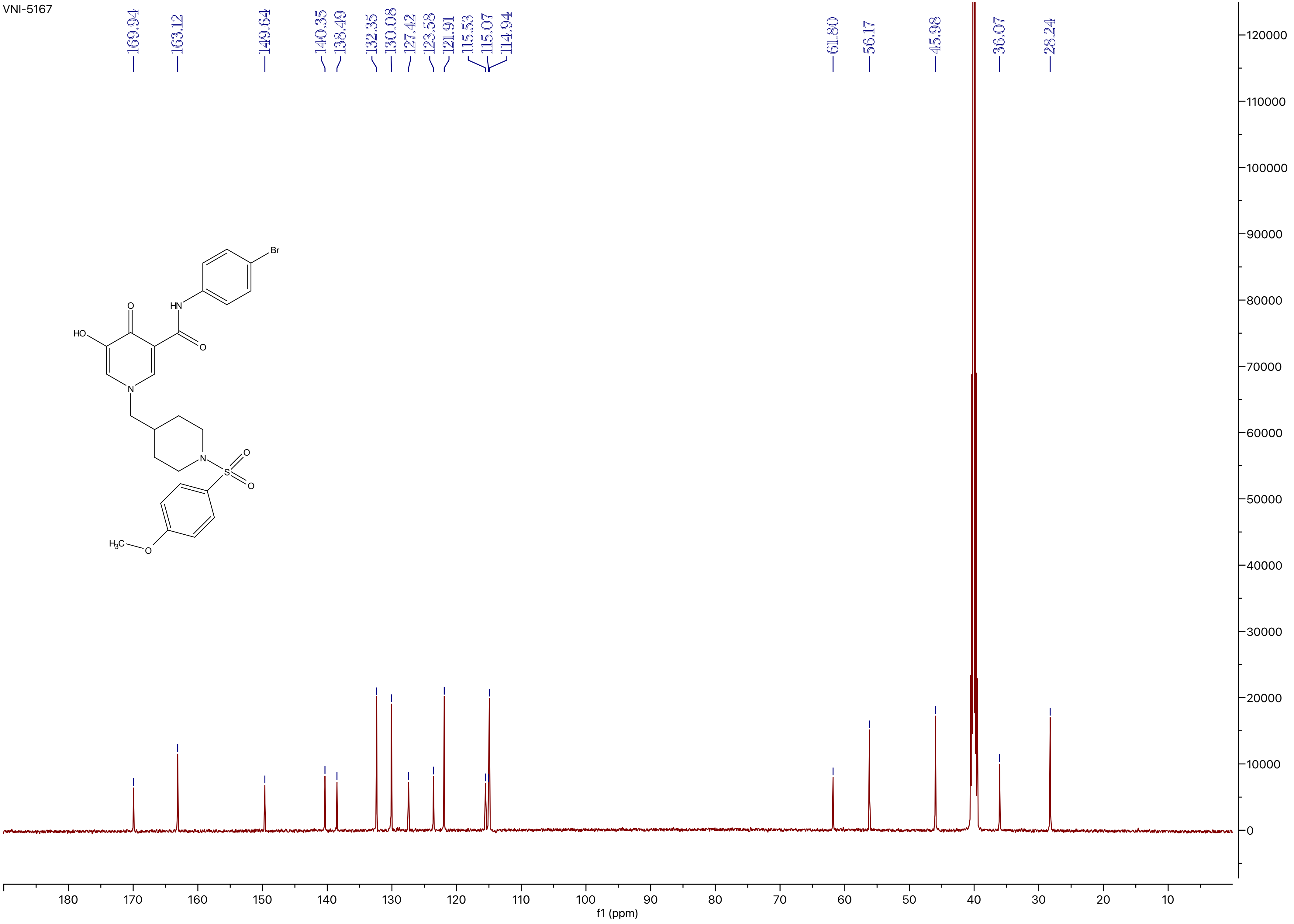
169.94
163.12
149.64
140.35
138.49
132.35
130.08
127.42
123.58
121.91
115.53
115.07
114.94
61.80
56.17
45.98
36.07
28.24
Br
HN
O
HO
O
N
N
S
O
O
$H_3C$
O
120000
110000
100000
90000
80000
70000
60000
50000
40000
30000
20000
10000
0
180
170
160
150
140
130
120
110
100
90
80
70
60
50
40
30
20
10
f1 (ppm)

VNI-5168

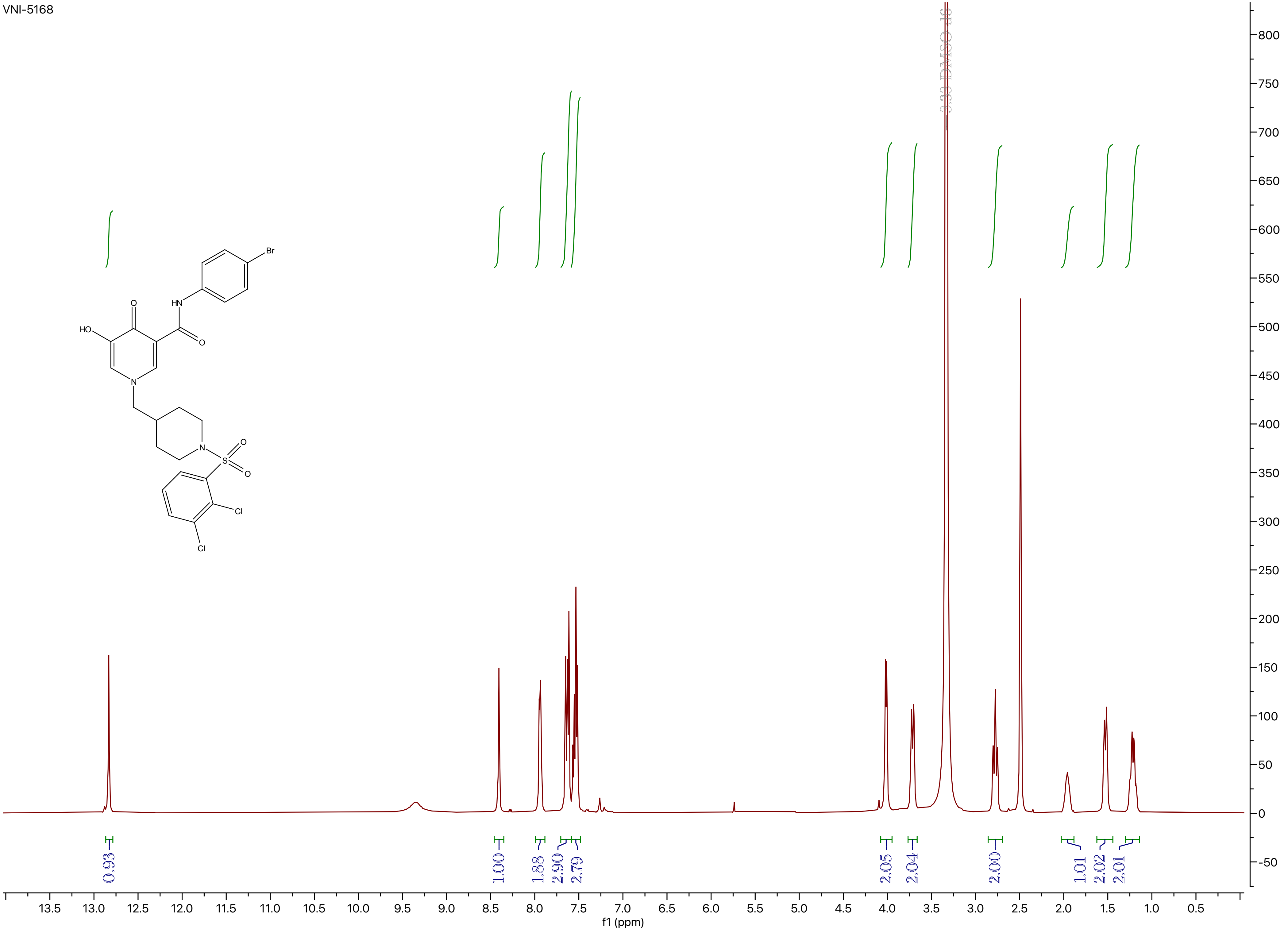

3.33 DMSO d6
0.93
1.00
1.88
2.90
2.79
2.05
2.04
2.00
1.01
2.02
2.01
f1 (ppm)
HO
O
HN
Br
N
S
Cl

VNI-5168

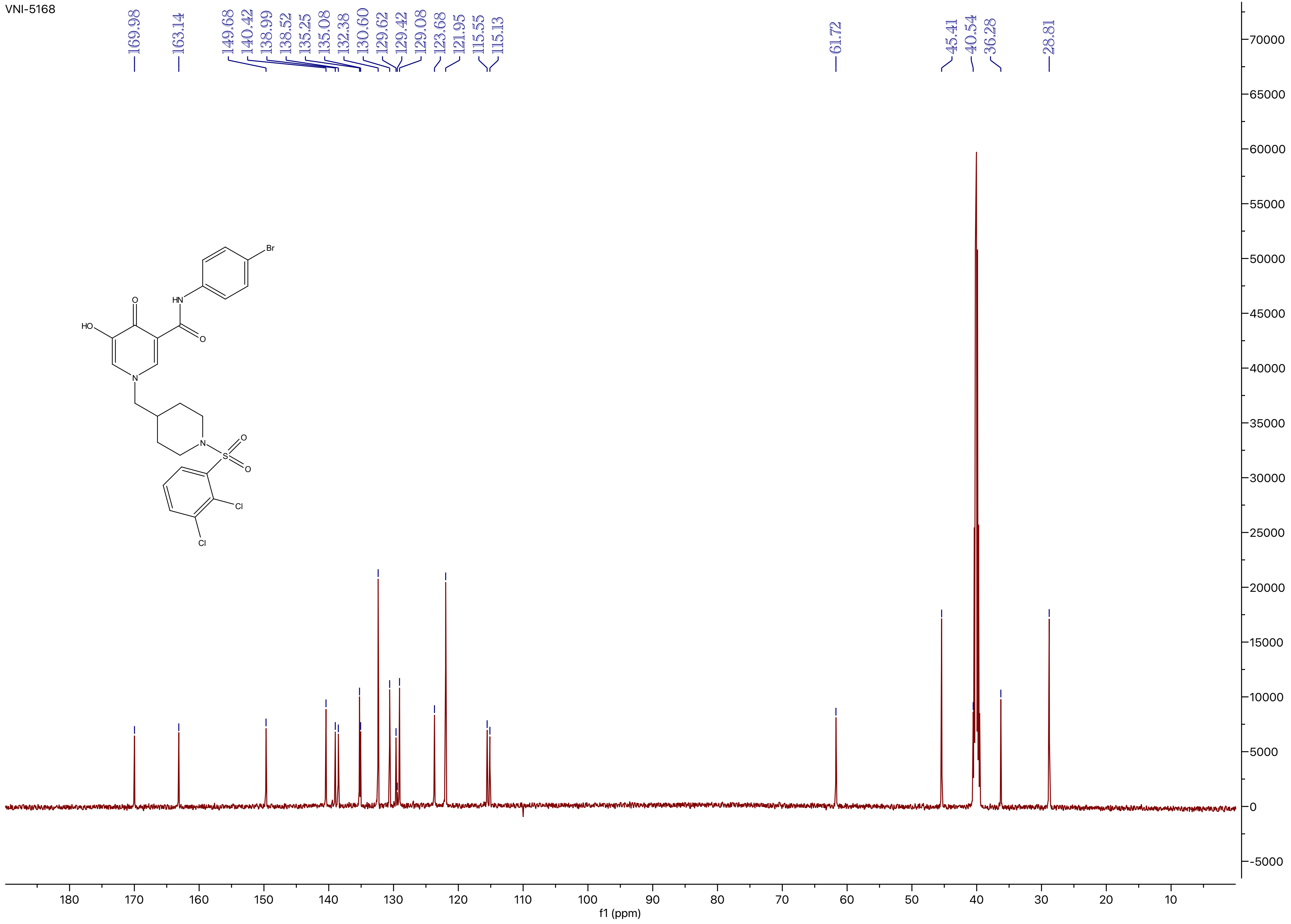
169.98
163.14
149.68
140.42
138.99
138.52
135.25
135.08
132.38
130.60
129.62
129.42
129.08
123.68
121.95
115.55
115.13
61.72
45.41
40.54
36.28
28.81
Br
HN
O
HO
O
N
N
S
O
O
Cl
Cl
70000
65000
60000
55000
50000
45000
40000
35000
30000
25000
20000
15000
10000
5000
0
-5000
180
170
160
150
140
130
120
110
100
90
80
70
60
50
40
30
20
10
f1 (ppm)

VNI-5170

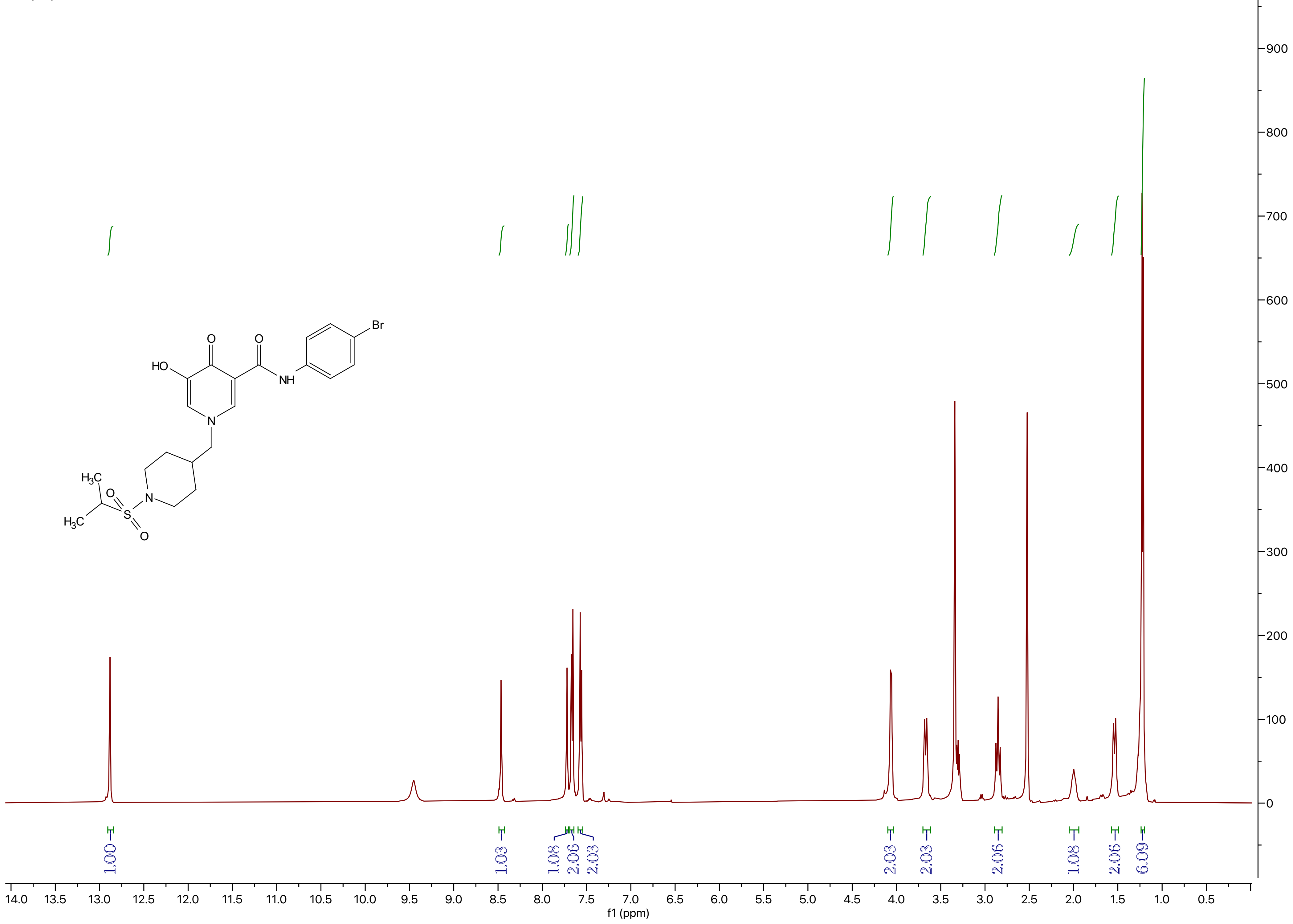

Br
O
O
HO
NH
N
H3C
O
N
S
H3C
O
1.00
1.03
1.08
2.06
2.03
2.03
2.03
2.06
1.08
2.06
6.09
14.0 13.5 13.0 12.5 12.0 11.5 11.0 10.5 10.0 9.5 9.0 8.5 8.0 7.5 7.0 6.5 6.0 5.5 5.0 4.5 4.0 3.5 3.0 2.5 2.0 1.5 1.0 0.5
f1 (ppm)
900
800
700
600
500
400
300
200
100
0

VNI-5170

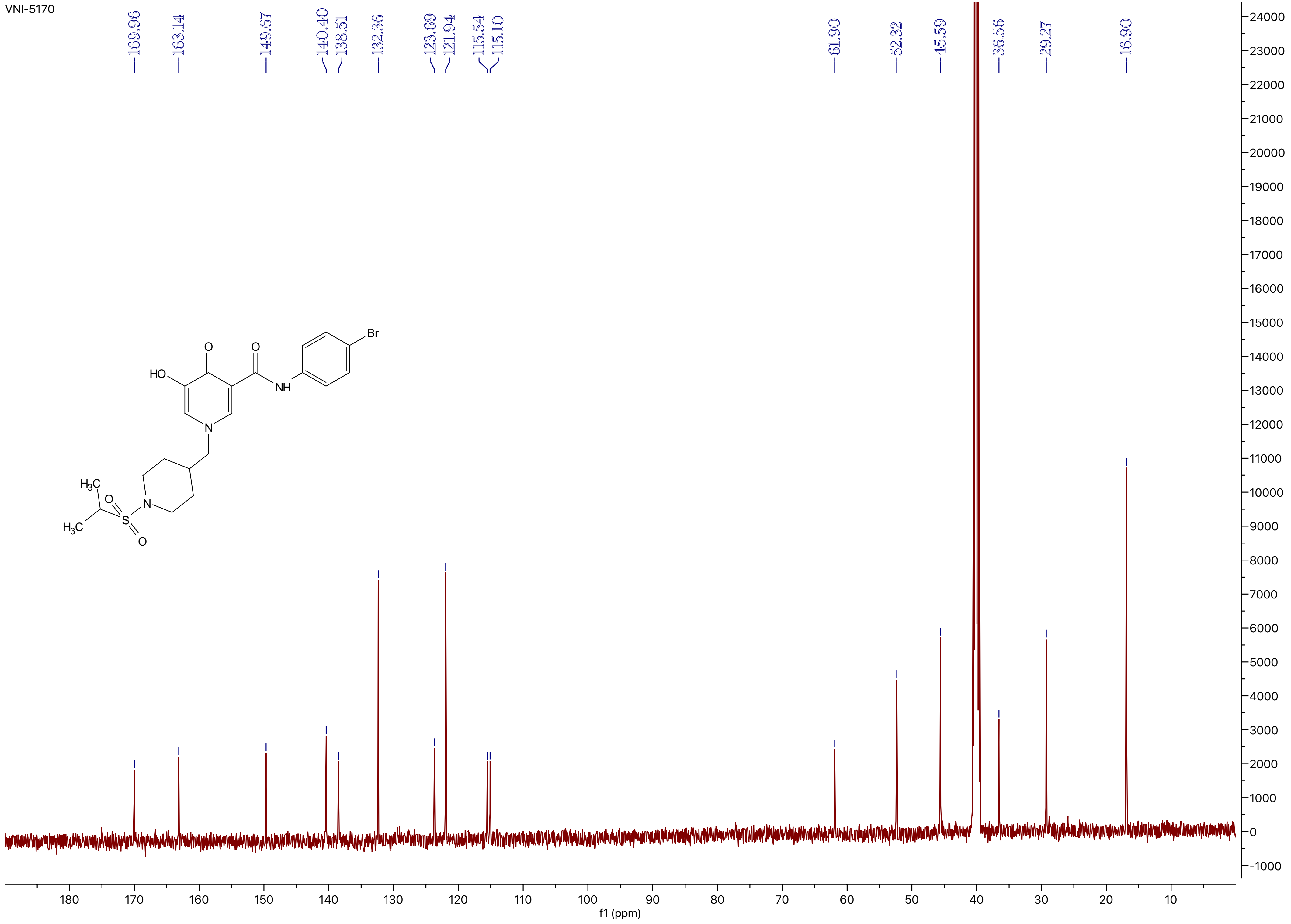
169.96
163.14
149.67
140.40
138.51
132.36
123.69
121.94
115.54
115.10
61.90
52.32
45.59
36.56
29.27
16.90
Br
O
O
HO
NH
N
$H_3C$
O
S
N
$H_3C$
O
f1 (ppm)

VNI-5171

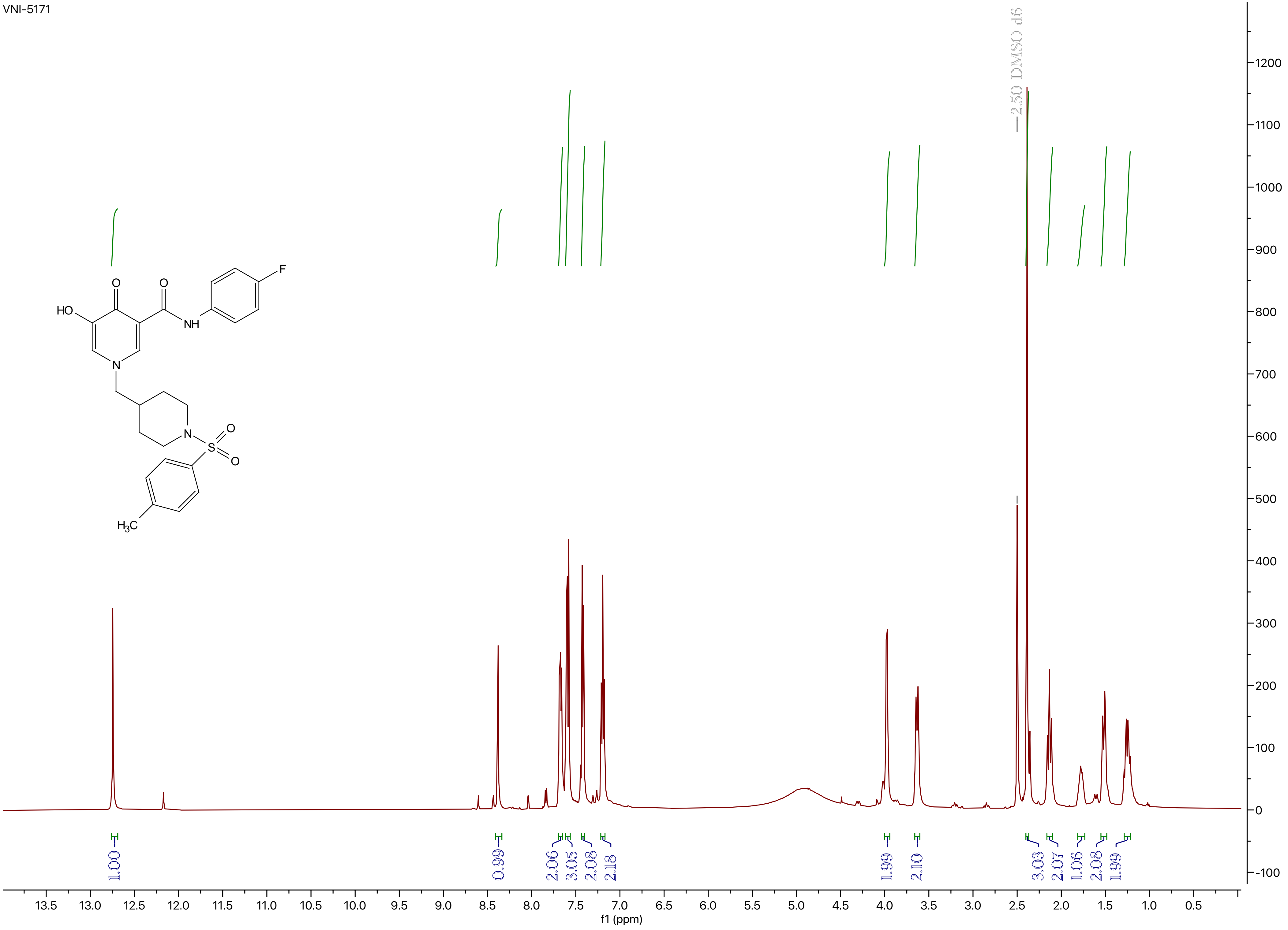
2.50 DMSO-d6
HO
O
O
NH
F
N
N
S
O
O
H3C
1.00
0.99
2.06
3.05
2.08
2.18
1.99
2.10
3.03
2.07
1.06
2.08
1.99
13.5 13.0 12.5 12.0 11.5 11.0 10.5 10.0 9.5 9.0 8.5 8.0 7.5 7.0 6.5 6.0 5.5 5.0 4.5 4.0 3.5 3.0 2.5 2.0 1.5 1.0 0.5
f1 (ppm)
1200 1100 1000 900 800 700 600 500 400 300 200 100 0 -100

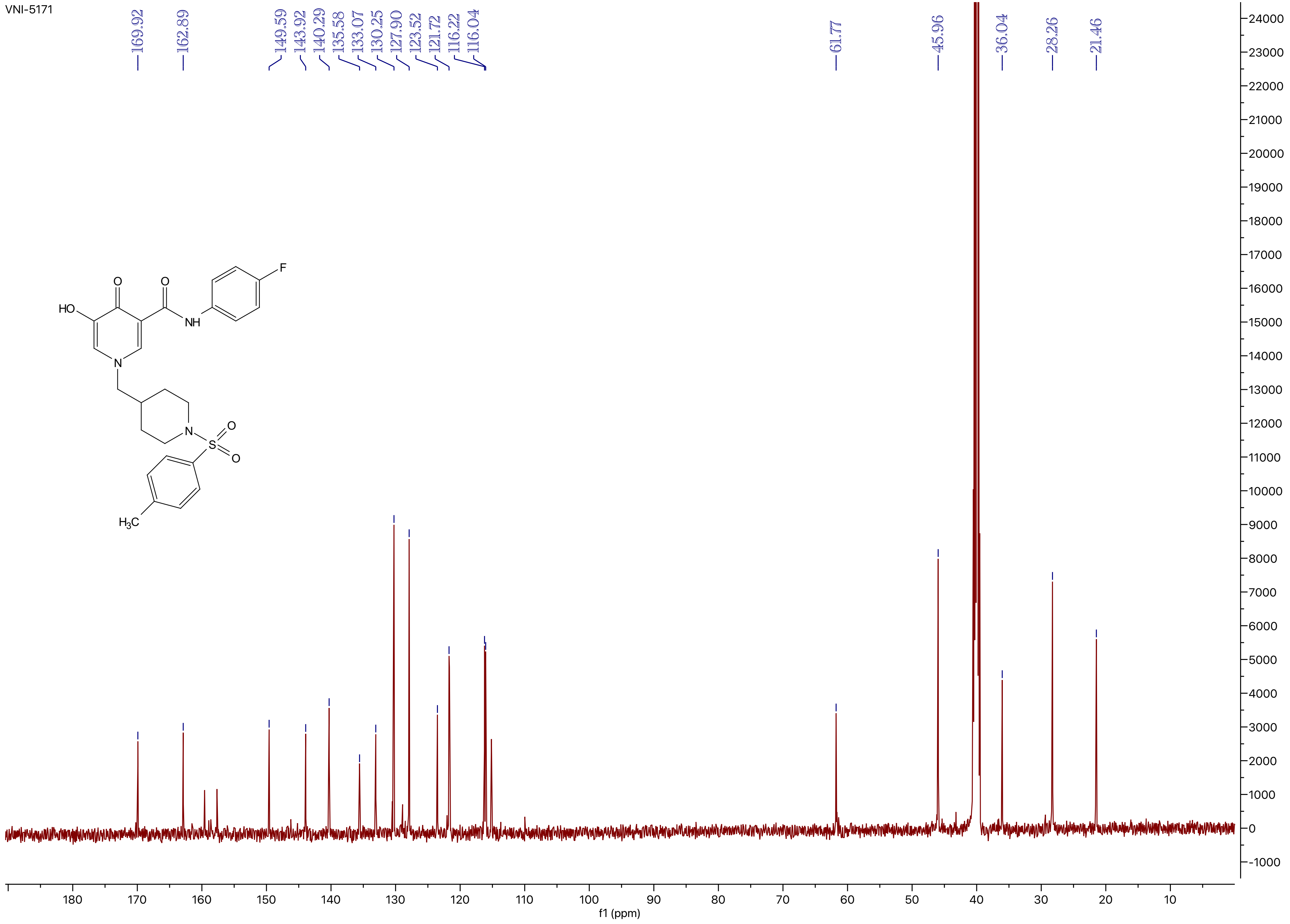
VNI-5171
169.92
162.89
149.59
143.92
140.29
135.58
133.07
130.25
127.90
123.52
121.72
116.22
116.04
61.77
45.96
36.04
28.26
21.46
HO
O
O
NH
F
N
N
S
O
O
H3C
f1 (ppm)

VNI-5172

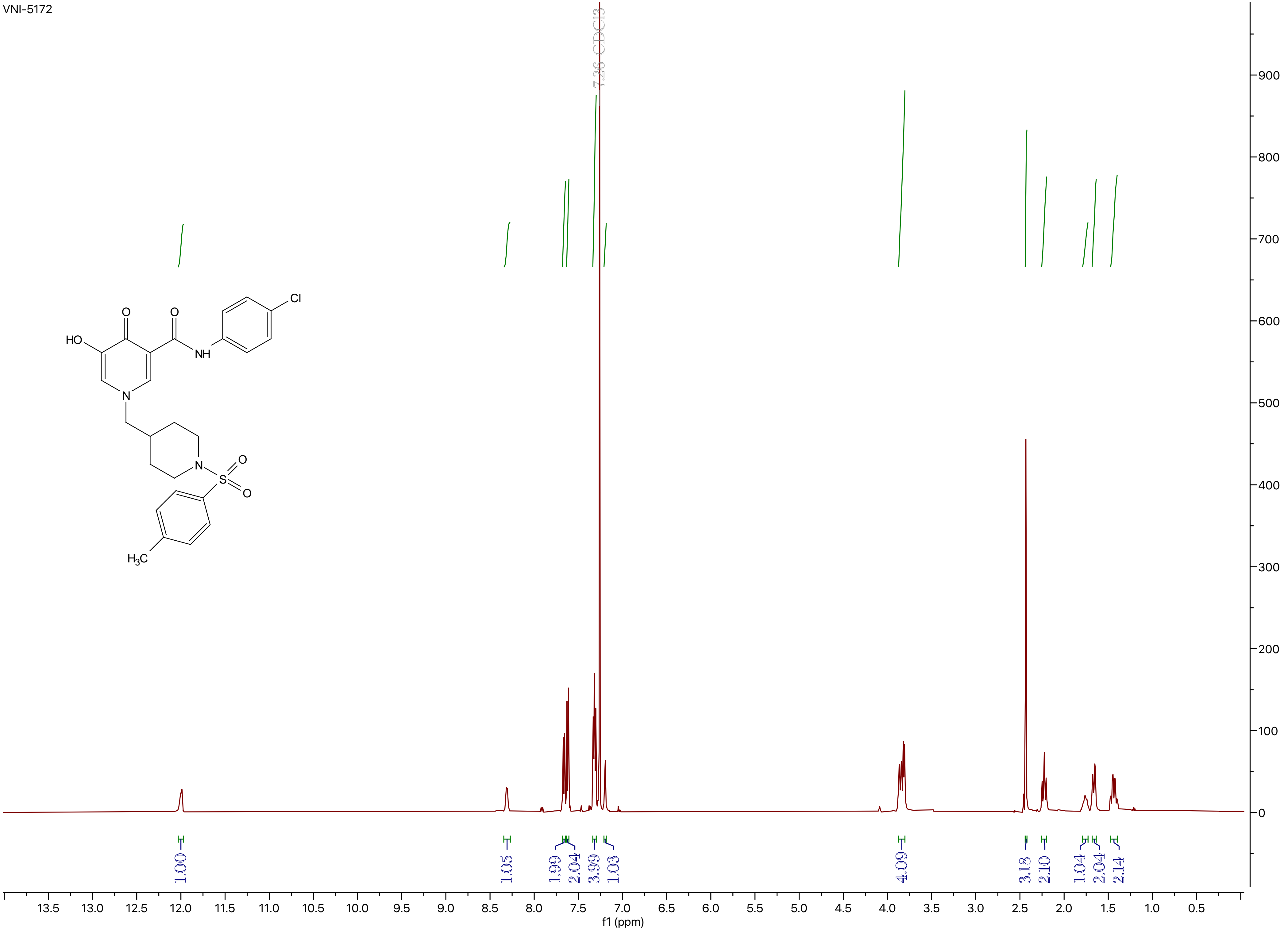
7.26 CDCl3
HO
O
O
NH
Cl
N
N
S
O
O
H3C
1.00
1.05
1.99
2.04
3.99
1.03
4.09
3.18
2.10
1.04
2.04
2.14
13.5
13.0
12.5
12.0
11.5
11.0
10.5
10.0
9.5
9.0
8.5
8.0
7.5
7.0
6.5
6.0
5.5
5.0
4.5
4.0
3.5
3.0
2.5
2.0
1.5
1.0
0.5
f1 (ppm)
900
800
700
600
500
400
300
200
100
0

VNI-5172

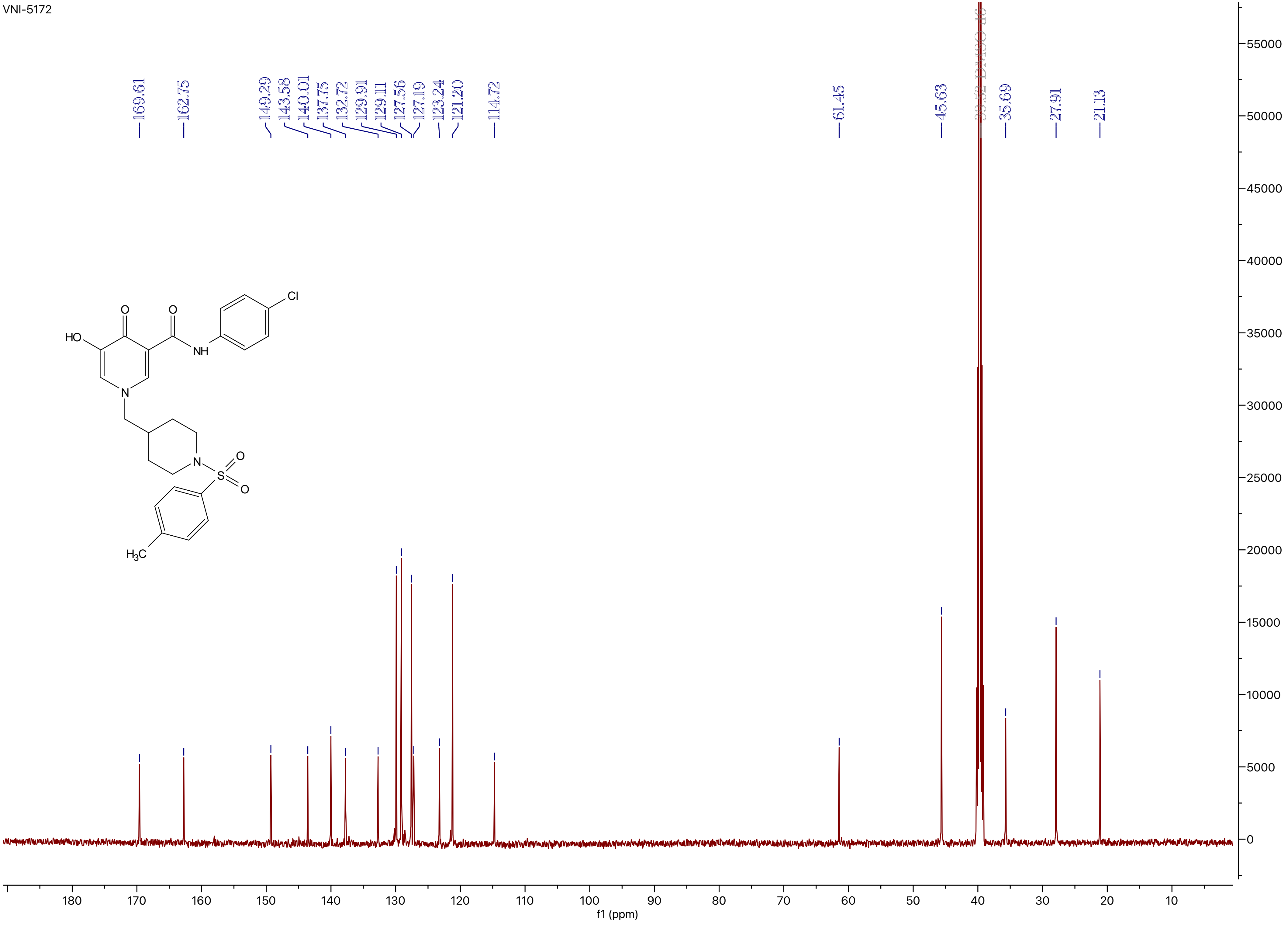

169.61
162.75
149.29
143.58
140.01
137.75
132.72
129.91
129.11
127.56
127.19
123.24
121.20
114.72
61.45
45.63
39.52 DMSO d6
35.69
27.91
21.13
55000
50000
45000
40000
35000
30000
25000
20000
15000
10000
5000
0
180
170
160
150
140
130
120
110
100
90
80
70
60
50
40
30
20
10
f1 (ppm)
HO
O
O
NH
Cl
N
N
S
O
O
$H_3C$
